# Removing unwanted variation from large-scale cancer RNA-sequencing data

**DOI:** 10.1101/2021.11.01.466731

**Authors:** Ramyar Molania, Momeneh Foroutan, Johann A. Gagnon-Bartsch, Luke Gandolfo, Gavriel Olshansky, Alexander Dobrovic, Anthony T Papenfuss, Terence P Speed

**Author notes:** These authors contributed equally (Anthony T Papenfuss, Terence P Speed).

## Abstract

The accurate identification and effective removal of unwanted variation are essential to derive meaningful biological results from RNA-seq data, especially when the data come from large and complex studies. We have used The Cancer Genome Atlas (TCGA) RNA-seq data to show that library size, batch effects, and tumor purity are major sources of unwanted variation across all TCGA RNA-seq datasets and that existing gold standard approaches to normalizations fail to remove this unwanted variation. Additionally, we illustrate how different sources of unwanted variation can compromise downstream analyses, including gene co-expression, association between gene expression and survival outcomes, and cancer subtype identifications. Here, we propose the use of a novel strategy, pseudo-replicates of pseudo-samples (PRPS), to deploy the Removing Unwanted Variation III (RUV-III) method to remove different sources of unwanted variation from large and complex gene expression studies. Our approach requires at least one roughly known biologically homogenous subclass of samples shared across sources of unwanted variation. To create PRPS, we first need to identify the sources of unwanted variation, which we will call batches in the data. Then the gene expression measurements of biologically homogeneous sets of samples are averaged within batches, and the results called pseudo-samples. Pseudo-samples with the same biology and different batches are then defined to be pseudo-replicates and used in RUV-III as replicates. The variation between pseudo-samples of a set pseudo-replicates is mainly unwanted variation. We illustrate the value of our approach by comparing it to the TCGA normalizations on several TCGA RNA-seq datasets. RUV-III with PRPS can be used for any large genomics project involving multiple labs, technicians, or platforms.

## Introduction

The effective identification and removal of unwanted variation are essential aspects of gene expression data analysis, particularly when the data come from large and complex studies. This variation can introduce artifactual or obscure true biological signals in the data and lead to inaccurate or misleading biological conclusions [1-5].

Library size differences between samples are a major source of unwanted variation in almost all RNA-seq studies, and these need to be accommodated to make gene expression measurements comparable across samples. Most RNA-seq normalizations adjust for library size differences across samples using global scaling factors based on library size or other statistical features of the raw count data, such as their upper-quartiles [6-8]. These normalizations simply divide the gene counts in each sample by a single scale factor. The implicit assumption underlying such methods is that all the gene-level counts are proportional to the scale factors and that it should be adequate to adjust them for library size in this way across samples. We will show situations where genes counts cannot all be adjusted for library size by a single scale factor, regardless of how it is calculated, presenting a challenge for current RNA-seq normalizations.

Tumor purity, the proportion of cancer cells in solid tumor tissues, is another major source of variation in cancer RNA-seq data. This variation has been viewed as an intrinsic characteristic of tumor samples and linked to several patient clinical outcomes in various cancer types [6-9]. However, variation in tumor purity can compromise downstream analyses in cancer RNA-seq studies [10-12]. Current RNA-seq normalizations and batch correction methods are not designed to remove this variation from the data. Importantly, adjusting counts for tumor purity variation using regression models risks removing important biological signal confounded with purity.

Batch effects are another major source of unwanted variation in large RNA-seq studies, where samples are necessarily processed across a range of experimental conditions. This variation can compromise accurate interpretation of the data and lead to misleading conclusions.

The majority of batch correction methods are based on linear regression. For individual gene expression, they fit a linear model with blocking term for batch variables. Then, the coefficient for each blocking term is set to zero and the corrected expression values are computed from the residuals. The implicit assumption underlying such methods is that the composition of biological populations is identical within each batch. Correcting gene expression counts for batch effects using these method risks removing biological signal confounded with batches.

Further, it has been shown that batch effects influence different sets of genes in different ways [3]. Sample wise normalization, including normalizations that relay on global scaling factors generally fail to remove this source of unwanted variation from the data.

We previously developed a new normalization method, Removing Unwanted Variation-III (RUV-III), that makes essential use of well-designed technical replicates and negative control genes to estimate unwanted variation and remove it from the data [5]. Briefly, RUV-III uses negative control genes and technical replicates to estimate sample-wise and gene-wise normalization parameters. However, the effectiveness of the RUV-III method is challenged in two different scenarios. Firstly, RUV-III cannot be used effectively in situations where no suitable technical replicates are available and well-distributed across the sources of unwanted variation. Secondly, since a sample’s tumor purity will be essentially the same across all of its technical replicates, the original RUV-III is not able to estimate and remove this kind of variation using standard technical replicates.

To overcome these challenges, we here propose a novel approach using of pseudo-replicates of pseudo-samples (PRPS) to deploy RUV-III to remove library size, tumor purity and batch effects from RNA-seq data. To use RUV-III with PRPS, we first need to identify the sources of unwanted variation and major biological populations in the data. Pseudo-samples are *in silico* samples created from small groups of samples that are roughly homogeneous with respect to unwanted variation and biology. The gene expression differences between two or more pseudo-samples with the same biology will largely be unwanted variation. RUV-III makes use of such differences together with negative control genes to estimate and remove unwanted variation from the data. Negative controls for RUV-III are genes that have reasonable expression levels and whose variation is largely unwanted, i.e. not of biological interest.

We will make use of RNA-seq data from several The Cancer Genome Atlas (TCGA) studies to show that RUV-III with PRPS can lead to highly effective normalization of RNA-seq data that is compromised by substantial library size differences, tumor purity variation and batch effects. We also present comprehensive strategies for revealing unwanted variation in large-scale RNA-Seq studies such as those of the TCGA project.

## Results

### TCGA RNA-seq datasets

The TCGA research network generated RNA-seq data from ∼11000 tumor samples obtained from 33 cancer types. To understand some potential sources of unwanted variation, fresh frozen tissue samples were collected from tissue source sites (TSS), allocated to 96-well sequencing-plates (hereafter called plates), and processed at various times (Table S1). Some TCGA RNA-seq datasets such as uveal melanoma and uterine carcinosarcoma were generated using a single plate. In general, plates are completely confounded with times, making it difficult to distinguish plate effects from time effects. There are also formalin-fixed paraffin-embedded samples among the TCGA RNA-seq samples, and these were excluded from the data discussed here.

Harmonized RNA-seq datasets (see Methods) are available in the form of raw gene counts, FPKM (fragments per kilobase per million) and FPKM, followed by upper-quartile normalization (FPKM.UQ). Low-quality samples and lowly expressed genes were excluded from the datasets prior to the analyses in this paper (see Methods).

### Library size, tumor purity, and plate effects are major sources of unwanted variation across TCGA RNA-seq datasets

We first considered the role of sample RNA-seq library size as a source of unwanted variation. For most cancer types, library sizes vary greatly both within and between years (Figure 1A). The first five principal components (PC) cumulatively are strongly associated with (log) library size in the raw gene counts (Figure 1B, first panel). Ideally, the gene expression levels of genes should have no association with library size in a well-normalized data. Consequently, any downstream analysis including dimensional reduction, gene co-expression, differential expression should also not be influenced by library size variation. The FPKM and FPKM.UQ normalizations reduced the effects of library size, but they showed shortcomings, high correlation between PCs and library size, in several cancer types (Figure 1B, first panel).

**Figure 1:**
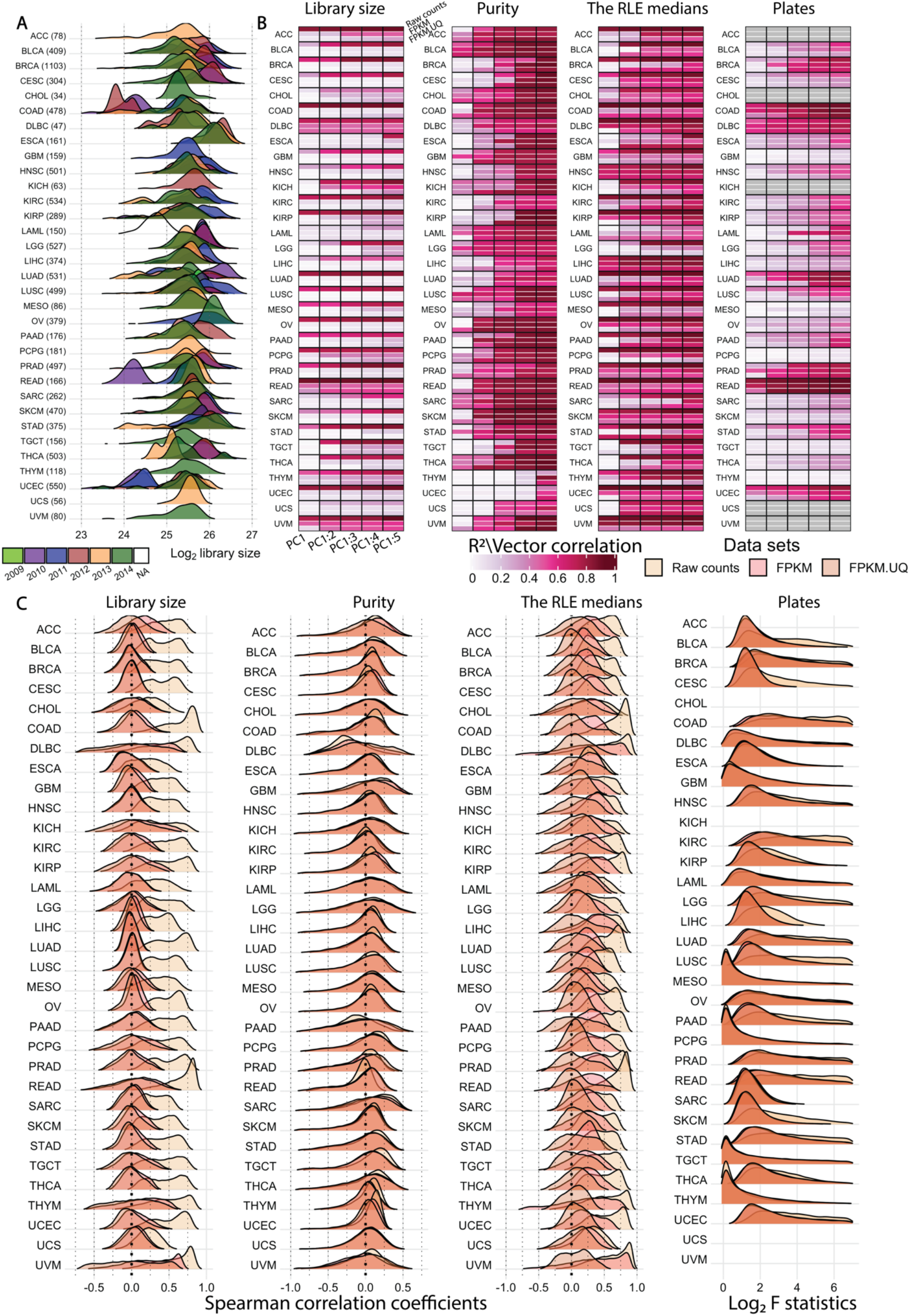
Unwanted variation in individual TCGA RNA-seq datasets. **A)** Distribution of (log_2_) library size coloured by years for each cancer type. The *year information* was not available for the LAML RNA-seq data. The library sizes are calculated after removing lowly expressed genes for each cancer type. **B)** R^2^ obtained from linear regression between the first, first and second, etc., cumulatively to the fifth principal component (PC) and library size (first panel), tumor purity (second panel), and Relative Log Expression (RLE) median (third panel) in the raw count, FPKM and FPKM.UQ normalized datasets. The fourth panel shows the vector correlation between the first 5 PC cumulatively and sequencing-plates in the datasets. Grey colour indicates that samples were profiled across a single plate. **C)** Spearman correlation coefficients between individual gene expression levels and library size (first panel), tumor purity (second panel) and RLE median (third panel) in the datasets. The fourth panel shows log_2_ F statistics obtained from ANOVA of gene expression levels by the factor: plate variable. Plates with less than three samples were excluded from the analysis. ANOVA was not possible for cancer types whose samples were profiled using a single plate.

For each cancer type, the association between an individual gene’s expression level and library size was quantified using Spearman correlation (Figure 1C, first panel). The results show that a large proportion of genes have high positive correlations with library size in the raw gene count datasets. However, there are reasonable numbers of genes whose expression levels have no correlation or a negative correlation with library size (Figure 1C, first panel). Supplementary Figure 1 shows that the association between gene-level raw counts and library size is partially explained by average gene expression level and is never constant. The FPKM and FPKM.UQ normalizations, which rely on a global scale factor to remove library size effects, create unwanted variation in the expression levels of genes whose expression has no association with library size, and increase the level of this unwanted variation in genes whose expression levels are negatively associated with library size. Supplementary figure 2A shows these issues are obvious across all the TCGA normalized datasets. This will be discussed in more detail for the READ and COAD datasets.

Next, we considered tumor purity as a source of unwanted variation. Using linear regression and Spearman correlation analyses we show that tumor purity is major source of variation in the TCGA datasets (Figure 1B, second panel and Figure 1C, second panel). FPKM and FPKM.UQ normalizations cannot correct for purity in cancer RNA-seq data. Thus, this variation remained in the TCGA normalized datasets (Figure 1B, second panel, and Supplementary Figure 2B). We discuss how tumor purity can compromise downstream analyses including gene co-expression and subtype identification, as was observed in the TCGA BRCA RNA-seq data.

The TCGA biospecimens were profiled across plates. Vector correlation analysis (see Methods) indicates the presence of plate effects in the raw gene counts and in the FPKM and FPKM.UQ normalized counts (Figure 1B, third panel). ANOVA analyses between individual gene expression levels and plate variables also show that large numbers of genes remain strongly associated with plates in the normalized datasets (Figure 1C, fourth panel). We have observed that the major biological populations are well distributed across plates in TCGA rectum, colon and breast cancer RNA-seq data indicating the absence of large confounding effects in the data.

Finally, we examined the medians of relative log-expression (RLE) [13] for the raw count and TCGA normalized data (see Methods). In the absence of unwanted variation, the RLE medians should be centred around zero, so any deviation from zero indicates the presence of unwanted variation.

Supplementary Figure 3 illustrate that the RLE medians of the raw count datasets deviate greatly from zero, which further confirms the presence of unwanted variation. Further, we examined the associations between the first five PC cumulatively and the RLE medians, and also assessed the correlation between individual gene expression with the RLE medians for each cancer type (Figure 1B, fourth panel). Ideally, we should see no associations; however, we see numerous associations in the raw counts and the FPKM and FPKM.UQ normalized datasets.

The association between gene expression and RLE medians were quantified by Spearman correlation (Figure 1C, third panel). Below we discuss the value of this analysis for the TCGA colon, rectum and breast cancer RNA-seq data.

Taken together, our results showed that all the TCGA RNA-seq datasets, both raw and normalized, are greatly affected by the three major sources of unwanted variation. Next, we use the rectum adenocarcinoma (READ), colon adenocarcinoma (COAD), and breast invasive carcinoma (BRCA) RNA-seq datasets to illustrate the effects of unwanted variation on certain down-stream analyses and show the performance and effectiveness of RUV-III with PRPS for these datasets. The details of each study are given separately below.

### TCGA rectum adenocarcinoma RNA-seq study

#### Study outline

The rectum adenocarcinoma (READ) RNA-seq study involved 176 assays generated using 14 plates over four years (Supplementary Figure 4). The RNA-seq library sizes vary greatly between samples profiled in 2010 and the rest of the samples (Supplementary Figure 4). We identified consensus molecular subtypes (CMS) using the R package CMScaller [14] in the data normalized by different methods. These subtypes will be used for both assessing the performance of normalization methods and creating pseudo-replicate of pseudo-samples for RUV-III normalization. See Supplementary Figures 5 and 6, and the Supplementary File for further details.

#### RUV-III removes substantial library size differences and plate effects in the data

Large library size differences between samples profiled in 2010 and the rest of the samples are clearly visible in the RLE and PCA plots (Supplementary Figure 7A and Figure 2A, top panel) of the raw count data. Although the FPKM and FPKM.UQ normalizations reduce the library size variation, both methods exhibited shortcomings by not fully mixing samples from different times (Figure 2A, top panel). PCA plots and linear regression between the first ten PC (cumulatively) and library size clearly illustrate that RUV-III with PRPS improved upon the FPKM and FPKM.UQ normalizations in removing the library size effects from the data (Figure 2A, top panel, and Figure 2C). Henceforth, we will assume that RUV-III is always carried out with PRPS. Spearman correlation analyses between the individual gene expression values and library size show that there are genes having clearly positive or clearly negative correlations with library size in the FPKM and FPKM.UQ normalized datasets, whereas most genes were not very correlated with library size in the RUV-III normalized data (Figure 2D). Further, differential expression analysis (see Methods) was performed between samples with high and low library size. The results were displayed in the form of p-value histograms (Figure 2E). Ideally, we should see little evidence of differential expression in these plots, whereas we see a lot in the FPKM and FPKM.UQ datasets, far more than in the RUV-III normalized data. Finally, the silhouette coefficient and Adjusted Rand Index (ARI) analyses (see Methods) showed that RUV-III effectively mixed samples across different key time intervals (Figure 2F).

**Figure 2.**
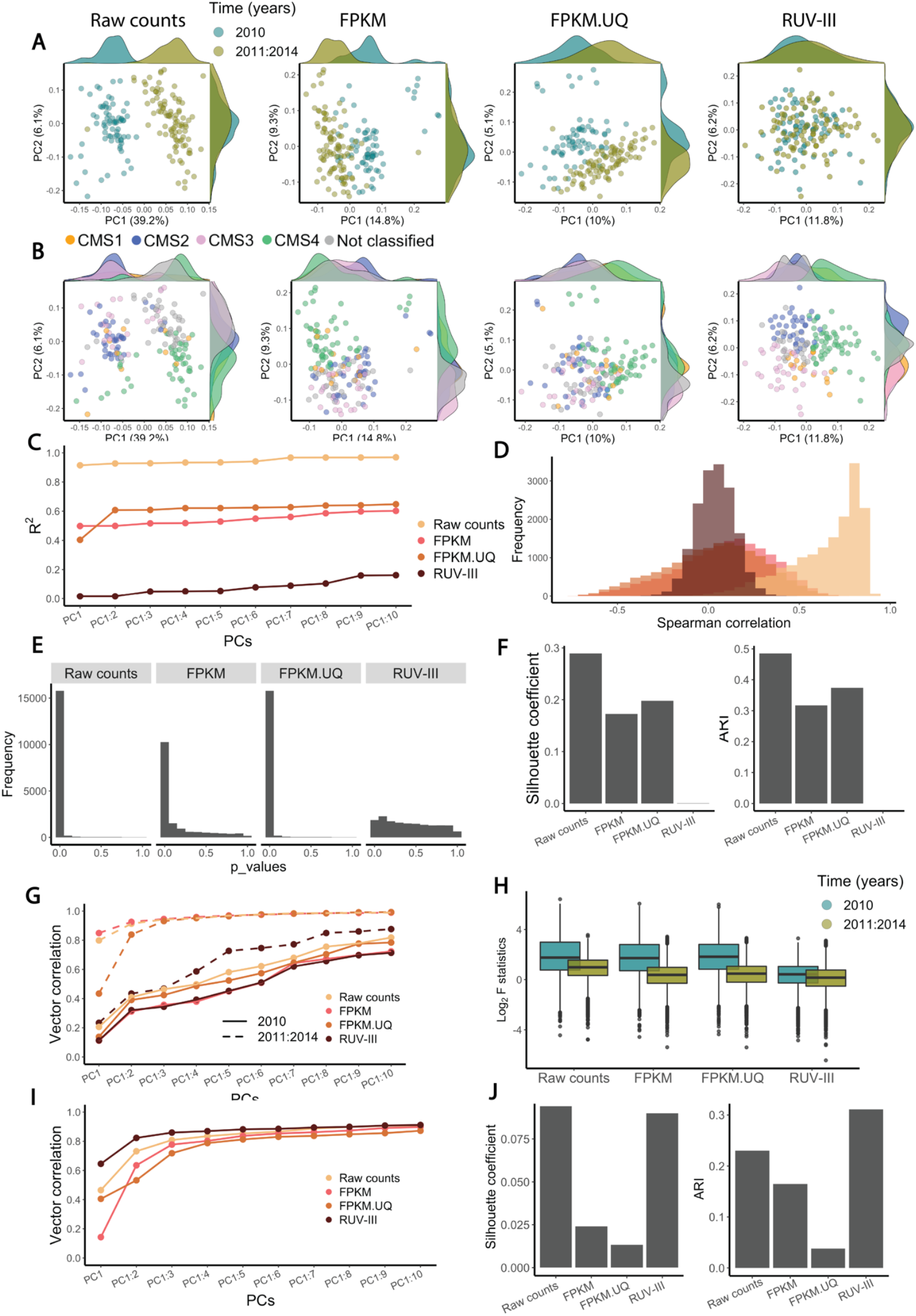
Normalization method performance on the TCGA READ RNA-seq data. **A)** The top row shows scatter plots of first two principal components for raw counts, FPKM, FPKM.UQ and RUV-III normalized data coloured by key time points (2010 vs. 2011-2014). **B)** Same as A coloured by CMS. **C)** A plot showing the R-squared (R^2^) of linear regression between library size and up to the first 10 principal components (successively). **D)** Histograms of Spearman correlation coefficients between the gene expression levels and library size. **E)** P-value histograms obtained from differential expression analysis between samples with low and high library size. **F)** Silhouette coefficients and ARI index for mixing samples from two different key time point. **G)** A plot showing the vector correlation coefficient between plates and up to the first 10 principal components within each time point. **H)** Boxplots of log2 F statistics obtained from ANOVA between gene expression and plates within each time points. **I)** A plot showing the vector correlation coefficient between CMS subtypes and up to the first 10 principal components. **J)** Silhouette coefficients and ARI index for messaging the separation of CMS subtypes.

To examine plate effects and separate this variation from the large library size differences in the data, we performed our evaluation within each key time interval. The results show that RUV-III clearly improves over the FPKM and FPKM.UQ normalizations in removing plate effects from the data (Figure 2G,H).

It should be noted that here we have not attempted to remove tumor purity variation from these data. Consequently, the tumor purity estimates obtained from the RUV-III and FPKM.UQ normalized data showed a high correlation with each other (Supplementary Figure 7B). This illustrates the ability of RUV-III to remove just the variation the user wishes to remove and no more, i.e. to retain other variation that is of biological origin.

We next explore the relationship between RLE medians and both library size and tumor purity for the differently normalized datasets (Supplementary Figure 7C). In the raw counts and the TCGA normalized datasets, RLE medians are strongly associated with both library size and tumor purity. The RLE medians of the RUV-III normalized data also show a strong association with tumor purity (Supplementary Figure 7D). These results were supported by comparisons of the Spearman correlation analyses between the individual gene expression levels and RLE medians with the same analyses between the individual gene expression levels and library size and with tumor purity (Supplementary Figure 8). Together, these results show the value of exploring the association of the RLE medians with known sources of unwanted variation in the data. Later we will show that the RLE medians have no correlation with gene expression in the TCGA BRCA RUV-III normalized data when variation in both library size and tumor purity were removed.

#### RUV-III improves the separation between consensus molecular subtypes

Colorectal cancers are classified into four transcriptomics-based subtypes, consensus molecular subtypes (CMS), with distinct features. PCA plots of the RUV-III normalized data show distinct clusters for the consensus molecular subtypes (CMS) of the READ RNA-seq samples, whereas these subtypes are not as clearly separated in the TCGA normalized datasets (Figure 2B). To confirm the CMS clusters in the PCA plots of the RUV-III normalized data, we applied PCA within the key time intervals in the FPKM and FPKM.UQ normalized datasets. The results show that the clustering of the CMS within each time interval in the FPKM.UQ data is highly consistent with that obtained with RUV-III using the full set of data. (Supplementary Figure 9). A vector correlation analysis between the first 10 PC (cumulatively) and the CMS confirmed that the RUV-III normalization leads to a better separation of the CMS clusters compared to that found in the TCGA normalized datasets (Figure 2I). These results were strengthened by Silhouette coefficient and ARI analyses (Figure 2J).

Further, gene set enrichment analyses showed that the CMS obtained from the RUV-III normalized data are associated with known gene signatures[14] (Supplementary Figure 6B). In addition, Kaplan-Meier survival analysis illustrated that the CMS obtained from the RUV-III normalized data are significantly associated with overall survival (log-rank test p-value ≈ 0, Supplementary Figure 10) which is consistent with previous studies [15]. These results suggest that the RUV-III normalization better preserves known biological signals in the data associated with CMS.

#### RUV-III improves gene co-expression and gene-level survival analyses

Unwanted variation introduced by sample library size differences can have two effects on estimates of gene co-expression analysis. It can lead to correlations between genes that are most likely uncorrelated. For example, the correlation between the *TMF1* and *BCLAF1* are ρ = -0.8 and ρ = 0.7 in the TCGA FPKM and FPKM.UQ normalized data, respectively, whereas we do not see this in either the RUV-III normalized data or in the TCGA READ microarray data (Figure 3A). Alternatively, unwanted variation can obscure correlations between gene-gene expression levels that likely to be truly correlated. For example, the overall correlation between the *MDH2* and *EIF4H* genes is ρ = -0.05 in the TCGA FPKM and FPKM.UQ normalized data respectively, whereas these genes exhibit a high correlation within each key time interval (Figure 3A). The overall correlation of these genes was 0.7 in the RUV-III normalized data, consistent with what was seen in the TCGA READ microarray data (Figure 3A).

**Figure 3:**
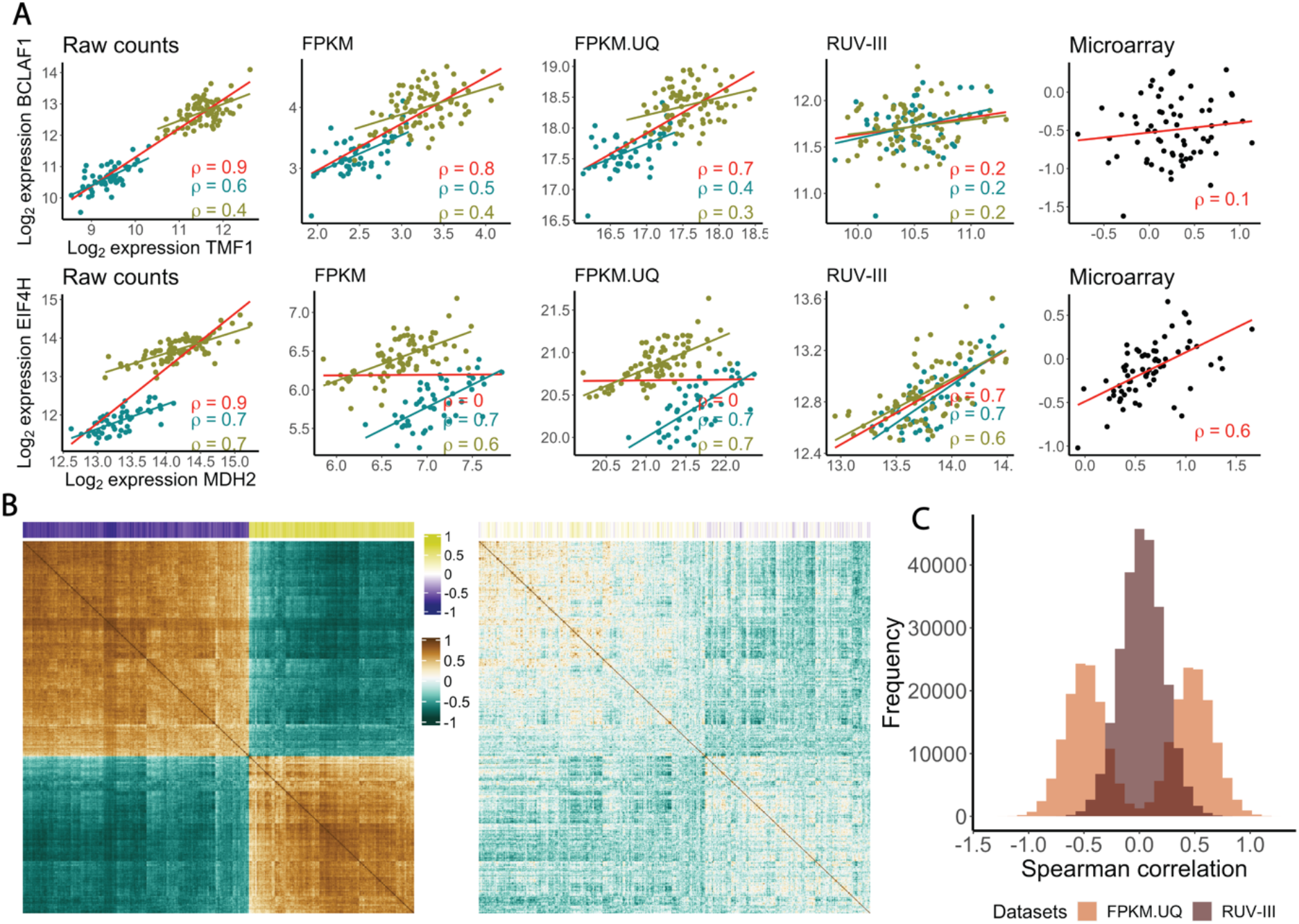
Gene co-expression analyses of TCGA READ RNA-seq data using different normalizations. **A)** First row: scatter plots of the gene expression levels of *TMF1* and *BCLAF1* in the TCGA READ raw counts and differently normalized data. The red line shows overall association, and the green and yellow lines show associations between the gene expression within 2010 samples and within the rest of the samples, respectively. Second row: same as first row, for MDH2 and EIF4H gene expression. **B)** The correlation matrix of expression levels of the genes with the 500 highest correlations with library size in the FPKM.UQ. The first plot is obtained using FPKM.UQ and the second plot is obtained using the RUV-III normalized data. The coloured bar along the top shows the correlation of individual genes with library size. The order of rows and columns is the same in both correlation matrices. **C)** Differences (ρ_microarray_ – ρ_RNA-seq_) of Spearman correlation coefficients for all possible gene-gene pairs calculated using the TCGA READ microarray and both the FPKM.UQ and RUV-III normalized RNA-seq data.

We extended this analysis to all possible gene-gene correlations of the genes that have the highest correlation with library size in the FPKM.UQ normalized data (Figure 3B). Strikingly, the results show numerous strong but likely spurious correlations between gene pairs in the FPKM.UQ normalized data, whereas using RUV-III significantly reduced these correlations (Figure 3B).

Figure 3C depicts the differences (ρ_microarray_ – ρ_RNA-seq_) between all possible gene-gene Spearman correlations ρ using the TCGA READ microarray data and the FPKM.UQ and RUV-III normalized data.

Seeking association between gene expression and survival is another downstream analysis that can be influenced by library size effects that have not been removed. For example, RUV-III revealed that the expression of the genes *RAB18, ATAD1* and *FBXL14* (Figure 4) are highly associated with overall survival in the TCGA READ RNA-seq data. However, the expression of these gene was not associated with survival in the FPKM and FPKM.UQ normalized data. The reason is clear from the expression patterns across time: dividing samples based on median expression was not biologically meaningful for these normalizations (Figure 4). Other examples are *PTPN14* and *CSGALNACT2*, whose associations with survival have been previously shown in colorectal cancer *(data not shown)* [16]. We found a remarkable number of genes whose expression levels were associated with survival using the RUV-III normalized data, which were not found using the FPKM and FPKM.UQ normalized data.

**Figure 4.**
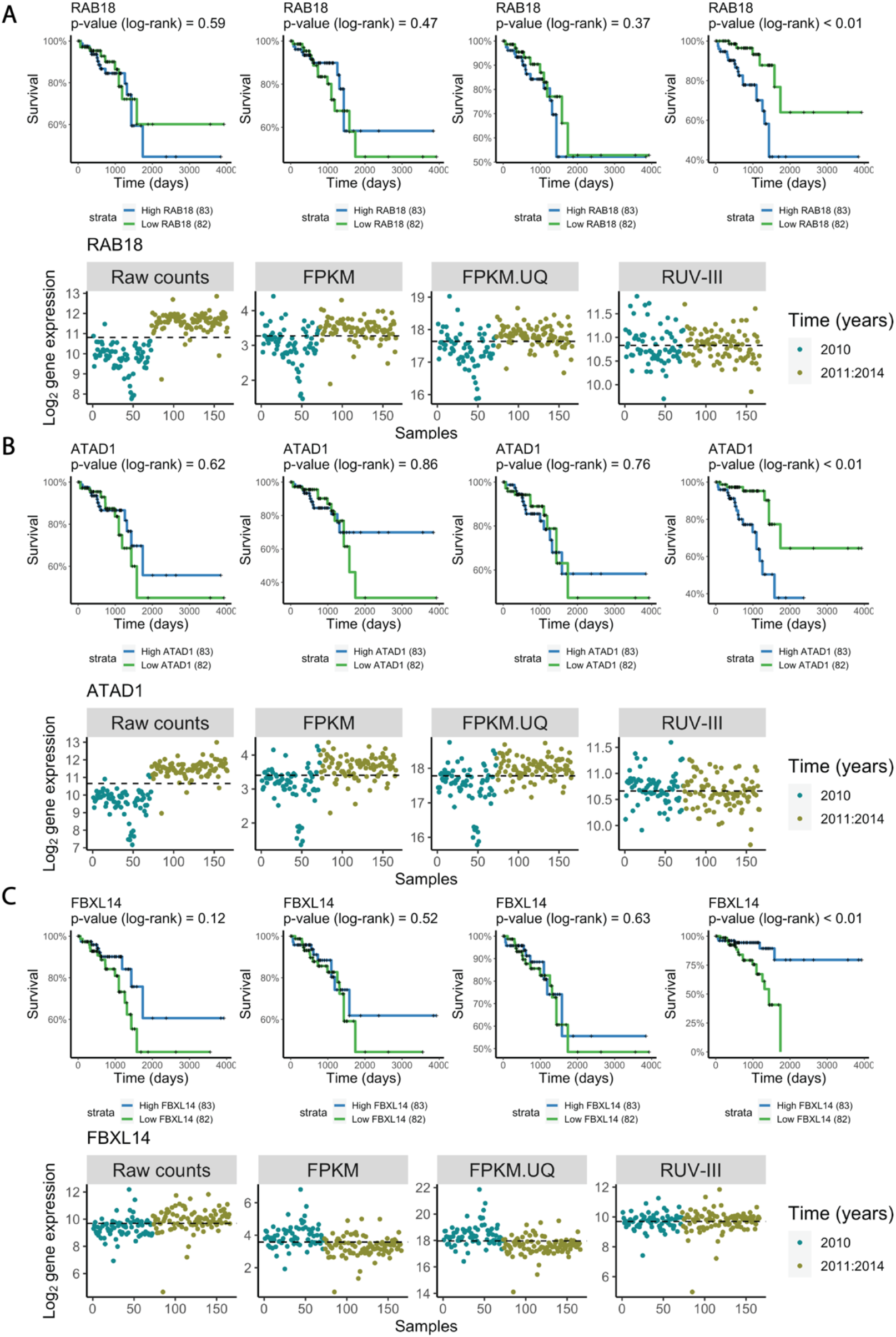
Association between gene expression and overall survival in differently normalized TCGA READ datasets. **A)** Upper part. Kaplan–Meier curves for samples with low (below median) and high (above median) expression of the *RAB18* gene. Lower part. Plots of the expression levels of *RAB18* across time. The dashed lines represent the median expression level of *RAB18*. ***B*** and ***C)*** As in **A**) for the genes *ATAD1 and FBXL14*.

#### Gene-level counts are not proportional to library size

The FPKM and FPKM.UQ normalizations rely on global scale factors based on library size or upper quartiles of samples in the raw count data (Figure 5A) to remove library size effects. These methods assume that gene-level counts all are proportional to the global scale factors. However, we show that in the READ raw count data different groups of genes exhibit different relationships to the global scale factors used in the FPKM and FPKM.UQ normalizations (Figure 5B).

**Figure 5.**
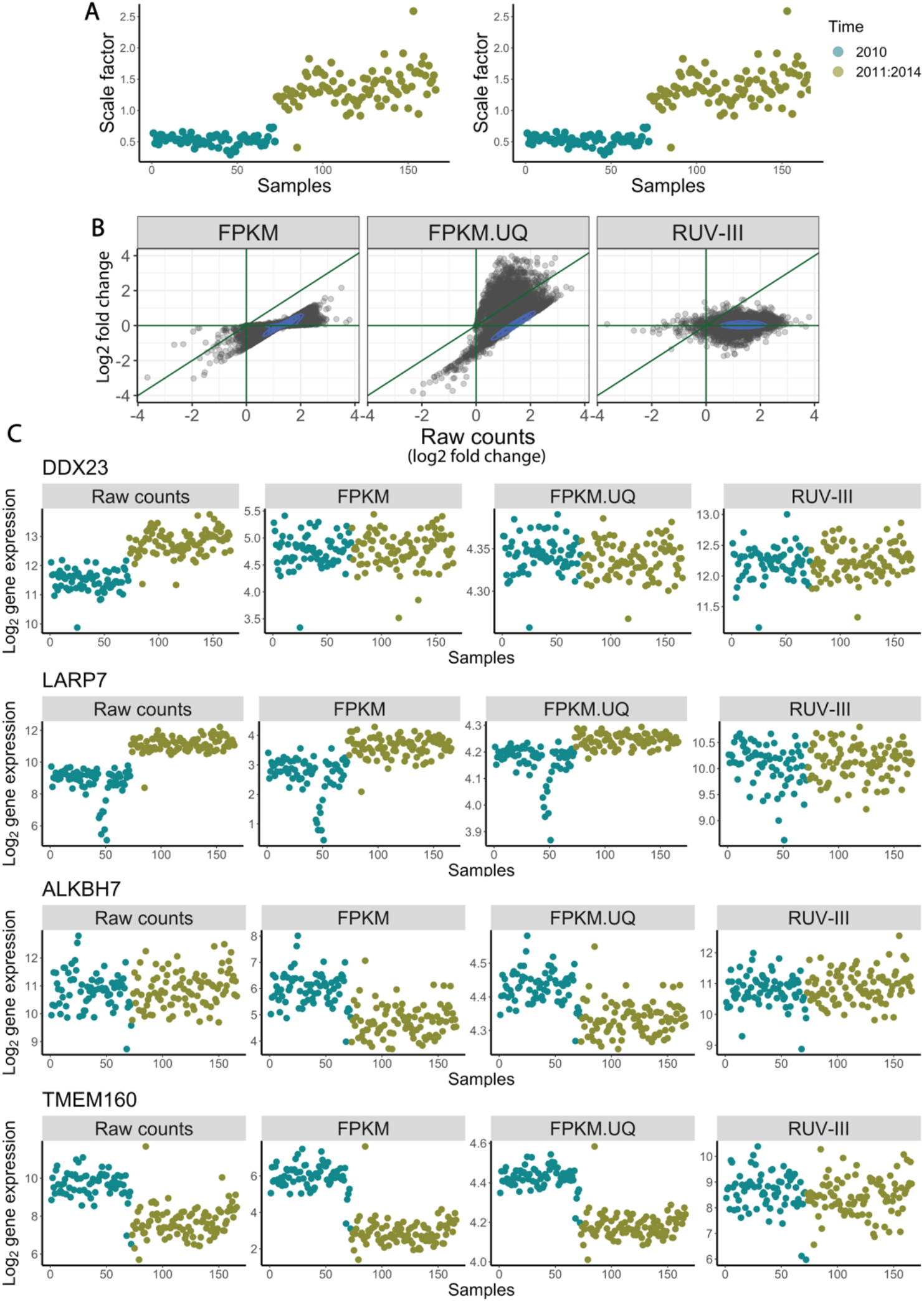
Relationship between gene level (log_2_) counts and (log_2_) library size in the TCGA READ RNA-seq data. **A)** Global scale factors obtained by sample library sizes (left) and upper-quartiles (right) of READ raw counts versus time. **B)** Scatterplots of log_2_-fold change obtained from differential expression analyses of gene expression levels with the major time variation: 2010 vs all other years; (log_2_) raw READ counts on the horizontal axes of all plots and differently normalized counts vertically. **C)** Expression patterns of four genes (*DDX23, LARP7, ALKBH7, TMEM160*) whose counts have different relationships with the global scaling factors calculated from the READ raw count data.

The first group consists of genes whose counts *are* proportional to the global scale factors. For these genes, the FPKM and FPKM.UQ normalizations are adequate to remove library size variation. The gene *DDX23* is an example from this group (Figure 5C, first row). The second group includes genes whose expression levels are *greater* than those expected using the global scaling factors, and so those factors are insufficient for adjusting their expression levels to be independent of library size. The *LARP7* gene represents the behavior of genes in this group (Figure 5C, second row). The third group contains genes such as *ALKBH7*, whose expression levels are not associated with library size in the data. The FPKM and FPKM.UQ normalizations introduce unwanted library size variation to the expression levels of genes in this group (Figure 5C, third row). Finally, there are genes such as TMEM160 whose expression levels relate to library size in a manner *opposite* to that motivating the use of global scaling factors. Applying scaling factors to such genes exacerbates rather than removes variation associated with library size (Figure 5C, fourth row).

It should be noted that we found the same issue in the TCGA RNA-seq datasets such as kidney chromophobe and uveal melanoma, where samples were profiled using a single plate (Figure 1C, first panel).

### TCGA colon adenocarcinoma RNA-seq study

The colon adenocarcinoma (COAD) RNA-seq study involved 479 assays generated across 4 years. As with the READ RNA-seq data, there are large library size differences between samples profiled in 2010 and the subsequent samples. The FPKM and FPKM.UQ normalizations removed library size effects from the data more effectively than was the case for the READ RNA-seq data, but these also had shortcomings. We refer to the Supplementary File and Supplementary Figures 11 to 23 for full details of this dataset and results analogous to those just presented for the READ data.

### TCGA breast invasive carcinoma RNA-seq study

#### Study outline

The breast invasive carcinoma (BRCA) RNA-seq study involved 1180 assays that were carried out on samples from 40 tissue source sites (TSS), distributed across 38 plates, and profiled over five years from 2010 to 2014 (Supplementary Figure 24). The samples profiled in 2010 and 2011 were done using one flow cell chemistry, and the remaining samples were profiled using a different flow cell chemistry (personal communication from TCGA). There were 94 adjacent normal breast tissue samples and 7 paired primary-metastatic samples in the study (Supplementary Figure 24). We used different approaches to identify the PAM50 (prediction analysis of microarray 50) breast tumor intrinsic subtypes for all these samples (refer to the Supplementary File and Supplementary Figures 25 and 26 for full details).

#### RUV-III removes the effects of tumor purity, flow cell chemistries and library size

As with most of the other TCGA RNA-seq studies (Figure 1), tumor purity is one of the major sources of variation in the BRCA study. For this dataset, we designed our PRPS in such a way as to remove the effects of tumor purity as well as other technical variation (see Methods).

Linear regression between the first ten PC cumulatively and tumor purity within the individual PAM50 subtypes showed that the RUV-III normalization substantially removed this variation from the data (Figure 6A). These results were supported by a differential expression analysis between samples with low and high tumor purity, and Spearman correlation analyses between individual gene expression levels and tumor purity within each of the PAM50 subtypes (Figure 6B,C). The variation of tumor purity estimated using the RUV-III normalized data was significantly smaller than that seen in the corresponding measurements on the FPKM.UQ normalized data (Figure 6D).

**Figure 6.**
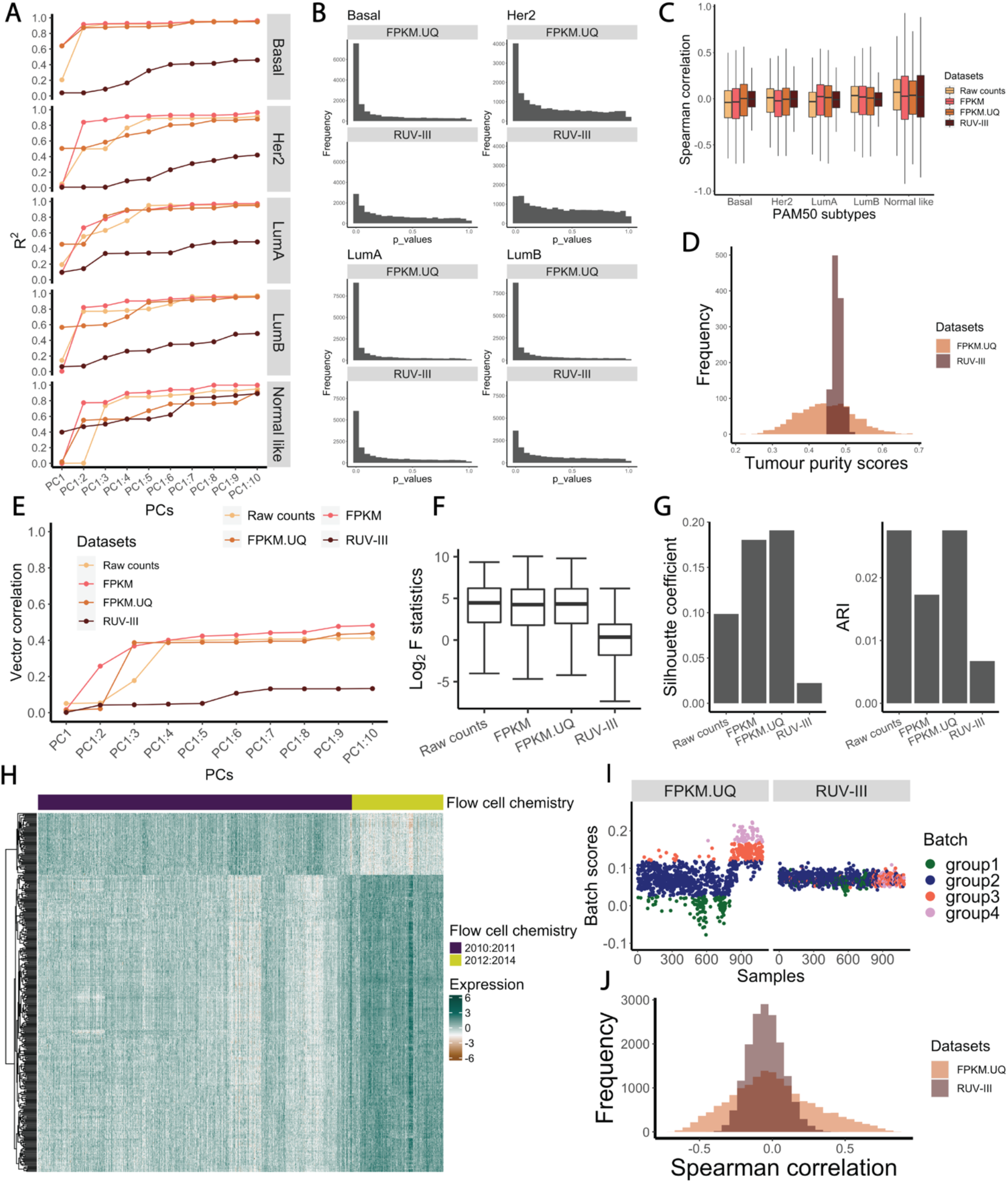
RUV-III removes tumor purity and flow cell chemistry variation from the TCGA BRCA RNA-seq data. **A)** R^2^ obtained from linear regression between the first 10 principal components (cumulatively) and tumor purity within individual PAM50 subtypes in the differently normalized datasets. **B)** *p*-value histograms of differential expression analysis between samples with low and high tumor purity within the four main PAM50 subtypes in the FPKM.UQ and the RUV-III normalized datasets. **C)** Boxplots of Spearman correlation coefficients between individual gene expression and tumor purity levels in the differently normalized datasets. **D)** Distributions of tumor purity scores in the FPKM.UQ and RUV-III normalized datasets. **E)** Vector correlation between the first 10 PC (cumulatively) and flow cell chemistry in the differently normalized datasets. **F)** Boxplots of log_2_ F statistics obtained from ANOVA between individual gene expression levels and the flow cell chemistry factor in the differently normalized datasets. **G)** Bar charts of silhouette coefficients and ARI indices showing the performance of different normalization methods in mixing samples from the two flow cell chemistries. **H)** Gene expression heatmap of 400 genes most highly affected by the flow cell chemistry change in the TCGA FPKM.UQ data (rows are clustered, columns are in chronological order of sample processing. **I)** Batch scores across samples in the FPKM.UQ (left) and RUV-III (right) normalized datasets. The batch scores were calculated by the singscore method using the 400 genes described in H. Samples were divided into 4 groups based on their batch scores. **J)** Spearman correlation coefficients between the batch scores and individual gene expression levels in the FPKM and RUV-III normalized datasets.

PCA plots of the FPKM and FPKM.UQ normalized datasets showed noticeable variation due to the different flow cell chemistries, whereas RUV-III effectively removed this variation from the data (Supplementary Figure 27A). This conclusion was supported by a vector correlation analysis between the first ten PC cumulatively and the binary flow cell chemistry variable, silhouette analyses, the ARI index, and ANOVA between individual gene expression measurements and the flow cell chemistry factor (Figure 6E,F,G and Supplementary Figure 27B and C).

An expression heatmap of the genes most highly affected by the flow cell chemistry difference showed that different genes are affected in different ways (Figure 6H). Interestingly, the heatmap also revealed two clusters within the samples processed by the first flow cell chemistry. This suggests that there are additional sources of unwanted variation of unknown origin within each flow cell chemistry. To explore this more fully, we took the set of genes most highly affected by the difference in flow cell chemistries and scored samples against this gene set (hereafter called the *batch score*) using the R/Bioconductor package singscore [17] on the FPKM.UQ normalized dataset (Figure 6I). Batch scores clearly distinguished samples from the flow cell chemistry batches and separated the samples into clusters within each flow cell chemistry. We then used cut-offs to divide the samples into 4 groups based on their batch scores. These groups were not visible in the batch scores obtained from the RUV-III normalized data (Figure 6I). Spearman correlation analyses showed that a surprising number of genes had either high positive or high negative correlations with the batch scores in the FPKM.UQ normalized data (Figure 6J), whereas these correlations were much lower in the RUV-III normalized data.

#### Tumor purity and flow cell chemistries effects compromise gene co-expression and survival analysis

Just as we saw above with library size, tumor purity variation can affect down-stream analyses such as gene co-expression and the association between gene expression levels and survival outcomes. As with library size, this variation can introduce correlation between genes that are probably un-correlated. For example, Figure 7A shows that the gene expression levels of *ZEB2* and *EST1* are both highly correlated with tumor purity. They also appear to be highly correlated with each other, but this is most likely a consequence of their correlations with tumor purity. The RUV-III normalized data and the breast cancer laser microdissection microarray data [18] showed that the expression levels of these two genes are uncorrelated (Figure 7B). To extend this observation, we selected 1300 genes whose gene expression levels are highly correlated with tumor purity and then calculated Spearman correlation between all possible pairs of these genes. In a matching analysis, we computed partial correlations between these pairs adjusting for tumor purity (see Methods). Figure 7C shows that there are many gene pairs that have high correlations, but these are mostly likely a consequence of their correlation with tumor purity.

**Figure 7.**
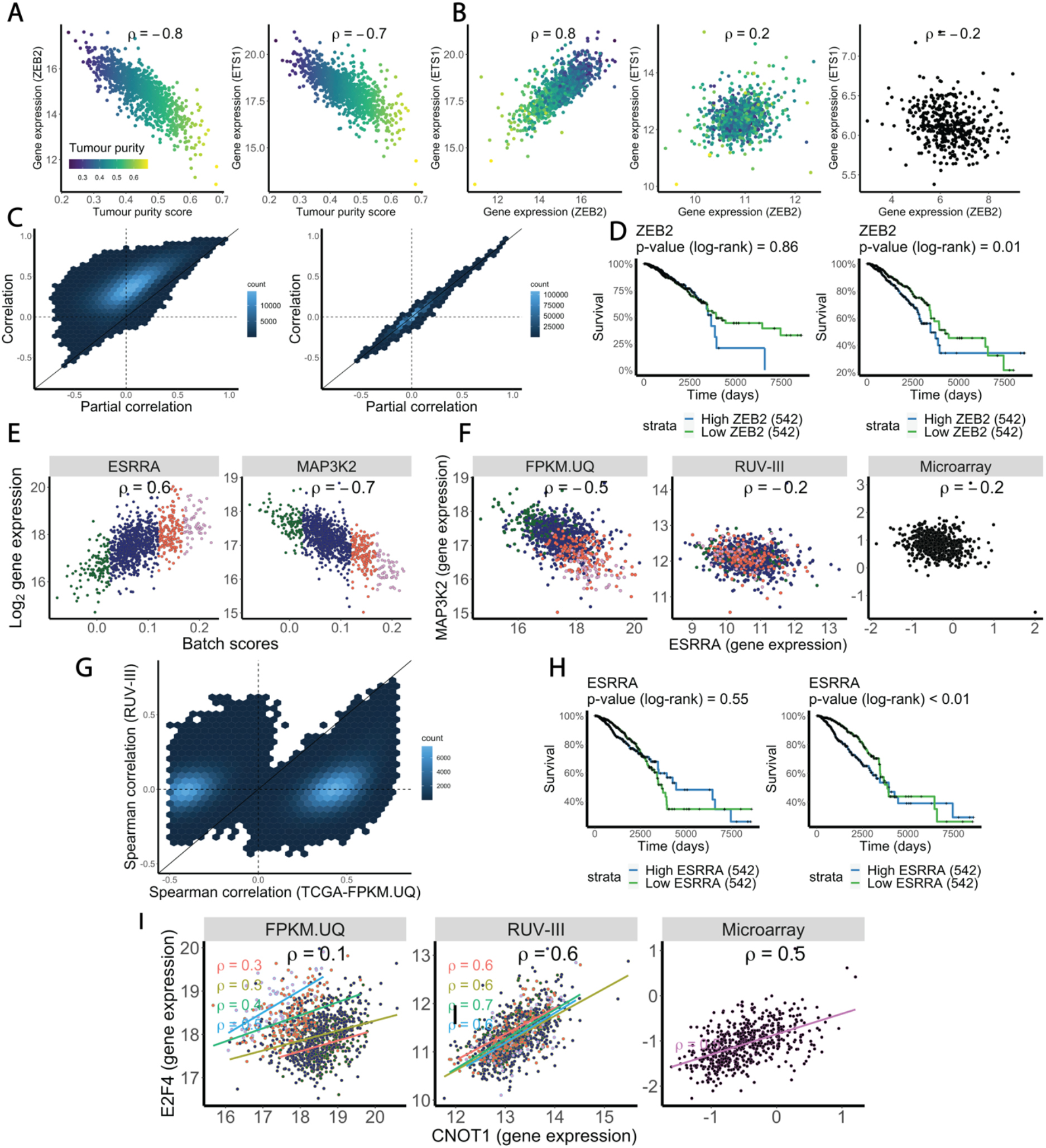
Impact of tumor purity and flow cell chemistry variation on gene co-expression and survival analysis in the TCGA BRCA RNA-seq data. **A)** relationship between tumor purity scores and the ZEB2 and ETS1 gene expression in the FPKM data. **B)** Scatter plots exhibit relationship between the ZEB2 and ETS1 gene expression in the FPKM data (left), the RUV-III normalized data (middle), and the laser capture microdissection microarray data (right). **C)** Scatter plots show the Spearman correlation coefficients and partial correlation coefficients for all possible pairs of the genes that have the 1300 highest correlations with tumor purity in the TCGA FPKM.UQ (left) and RUV-III normalized data (right). **D)** Kaplan Meier survival analysis shows the association between the ZEB2 gene expression and overall survival in the FPKM.UQ (left) and the RUV-III normalized data (right). **E)** Relationship between the *ESSRA* and *MAP3K2* gene expression with the batch scores in the FPKM.UQ data. **F)** Scatter plots show the relationship between the ESSRA and MAP3K2 gene expression in the FPKM.UQ (left), the RUV-III normalized data (middle) and the TCGA BRCA microarray data (right). **G)** Scatter plots display Spearman correlation coefficients of all possible pairs of genes that highly affected by flow cell chemistries in the FPKM.UQ and the RUV-III normalized data. **H)** Kaplan Meier survival analysis shows the association between the ESSRA gene expression and overall survival in the FPKM.UQ (left) and the RUV-III normalized data (right). **I)** Scatter plots exhibit the relationship between the E2F4 and CNOT1 gene expression in the FPKM.UQ (left), the RUV-III normalized data (middle) and the TCGA BRCA microarray data (right).

Variation in tumor purity can also affect the association between gene expression levels and survival outcomes. High levels of expression of the gene *ZEB2* are associated with a poor outcome in the RUV-III normalized data, but this was obscured by variation in tumor purity in the FPKM.UQ normalized data. Another example is the gene *STAB1*, whose expression levels are associated with survival in several cancer types, including breast cancer [19-21]. However, this association was only evident in the present data after removing variation in tumor purity. We found many more examples of such genes using the RUV-III normalized data.

The complex unwanted variation arising from the change in flow cell chemistry and the unknown source noted above clearly compromises estimates of gene co-expression in the FPKM.UQ normalized dataset. It introduces correlations between pairs of genes that are most likely not correlated. For example, the expression levels of the genes *ESRRA* and *MAP3K2* are positively correlated in this dataset, suggesting a shared co-expression network or that these two genes are regulated by the same transcriptional regulator. This would be a novel finding, but this correlation seems to be a consequence of the unwanted variation in the data (Figure 7E), for we do not see it in either the

RUV-III normalized data or the TCGA BRCA microarray data (Figure 7F). To extend this analysis, we first selected the genes that had the 1000 highest correlations with the batch scores in the FPKM.UQ normalized data and calculated all gene-gene correlations between them in both the FPKN.UQ and RUV-III normalized datasets. Figure 7J shows that a large number of gene pairs have high correlations in the FPKM.UQ normalized data, something we do not see in the RUV-III normalized data. Interestingly, the overall correlation in the FPKM.UQ normalized data between expression of the genes *E2F4* and *CNOT1* genes is *ρ = 0*.*1*, while the average of the correlations of these genes *within* each of groups 1 to 4 of the unknown source of unwanted variation is *ρ = 0*.*6* (Figure 7I). Both the RUV-III normalized and the TCGA microarray data show a high positive correlation between the expression levels of the *E2F4* and *CNOT1* genes.

Supplementary Figure 28 shows that the RUV-III normalization removed library size and plate effects from this dataset more effectively than was the case with the FPKM and FPKM.UQ normalizations.

#### RUV-III improves the separation of the PAM50 clusters

Breast cancers intrinsic subtypes including Luminal A, Luminal B, HER2-enriched, Basal-like, and Normal-like [22], are based on a 50-gene expression signatures (PAM50)[23]. PCA plots, vector correlation between the first ten PC cumulatively and the PAM50 subtypes, Silhouette coefficients, and ARI (Figure 8A,B,C) all show that the RUV-III normalization led to better separation of PAM50 subtypes in the BRCA RNA-seq data. Kaplan-Meier survival analysis shows that the PAM50 calls obtained using RUV-III normalized data exhibit stronger associations with survival than do those obtained using the FPKM or FPKM.UQ normalizations (Supplementary Figure 25B).

**Figure 8.**
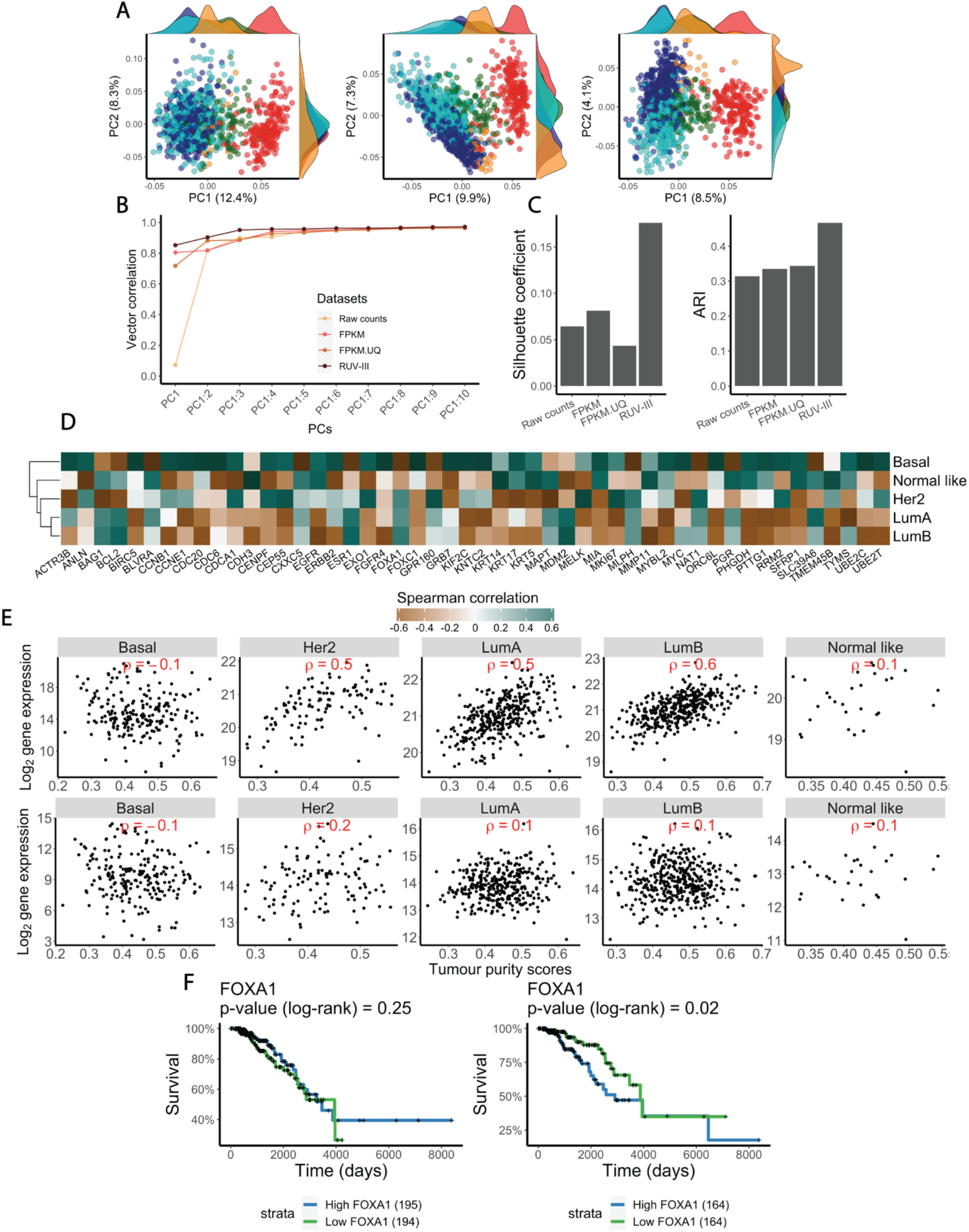
RUV-III improves the PAM50 clusters in the TCGA BRCA RNA-seq data. **A)** Scatter plot of the first two principal components colored by the PAM50 subtypes in the FPKM (left), FPKM.UQ (middle), and the RUV-III (right) normalized datasets. **B)** Vector correlation analysis between the first 10 principal components (cumulatively) and the PAM50 subtypes in the differently normalized datasets. C) Silhouette coefficients and ARI showing how the PAM50 clusters are separated in the differently normalized datasets. D) The heatmap gives the Spearman correlation coefficients between the expression levels of the PAM50 signature genes and the tumor purity scores in the FPKM.UQ data. E) Scatter plots show relationship between the gene expression levels of FOXA1 and tumor purity within the individual PAM50 subtypes in the FPKM.UQ (first row) and the RUV-III normalized data (second row). F) Kaplan Meier survival analyses for samples with low and high expression of FOXA1 gene in Luminal-A subtype in the FPKM.UQ (left) and the RUV-III normalized data (right).

Additionally, Spearman correlation analysis showed that several PAM50 genes exhibit high correlation with tumor purity in the FPKM.UQ normalized data (Figure 8D). For example, expression of the gene *FOXA1* is highly associated with tumor purity in the Her2, Luminal A and Luminal B subtypes in the FPKM.UQ normalized data (Figure 8E). This observation suggests that variation in tumor purity might compromise the identification of PAM50 subtypes. In addition, this might also explain the differences between the PAM50 calls obtained from RUV-III normalized data, where the variation of tumor purity has been removed, and those obtained from the FPKM and FPKM.UQ normalized datasets (Supplementary Figure 25A).

We explored the association between the expression levels of the PAM50 genes and survival within each of the PAM50 subtypes using both the FPKM.UQ and RUV-III normalized data. Interestingly, we found with the RUV-III normalized data that higher expression of *FOXA1* is associated with poorer outcome in the Luminal A subtype, a conclusion that was obscured by tumor purity in the TCGA FPKM.UQ data (Figure 8F).

## Discussion

The main goal of RNA-seq normalization is to effectively remove unwanted variation while preserving variation due to biology. A suitable normalization method for cancer RNA-seq data must be able to remove variation introduced by library size, batch effects, tumor purity (where appropriate), and other technical variation in data.

We proposed a new approach, the use of pseudo-replicates of pseudo-samples (PRPS), to deploy RUV-III for RNA-seq normalization. We made use of data from three TCGA RNA-seq studies to compare the performance of RUV-III plus PRPS with the state-of-art normalizations proposed for RNA-seq data. Our comparisons are based on statistical summaries, biological positive and negative controls, and concordance with the corresponding TCGA microarray data or with other data from independent studies of the same issues.

We began by carefully identifying different sources of unwanted variation in all the TCGA RNA-seq raw count, FPKM and FPKM.UQ normalized datasets. We showed that library size, tumor purity and plate or time effects are major sources of unwanted variation in these studies, and that they can compromise down-stream analyses such as tumor subtype identification, gene co-expression, and association between gene expression and survival. This unwanted variation is likely to affect other down-stream analyses not investigated in this study.

In the TCGA READ RNA-seq study, noticeable library size differences between samples remained in the FPKM and FPKM.UQ normalized data due to the presence of genes whose raw counts showed weak or negative association with library size. In such situations, normalizations which rely on a global scale factor can introduce rather than remove library size variation. We saw that this occurred in the majority of TCGA cancer studies, even those that used a single plate. We took advantage of the ability of RUV-III to remove library size effects only from genes that have associations with library size. RUV-III with PRPS effectively removed the library size effects from the READ RNA-seq data and led to better down-stream analyses of gene-gene co-expression and association of gene expression with survival. We showed that the TCGA normalized COAD RNA-seq data were affected by the same sources of unwanted variation that were found in the corresponding READ RNA-seq data, although their effects were less severe in the COAD compared to the READ data. This shows that even unwanted variation from largely the same sources can affect different cancer studies in different ways.

It should be noted that we did not attempt to remove the effect of variation in tumor purity from the TCGA READ and COAD studies. We then showed that the variation due to tumor purity was similar for the TCGA and the RUV-III normalized datasets. This demonstrated the ability of RUV-III with the PRPS approach to remove just the sources of unwanted variation that users want to remove, and no more. In the TCGA BRCA RNA-seq data, we designed our PRPS to remove variation in tumor purity as well as other sources of unwanted variation including library size, flow cell chemistry and plate effects. The results demonstrated that we achieved this goal. We also showed that variation in tumor purity can compromise cancer subtype identification, gene co-expression and survival analyses. We used laser capture microdissection gene expression data to demonstrate the effects of tumor purity on gene co-expression analysis in the FPKM.UQ normalized data. RUV-III with PRPS effectively removed all tumor purity variation and revealed biological signals which were obscured by this variation.

We showed that the use of two flow cell chemistries introduced unwanted variation into the BRCA RNA-seq data which compromised several down-stream analyses. It introduced correlations between genes that were not truly associated and obscured the correlation between genes that were truly associated. The TCGA consortium assessed 570 breast cancers using both microarray and RNA-seq and so the microarray data could be used as an orthogonal platform to compare the gene expression patterns and their correlations in differently normalized datasets. The results of this comparison showed that the agreement between the RUV-III normalized data and the microarray data was much better than that found with the two TCGA normalized datasets.

It should be noted that we could have created the PRPS using adjacent normal tissue, which is more homogeneous than cancer tissue, had these samples been more uniformly distributed across batches. However, we were not able to use such samples to create PRPS across the two flow cell chemistries for the BRCA study, as all 94 adjacent normal breast tissue samples were profiled using just one of the two chemistries. We found the same problem for the TCGA READ and COAD RNA-seq data. In these datasets, all normal adjacent tissues were profiled within a key time interval.

It should be also emphasised that technical replicates cannot be used to remove variation in tumor purity from the data, as this purity is essentially the same across any set of technical replicates. In large-scale genomics studies such as TCGA, samples are inevitably profiled using different reagents and platforms at different times, which can introduce unwanted variation into the data. RUV-III with PRPS provides a new tool to remove unwanted variation from such large studies. Although we showed the effectiveness of our RUV-III with PRPS approach for normalizing the TCGA RNA-seq datasets with no technical replicates, we still strongly recommend including technical replicates in study designs. We also recommend distributing the biology of interests across batches, to the extent that this is possible, as that can really assist the use of RUV-III with PRPS.

## Material and methods

### Data sets

The TCGA consortium aligned RNA sequencing reads to the hg38 reference genome using the STAR aligner and quantified the results at gene level using the HTseq and Gencode v22 gene-annotation [24]. The TCGA RNA-seq data are publicly available in three formats: raw counts, FPKM and FPKM with upper-quartile normalization (FPKM.UQ). All these formats for individual cancer types (33 cancer types, ∼ 11000 samples) were downloaded using the TCGAbiolinks R/Bioconductor package (version 2.16.1) [25]. The TCGA normalized microarray gene expression data were downloaded from the Broad GDAC Firehose repository (https://gdac.broadinstitute.org), data version 2016/01/28. Tissue source sites (TSS), and batches of sequencing-plates were extracted from individual TCGA patient barcodes (https://docs.gdc.cancer.gov/Encyclopedia/pages/TCGA_Barcode/), and sample processing times were downloaded from the MD Anderson Cancer Centre TCGA Batch Effects website https://bioinformatics.mdanderson.org/public-software/tcga-batch-effects. Pathological features of cancer patients were downloaded from the Broad GDAC Firehose repository (https://gdac.broadinstitute.org). The details of processing the TCGA BRCA RNA-seq samples using two flow cell chemistries were received by personal communication from Dr. K Hoadley. The TCGA survival data reported by Liu *et al*. [26] were used in this paper. Two breast cancer laser captured microdissected (LCM) microarray datasets were downloaded from the NCBI Gene Expression Omnibus (GSE78958 [18], GSE54002 [27]). The consensus measurement of purity estimation (CPE) were downloaded from the Aran *et al*. study [10].

### Filtering samples and genes

We applied the following filtering steps to the individual TCGA RNA-seq datasets. Plates with fewer than 3 samples were removed. Samples with log_2_ library sizes three median absolute deviations lower than the median of all log_2_ library sizes were excluded from the data.

The R/Bioconductor package biomaRt (version 2.48.3) which version was used to annotate genes. All pseudo-genes and immunoglobulin genes were excluded. For the pan-cancer analyses, we retained genes with at least 15 raw read counts in at least 20% of samples. We considered numbers of samples in the biological subpopulations and sources of unwanted variation when removing lowly expressed genes in the TCGA READ, COAD and BRCA data. To do so, we kept genes that have at least 15 counts in the smallest biological subpopulations within each of the key time intervals in the datasets.

### Tumor purity estimates

We estimated tumor purity for all cancer samples using the stromal and immune gene signatures (282 genes, Supplementary Table S2) from Yoshihara *et al*. study [28] and the R/Bioconductor package *singscore* version (1.12.0) [17]. The stromal&immune scores were transformed to 1— stromal&immune scores (singscore tumor purity) for down-stream analyses. These measurements are called tumor purity scores in this study. The tumor purity scores showed high positive correlation (mean = 0.95, Pearson correlation) with the ESTIMATE measurements, and a lower correlation with the CPE from Aran *et al*. study (Supplementary Figure 30).

### Sample library size

Sample library sizes were obtained by adding all gene raw counts for individual samples after removing pseudo-genes, immunoglobulin and lowly expressed genes. All sample library sizes are transformed to log_2_ in this study.

### Pseudo-replicate of pseudo-samples

We used our recently developed normalization method Removing Unwanted Variation III (RUV-III), which makes essential use of technical replicates and negative control genes to estimate unwanted variation and remove it from the data [5]. Ideally, technical replicates are placed across batches so that unwanted variation between any pair of batches is captured via differences of expression values between technical replicates. As there are no technical replicates in the TCGA RNA-seq datasets, we developed a new approach— *pseudo-replicates of pseudo-samples (PRPS)* —to be able to use RUV-III to remove unwanted variation from the data.

In order to use RUV-III with PRPS, we first need to identify relatively homogenous biological sub-populations among the samples and locate the sources of unwanted variation in the data we aim to remove. We illustrate the process for TCGA COAD and BRCA RNA-Seq studies. For example, with the TCGA COAD samples, we regarded the 12 combinations of 4 consensus molecular subtypes (CMS1, CMS2, CMS3, CMS4) with 3 microsatellite instability (MSI) statuses (MSI-high, MSI-low, MS-stable) as defining biological sub-populations. The key time intervals 2010 and 2011-2014 contribute the main unwanted variation that we saw after preliminary exploration of the data. As these times are totally confounded with sequencing plates (i.e. different plates are used across different times), we considered plates to be the source of unwanted variation when defining PRPS. As a result, to remove unwanted plate to plate variation in the COAD data without removing biology, we created sets of *pseudo-samples* as follows: a) select those biological sub-populations out of the 12 mentioned above that have at least 3 samples in at least two plates, while also ensuring that in the end there are samples from plates within and across the two key time points; b) average the gene expression levels of the corresponding samples within the individual plates to create one pseudo-sample. Having done this for all 12 biological sub-populations, we suppose all the pseudo-samples created across plates for a particular biological sub-population to form a pseudo-replicate set.

For the BRCA data, in addition to the sequencing plates, flow cell chemistry was also confounded with two key time intervals (2010-2011 and 2012-2014). We considered the PAM50 subtypes (Basal, Her2, Luminal A, Luminal B and Normal like) to define the biological subpopulations and then created several sets of PRPS across plates in the manner described for the COAD study. For removing the effect of tumor purity in the BRCA data, we defined sets of PRPS for each PAM50 subtype in addition to those that we created for removing plate to plate variation. We did this by selecting the samples with the 5 highest and the samples with the 5 lowest values of tumor purity *within* each PAM50 subtype. Then we created two pseudo-samples within each PAM50 subtype by averaging the gene expression values across each set of 5 high purity samples and each set of 5 low purity samples. Finally, the two pseudo-samples (average high and average low purity) created for each PAM50 subtype were regarded as forming a pseudo-replicate set, i.e. a pair of pseudo-duplicates. These are then used in RUV-III, see below.

### Choice of negative control genes and k

Negative control genes for RUV-III are genes that *are not* highly affected by the biological factors of interest, but *are* affected by one or more forms of unwanted variation in the data. We have previously [5] explained that our approach to negative controls is pragmatic: if regarding a set of genes as negative controls helps to remove unwanted variation using RUV-III, as evaluated by various metrics, then whether or not they are ideal negative control genes is not our concern. For the different cancer types discussed in this paper, we used different sets of negative control genes derived from either the literature (e.g. housekeeping genes or genes found to be stable in the same or a closely related biological context) or the data itself (e.g. genes found to exhibit little or no biological but clear unwanted variation). Candidate control genes have their effectiveness evaluated using various metrics following their use in RUV-III. It should be noted that unwanted variation mostly affects different subsets of genes in different ways.

In order to use RUV-III, a dimension K of unwanted variation needs to be determined. To find a suitable value, we repeated the analysis with a range of values of K and evaluated the quality of each analysis using different statistical metrics and prior biological knowledge. RUV-III is generally robust to overestimating K, but not always.

#### RUV-III normalization with PRPS for READ

As described above, the 11 combinations (we do not have CMS4_MSI-H) of the 4 CMS subtypes called by the CMScaller R package on the READ FPKM and FPKM.UQ RNA-seq data (consensus calls were selected), and the 3 MSI statuses were considered to be homogenous biological populations for the purpose of creating PRPS. Supplementary Figure 31 displays the numbers of each 11 subpopulations within the individual plates. We created pseudo-samples from plates that have at least 2 samples of at least one of the 11 subpopulations population. Supplementary Figure 31B shows the library size of pseudo-samples within each subpopulation.

A set of negative control genes were selected in the following way. First, an ANOVA was carried out on FPKM.UQ normalized gene expression levels using CMS subtypes as the factor, and the genes with lowest F statistics selected (900 genes). PCA plots of the READ RNA-seq raw counts using the negative control genes showed that they capture the differences between the key time intervals and do not capture CMS subtype differences (Supplementary Figure 31C).

#### RUV-III normalization with PRPS for COAD

Here we first defined CMS using the R package CMScaller on the COAD FPKM and FPKM.UQ RNA-seq data and selected the samples receiving the same CMS call for both (406 out of 479 samples). We used these CMS and MSI to define homogenous biological populations for the purpose of creating the PRPS (Supplementary Figure 32). We used a slightly complicated approach to select a suitable set of negative control genes for the COAD study as follows: a) carry out an ANOVA on the FPKM.UQ normalized gene expression values with CMS subtypes as the factor; b) calculate Spearman correlations between FPKM.UQ normalized gene expression values and tumor purity; c) calculate Spearman correlations between FPKM.UQ normalized gene expression values and the average expression level of a set of housekeeping genes [29]; and then d) selecting genes (262 genes) that have lowest F statistics (F statistics < 20) in a), the lowest correlation (ρ < 0.3) in b), and the highest correlations (ρ > 0.9) in c).

PCA plots of the TCGA COAD RNA-seq raw count using negative control genes show that they capture the key time differences (Supplementary Figure 32C)

#### RUV-III normalization with PRPS for BRCA

The PAM50 subtypes were assigned by using the genefu R package with the FPKM and FPKM.UQ normalized data. We selected samples with consensus PAM50 subtypes from the two datasets. Two different groups of pseudo-samples were then created to capture the plate&flow cell chemistries and tumor purity effects (Supplementary Figure 33).

The negative control genes were selected as follows: a) carry out an ANOVA on the FPKM.UQ normalized gene expression values with PAM50 subtype as the factor, within each flow cell chemistry; b) carry out a similar ANOVA with flow cell chemistry as the factor; c) calculate Spearman correlations between FPKM.UQ normalized gene expression and purity values within the PAM50 subtypes; d) calculate similar Spearman correlations with library size but with the raw counts; e) selecting genes (4500 genes) with the lowest F statistics from (F statistics < 20) a), the highest F statistics (F statistics > 100) from b), and the highest correlations (abs(ρ > .07) from c), the highest correlations (ρ > .07) from d). PCA plots of the TCGA BRCA RNA-seq raw count using the negative control genes show that these genes capture all sources of unwanted variation in the data (Supplementary Figure 34C).

### Principal Component Analysis (PCA)

The principal components (in this context also called singular vectors) of the sample × transcript array of log-counts are the linear combinations of the transcript measurements having the largest, second largest, third largest, etc. variation, standardized to be of unit length and orthogonal to the preceding components. Each will give a single value for each sample. In this paper PCA plots are of the second principal component (PC) values vs the first PC values and of the third PC vs the first PC. The calculations are done on mean-corrected transcript log counts, using the R code adopted from the EDAseq (version 2.26.1) R package [30].

### Relative log expression (RLE) plots

RLE plots [13] are used to reveal trends, temporal clustering and other non-random patterns resulting from unwanted variation in a dataset. To generate our RLE plots, we first formed the log-ratio *log[y*_*ig*_*/y*_*g*_] of the raw count *y*_*ig*_ for gene *g* in the sample labelled *i* relative to the median value *y*_*g*_ of the counts for gene g taken across all samples. We then generate a boxplot from all the log ratios for sample *i*, and plotted all such boxplots along a line, where *i* varies in a meaningful order, usually sample processing date. An ideal RLE plot should have its medians centered around zero while its box widths, their interquartile ranges (IQR) should be similar in magnitude, see [14] for more discussion. Because of their sensitivity to unwanted variation, we also examined datasets for possible relationships between RLE medians with potential sources of unwanted variation and individual gene expression levels. In the absence of any influence of unwanted variation in the data we should see no such associations.

### Vector correlation

We used the Rozeboom squared vector correlation [31] to quantify the strength of (linear) relationships between two *sets* of variables such as the first k PCs (1≤k≤5) and dummy variables representing time, batches, or plates. Not only does this quantity summarize the full set of canonical correlations, but it also reduces to the familiar R^*2*^ from multiple regression (see below) when one of the variable sets contains just one element.

### Linear regression

*R*^*2*^ values of fitted linear models are used to quantity the strength of the (linear) relationships between a single quantitative source of unwanted variation such as sample (log) library size or tumor purity and global sample summary statistics such as the first k PC (1≤k≤10). The *lm()* R function was used for such analyses.

### Partial correlation of gene expression controlling for tumor purity

Partial correlation is used to estimate Pearson (linear) correlation between two variables while controlling for one or more other variables [32]. We computed the partial correlation between the expression levels of pairs of genes controlling for tumor purity using the *pcor*.*test()* function from the R package *ppcor (version 1*.*1)* [32].

### Analysis of variance (ANOVA)

Analysis of variance (ANOVA) enables us to assess the effects of a given qualitative variable (which we call a factor) on gene expression measurements across any set of groups (labelled by the levels of the factor) under study. We use ANOVA *F* statistics to summarize the effects of a qualitative source of unwanted variation (e.g. batches) on the expression levels of individual genes, where genes having large *F* -statistics are deemed to be affected by the unwanted variation. We also use ANOVA tests (the *aov()* function in R) to assign *P*-values to the association between tumor purity and molecular subtypes.

### Silhouette coefficient analysis

We use Silhouette coefficients analysis to assess the separation of biological populations and batch effects. The silhouette function uses Euclidean distance to calculate both the similarity between one patient and the other patients in each cluster, and the separation between patients in different clusters. A better normalization method will lead to higher and lower silhouette coefficients for biological and batch labels respectively. The silhouette coefficients were computed using the function *silhouette()* from the R package *cluster* (version 2.1.2) [33].

### Adjusted Rand index (ARI)

The Adjusted Rand Index [34] is the corrected-for-chance version of the Rand Index. The ARI measures the percentage of matches between two label lists. We used the ARI to assess the performance of normalization methods in terms of sample subtypes separation and batch mixing. We first calculated principal components and used the first 3 PC to perform ARI.

### Differential expression analysis

Differential (DE) expression analyses were performed using the Wilcoxon signed-rank test with log_2_ transformed raw counts and normalized data [35]. To evaluate the effects of the different sources of unwanted variation on the data, differential expression analyses were performed across batches. In the absence of any batch effects, the histogram of the resulting unadjusted p-values should be uniformly distributed. The *wilcox*.*test()* R function was used for this analysis.

### Identification of unwanted variation in TCGA RNA-seq datasets

We make use of both *global* and *gene-level* approaches to identify and quantify unwanted variation in RNA-seq datasets (Supplementary Figure 36). These approaches are also used to assess the performance of different normalization methods as removers of unwanted and preservers of biological variation in the data.

Our *global* approaches involve the use of principal component analysis (PCA) plots, linear regression, vector correlation analyses, silhouette coefficients, adjusted rand indices (ARI), and relative log expression (RLE) plots [13] (Figure 1C, first panel). Our PCA plots (see above) are each of the first three principal components (PC) against each other, coloured by known sources of unwanted variation, e.g. time, or known biology, e.g. cancer subtypes. Linear regression is used to quantify the relationship between the first few PC and continuous sources of unwanted variation such as (log) library size. The R^2^ calculated from the linear regression analyses indicates how strongly the PC capture unwanted variation in the data, and we do these calculations cumulatively, i.e. continuous source vs all of {PC_1_,…,PC_k_}, for k = 1,…,5 or 10. Similar to linear regression, we used vector correlation analysis to assess the effect on the data of discrete sources of unwanted variation such as years or year intervals. Silhouette coefficients and adjusted Rand indices (ARI) were used to quantify how well experimental batches are mixed and known biology is separated. Finally, relative log expression (RLE) plots [13] were used to assess the performance of different normalizations in terms of removing unwanted variation from the data. We also explored the relationship between the medians and the inter-quartile ranges (IQR) of the RLE plots with sources of unwanted variation.

The *gene-level* approach includes differential expression analyses between experimental batches, looking at *p*-value histograms and assessing the expression levels of negative control genes (see above), positive control genes (genes whose behaviour we know), Spearman correlation and ANOVA between individual gene expression and sources of unwanted variation. These methods assess and quantify the effects of unwanted variation on individual gene expression levels in the RNA-seq datasets. We refer to the Methods section for more details about the assessment tools.

## Code availability

All scripts were used to generate the main and supplementary figures present here is available on GitHub at: https://github.com/RMolania/TCGA_PanCancer_UnwantedVariation.

## Acknowledgment

We thank Drs Paul Spellman, Hui Shen, Victoria Wang and Vel Gayevskiy for their helpful comments on the near final draft. Thanks to the TCGA Research Network for generating the data used in this study and groups who have made the raw and normalized datasets publicly available.

## Author Contributions

R.M., A.D., A.T.P. and T.P.S. designed the overall approach. R.M, J.G.B., G.O., T.P.S developed the pseudo-replicate of pseudo-samples approach. R.M., M.F., and L.G. performed data analysis. R.M., M.F., A.T.P., and T.P.S. wrote the manuscript, which was revised and approved by all authors.

## Funding

R.M. and A.T.P. were supported by the Galli Medical Trust and the Galli Next Generation Cancer Discoveries Initiative. R.M. was supported by funding from the Ovarian Cancer Research Foundation. A.T.P. was supported by a National Health and Medical Research Council (NHMRC) Senior Research Fellowship (1116955). The research benefitted by support from the Victorian State Government Operational Infrastructure Support and Australian Government NHMRC Independent Research Institute Infrastructure Support. M.F. is funded by the Prostate Cancer Foundation Young Investigator (PCF-YI) Award (USA). A. D, funding for this work from the National Breast Cancer Foundation Grant II-RS-19-108.

## Notes

### Competing Interest Statement

The authors have declared no competing interest.

