## Supplementary File for "Removing unwanted variation from large-scale cancer RNA-sequencing data"

*Corresponding authors

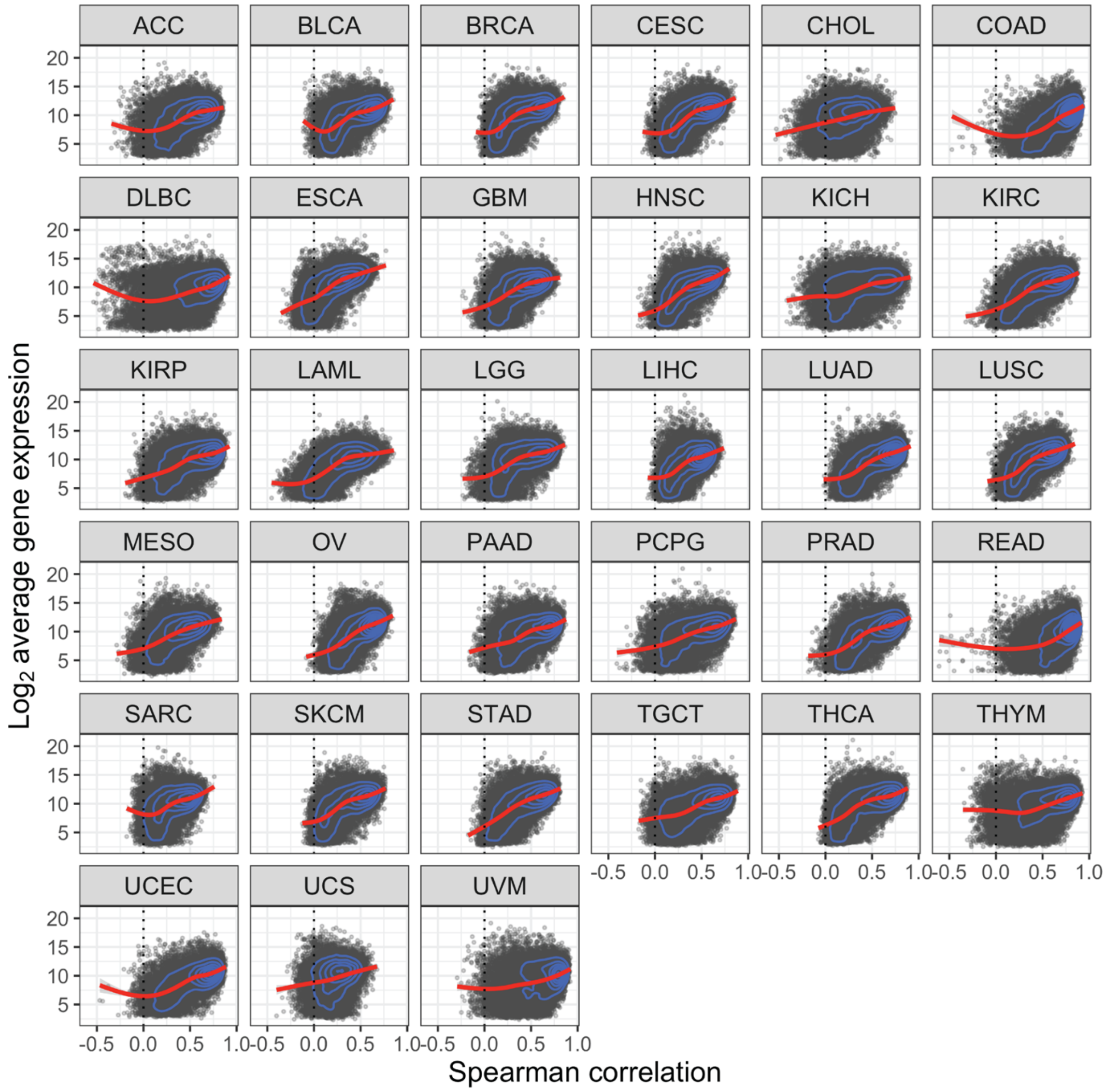


**Supplementary Figure 1. Relationships between average gene expression levels and correlation of raw gene-level counts with library size in the TCGA RNA-sequencing studies.** X axes show Spearman correlation coefficients between $\log_2$ raw gene counts and library size, Y axes show $\log_2$(average (raw count + 1)) the TCGA RNA-sequencing raw counts. Spearman correlations were computed using $\log_2$ of library size and $\log_2$(gene counts+1).

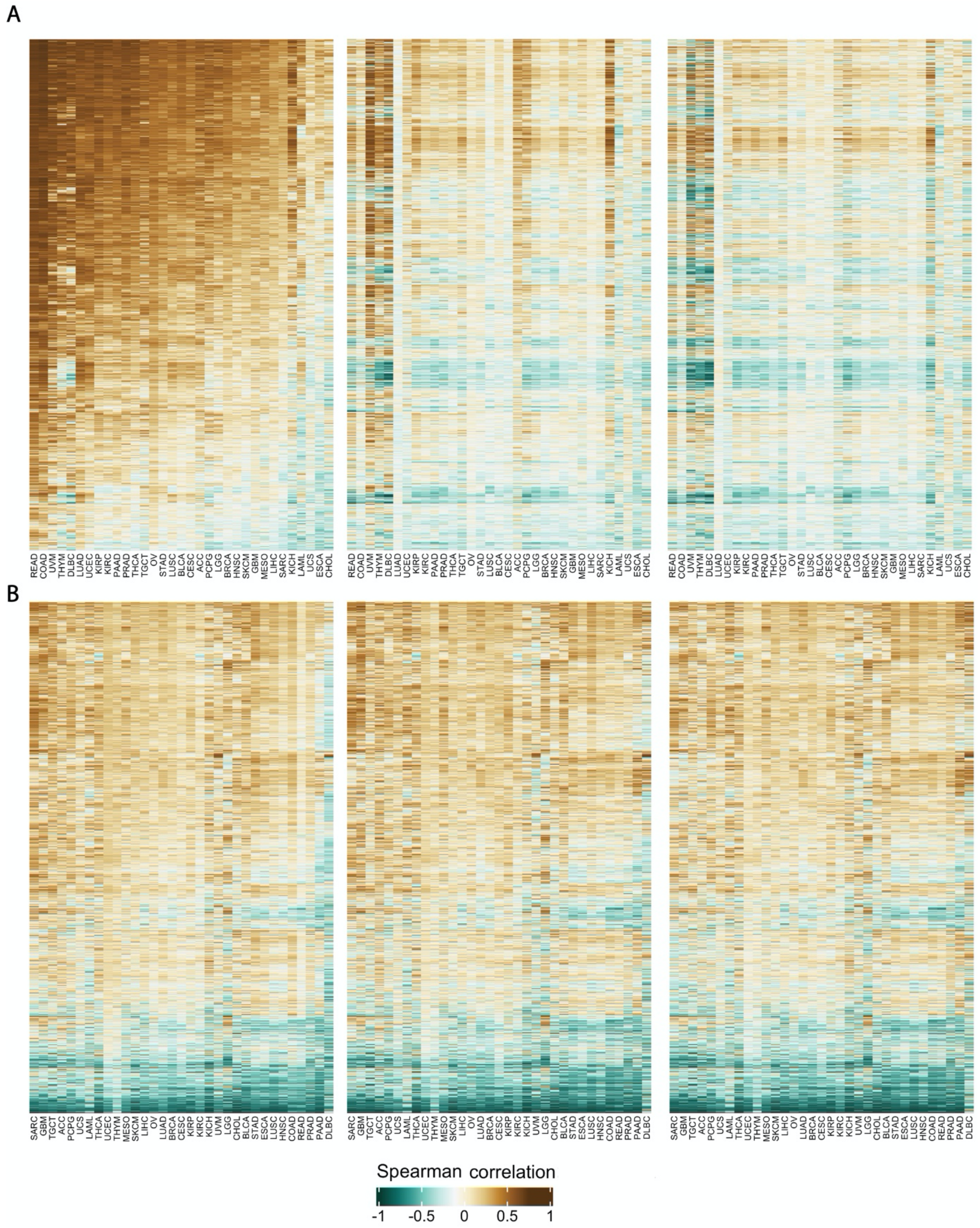


**Supplementary Figure 2. Relationships between gene expression and library size and tumour purity in the TCGA RNA-seq studies. A)** Heatmaps show Spearman correlation coefficients between individual gene expression values and library size in the raw (first heatmap), FPKM normalized (second heatmap), and FPKM.UQ normalized (third heatmap) datasets. **B)** Heatmaps show Spearman correlation coefficients between individual gene expression levels and tumour purity scores in raw counts (first heatmap), FPKM normalized (second heatmap), and FPKM.UQ normalized (third heatmap) counts. The order of genes (rows) and studies (columns) are the same across all heatmaps.

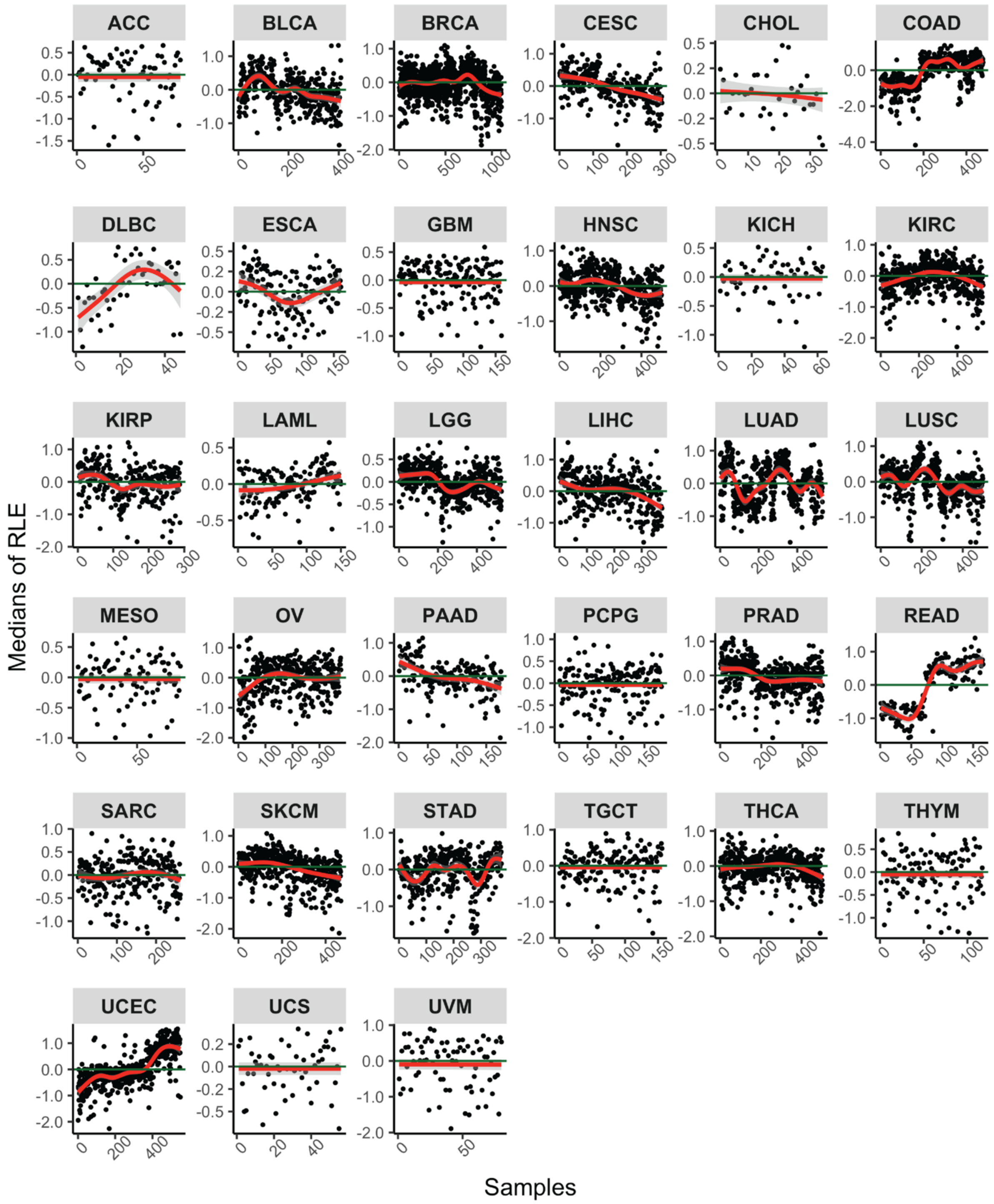


**Supplementary Figure 3. Processing date trends in the relative log expression (RLE) plot medians for the TCGA RNA-seq raw counts.** Lowly expressed genes were removed prior to creating the RLE plots.

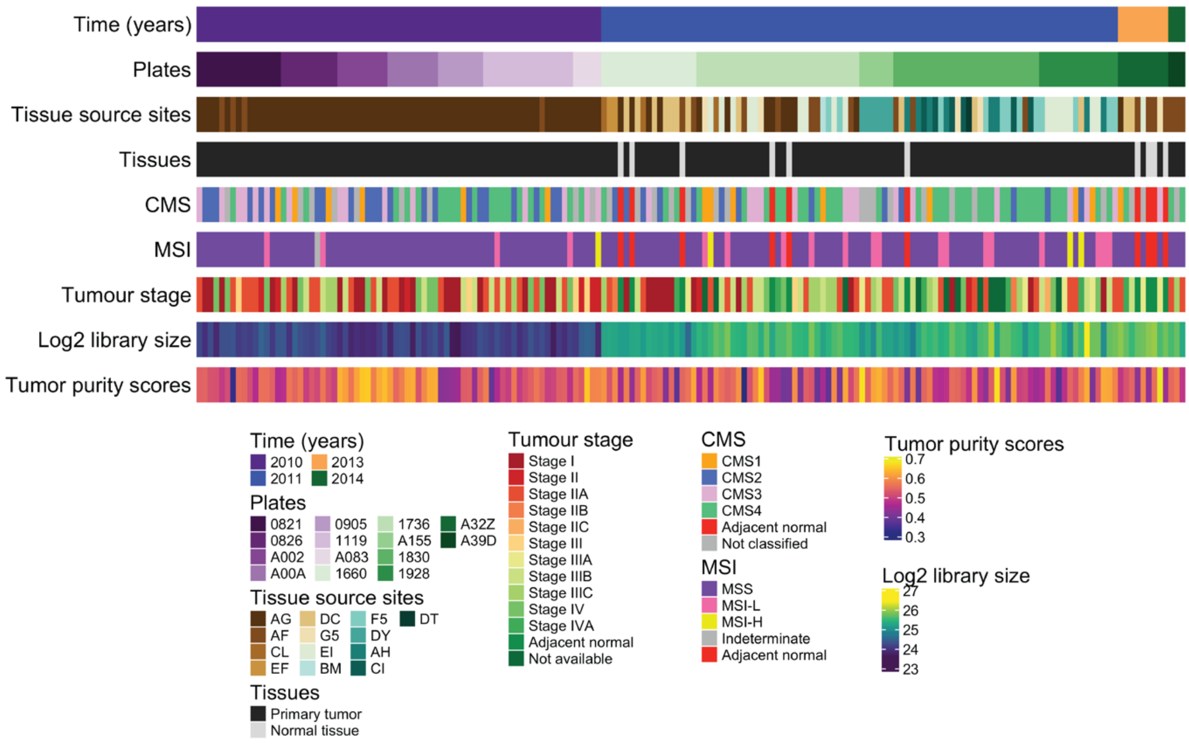


**Supplementary figure 4. Outline of the TCGA rectum adenocarcinoma (READ) study and RNA-seq data. A)** 176 rectum adenocarcinoma and adjacent normal tissues were collected from 13 tissue source sites (TSS) and distributed across 14 sequencing plates for profiling at 14 time points over a span of 4 years. The consensus molecular subtypes were obtained using the R package CMScaller on the FPKM.UQ normalized data. MSI and tumour stage assignments were obtained from the pathological reports in the TCGA clinical data. The library sizes are calculated after removing lowly expressed genes and $log_2$ transformed. The tumour purity scores are obtained from 1- stromal&immune scores (see Methods).

### The consensus molecular subtypes of the TCGA READ RNA-seq samples

Colorectal cancers (CRCs) can be classified into four widely accepted consensus molecular subtypes (CMS) based on their gene expression profiles [1] . This classification provides a framework for stratifying the treatment of patients with colon and rectum cancer. We used the CMScaller R package [2] to identify the CMS of the TCGA READ RNA-sequencing samples. The CMScaller provides a classification based on pre-defined cancer-cell intrinsic CMS templates. We applied that classifier to all samples and also to samples within the key time intervals (2010 and 2011:2014) using the FPKM and FPK.UQ normalized datasets (Supplementary Figure 5). The reason for applying the classifier within each key time interval was to assess the effect of large library differences on the CMS classifications. We also identified CMS using the RUV-III normalized data (Supplementary Figure 6 A and B). We found a high concordance between the CMS obtained using the different strategies and datasets normalized by different methods (Supplementary Figure 6 C). The CMS obtained from the FPKM.UQ normalized data were used to create pseudo-samples for RUV-III normalization

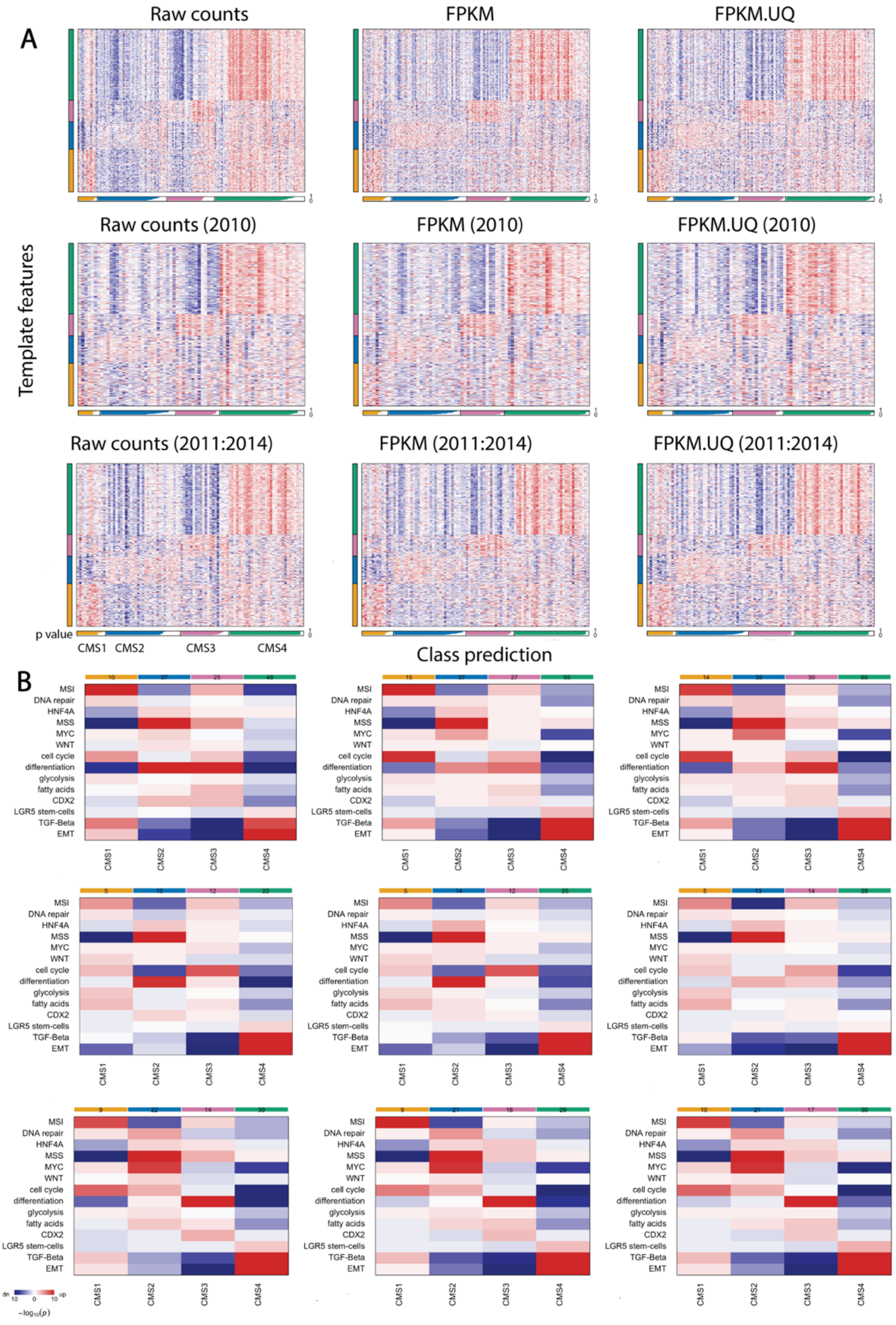


**Supplementary Figure 5. Identification of consensus molecular subtypes (CMS) in the TCGA READ RNA-seq studies. A)** Heatmaps show the relative expression levels of CMS marker genes (vertical bar) with classifications indicated below (horizontal bar, white indicating prediction confidence p-values). **B)** Heatmaps show the results obtained from gene set enrichment analysis (Camera), confirming enrichment of known characteristics in each CMS group. Heatmaps show statistical significance and red and blue indicates direction of change.

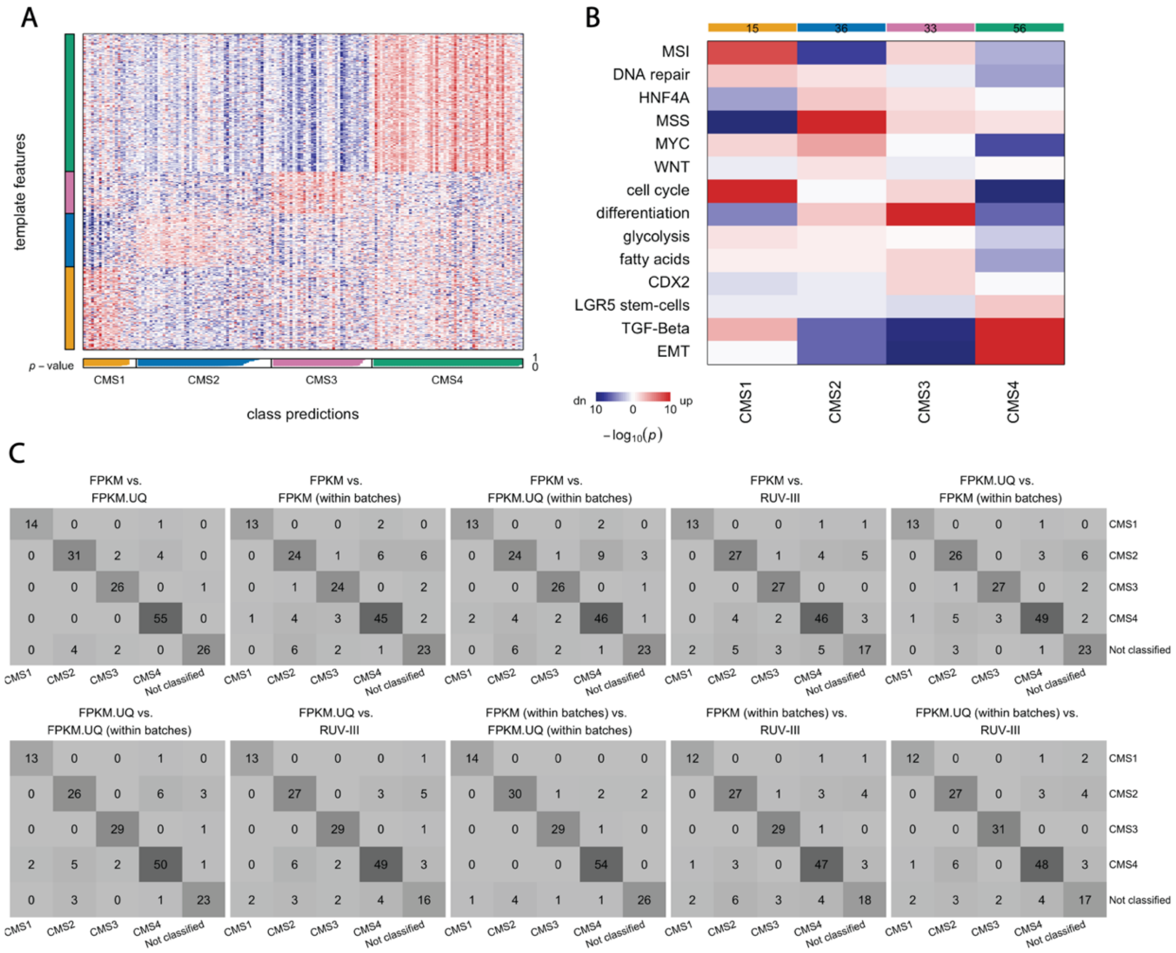


**Supplementary Figure 6. Identification of consensus molecular subtypes (CMS) in the RUV-III normalized data. A)** Heatmap shows the relative expression levels of CMS marker genes (vertical bar) with classifications indicated below (horizontal bar, white indicating prediction confidence p-values). **B)** Heatmap shows the results obtained from gene set enrichment analysis (Camera), confirming enrichment of known characteristics in each CMS group. Heatmaps show statistical significance and red and blue colours indicates direction of change. **C)** The tables show CMS calls obtained from different strategies and datasets normalized by different methods.

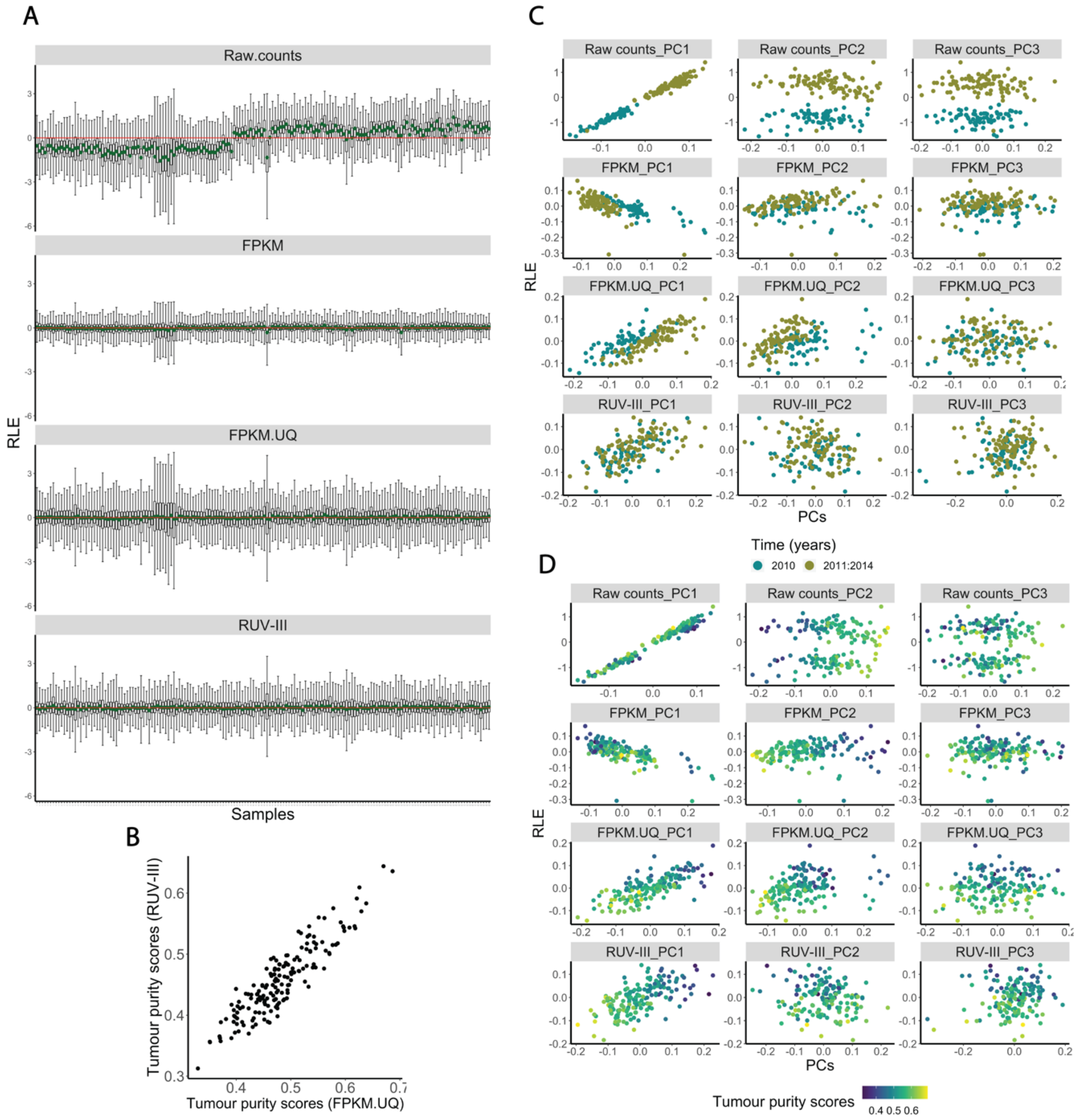


**Supplementary Figure 7. RLE plots, tumour purity estimates and relationships between the RLE plot medians and principal components in the TCGA READ RNA-sequencing data. A)** RLE plots of raw counts, TCGA and RUV-III normalized data. **B)** Scatter plot shows tumour purity scores obtained from the TCGA FPKM.UQ and RUV-III normalized data. Note, no attempt was made to remove tumour purity variation by RUV-III normalization**. C)** The scatter plots show the relationships between the first three principal components and the RLE plot medians of raw counts, TCGA and RUV-III normalized data coloured by key time intervals. **D)** The same as C, coloured by the tumour purity scores.

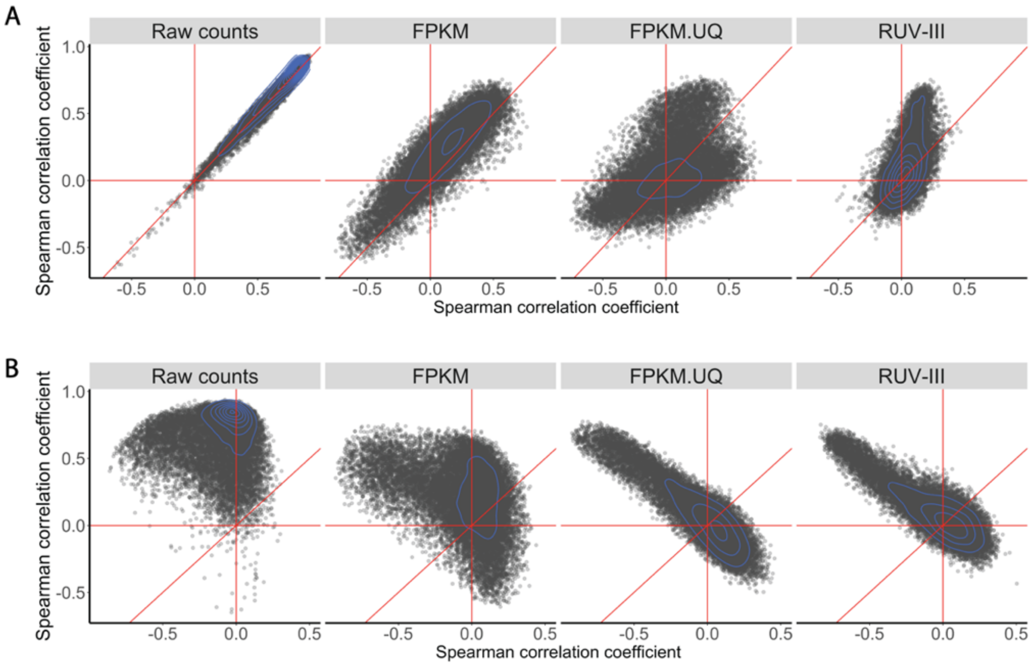


**Supplementary Figure 8. Spearman correlation analysis between individual gene expression values and the RLE plot medians, library size, and tumour purity in the TCGA READ RNA-seq datasets. A)** Scatter plots show Spearman correlation coefficients between individual gene expression levels and the RLE plot medians (Y axis) and library sizes (X axis) in the different datasets. **B)** Scatter plots display Spearman correlation coefficients between individual gene expression values and RLE plot medians (Y axis), and tumour purity (X axis) in the different datasets. The diagonal lines are the 45-degree line.

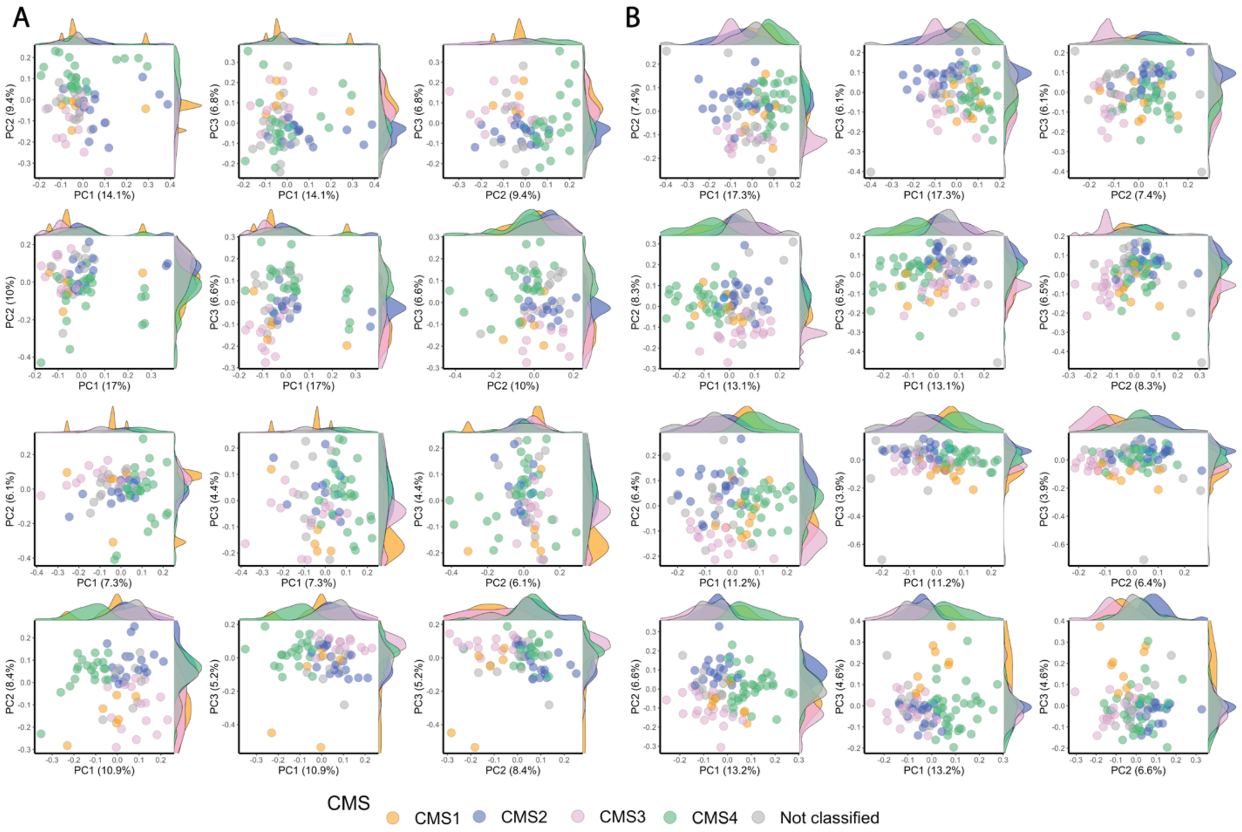


**Supplementary Figure 9. PCA plots coloured by CMS with each key time interval in the TCGA READ RNA-seq data normalized by different methods. A)** PCA plots of different datasets (from top to bottom: raw counts, FPKM, FPKM.UQ and RUV-III normalized data for samples profiled in 2010. **B)** Same as A, for samples profiled in 2011-2014. The CMS were obtained separately within each time interval.

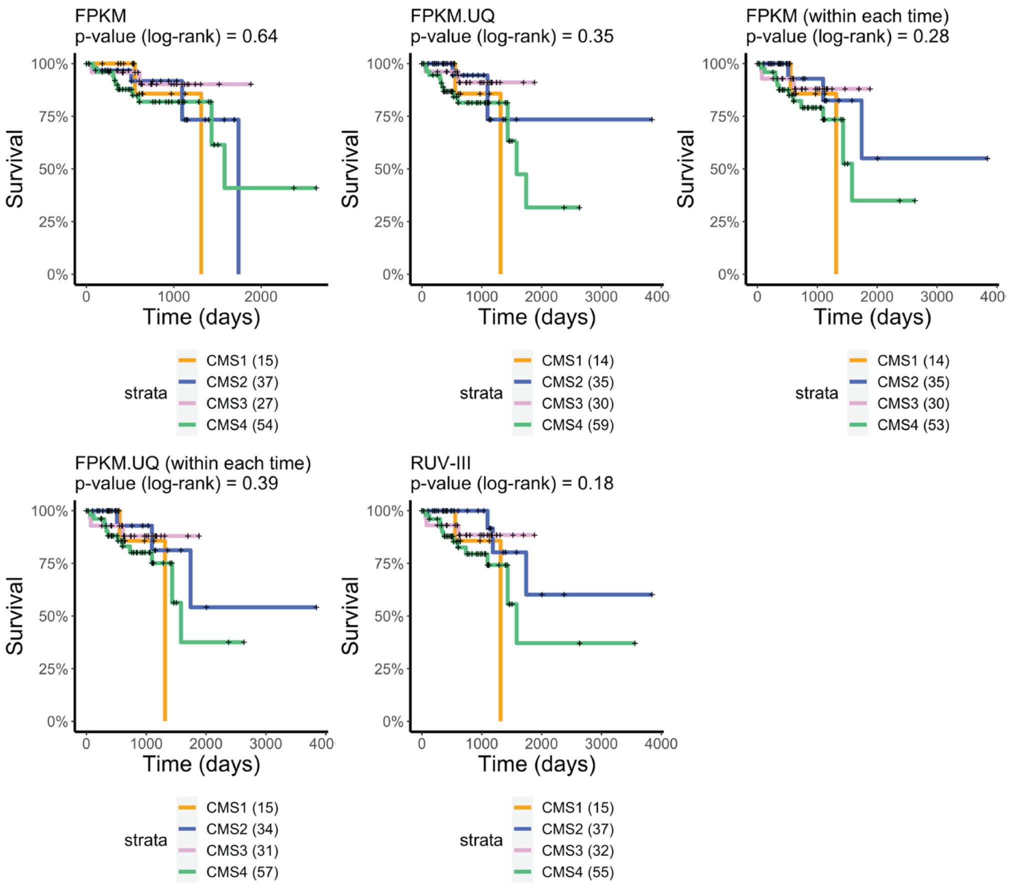


**Supplementary Figure 10. Kaplan-Meier survival curves by CMS using the TCGA READ RNA-seq data normalized by different methods.** Prognostic value of CMS with Kaplan-Meier survival analysis in FPKM, FPK.UQ and RUV-III normalized data for overall survival. The CMS for FPKM and FPKM.UQ data were identified either in the full set of samples or within each key time interval**.**

### TCGA COAD RNA-seq study

#### *Study outline*

The TCGA colon adenocarcinoma (COAD) RNA-seq study consists of 479 assays generated across 4 years (Supplementary Figure 11). There are 40 adjacent normal colon tissues that were profiled after 2010. The median library size of samples assayed in 2010 is less than a half that of the rest of the samples (Supplementary Figure 11). The consensus molecular subtypes (CMS) of these samples were identified using the R package CMScaller [2] on the raw counts and differently normalized datasets (Supplementary Figure 12 and 13).

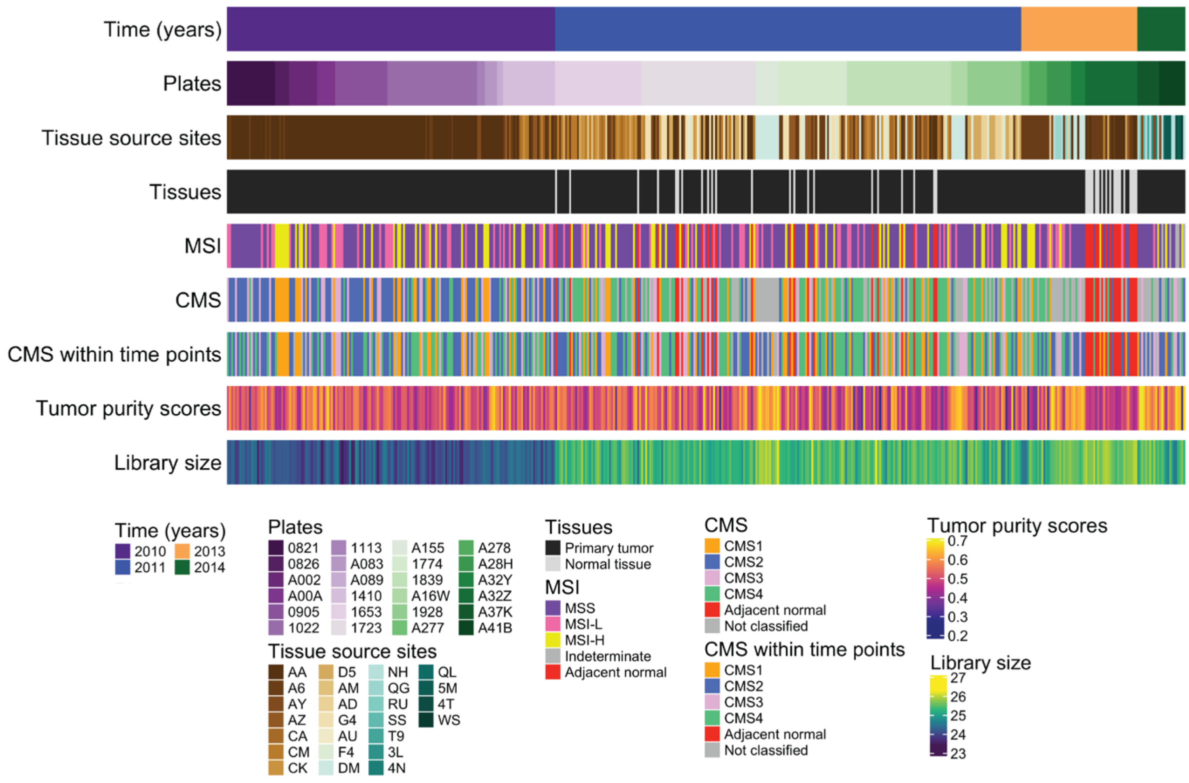


**Supplementary Figure 11. Outline of the TCGA colon adenocarcinoma (COAD) RNA-seq study. A)** 479 colon adenocarcinoma and adjacent normal tissues were collected from 25 tissue source sites (TSS), and distributed to 24 sequencing-plates for profiling at 20 different times over a span of 4 years. The microsatellite instability (MSI) and tumour stage assignments were obtained from the TCGA pathological reports. The consensus molecular subtypes (CMS) were identified using the CMScaller R package on the FPKM.UQ normalized data in two different ways, across all samples and within key time intervals (2010 and 2010-2014). The sample library sizes are calculated after removing lowly expressed genes and are $log_2$ transformed. The tumour purity scores are obtained from 1- stromal and immune scores (see Methods).

### The consensus molecular subtypes of the TCGA COAD RNA-seq samples

As with the TCGA READ RNA-seq data, we used the R package CMScaller to identify the consensus molecular subtypes (CMS) of the TCGA COAD samples. We applied the classifier to the raw counts and the FPKM, FPKM.UQ, and RUV-III normalized datasets (Supplementary Figure 12 and 13 A and B). We also applied the classifier within each key time interval, 2010 and 2011-2014, in the TCGA normalized datasets to assess the effects of the large library size differences between the time intervals on the CMS classifications. The results showed that the proportion of un-classified samples was significantly smaller when the classifier was applied within each key time interval compared with that when applied across all samples together (Supplementary Figure 13 C and D). Supplementary Figure 13 E shows that the CMS obtained using the full set of the FPKM and FPKM.UQ data are associated with library size. We also found the CMS identified in the RUV-III normalized data are highly concordant with the CMS obtained using the TCGA normalized datasets within each key time point. These results show the effect of library size on the identification of CMS in the full sets of FPKM and FPKM.UQ normalized data.

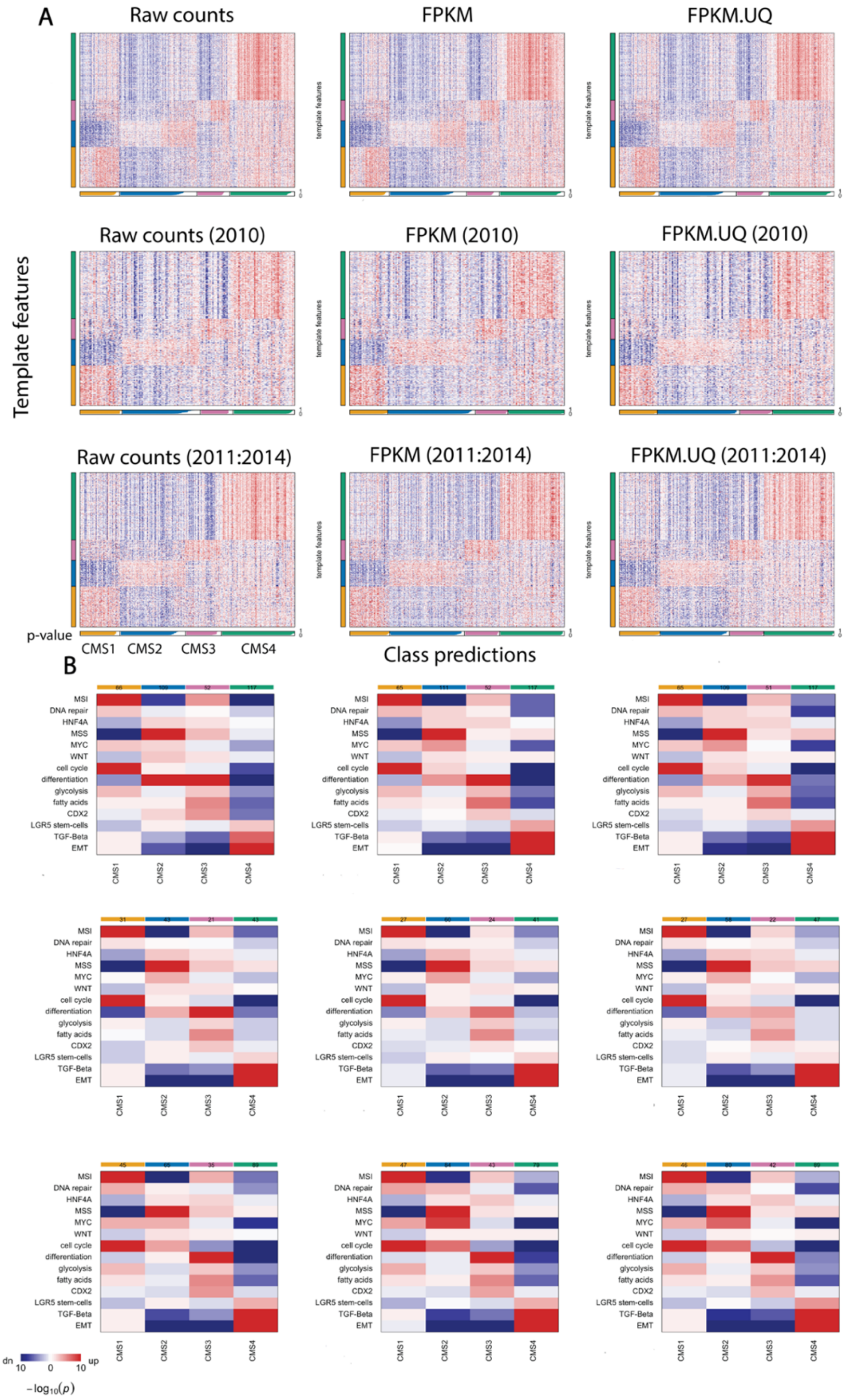


**Supplementary Figure 12. Identification of consensus molecular subtypes in the TCGA COAD RNA-seq data. A)** Heatmaps show the relative expression levels of subtype marker genes (vertical bar) with classifications. CMS1-CMS4, designated below (horizontal bar, white colour display prediction confidence p-values). The CMS classification were performed across all samples and within each key time intervals (2010 and 2010-2014) to assess the impact of batch effects. **B)** Heatmaps show the results from gene set analysis, confirming enrichment of known characteristics in each CMS group. Heatmaps show statistical significance and red and blue colours indicates direction of change.

It has been shown that different CMS are associated with various gene signatures [2]. Gene set expression analyses show that the CMS obtained from the FPKM, FPKM.UQ and RUV-III normalization are associated with the known gene signatures. (Supplementary Figure 12, B and 13, B). These results suggested that the different normalizations preserve known biological signals associated with CMS in the data.

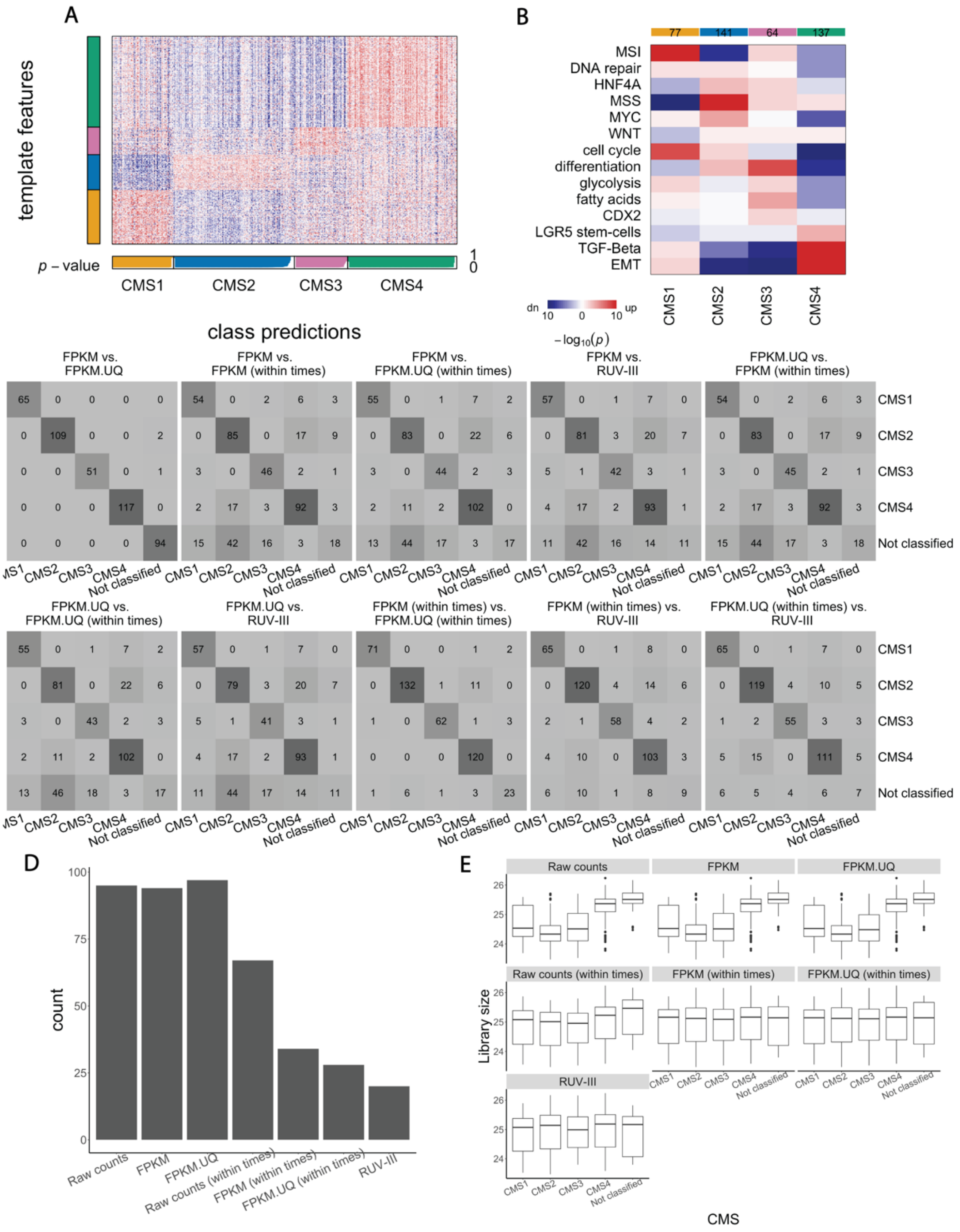

**Supplementary figure 13. Identification of consensus molecular subtypes (CMS) in the TCGA COAD RNA-sequencing study using the RUV-III normalized data and comparing CMS obtained after normalizing by different methods. A)** Heatmap shows the relative expression levels of subtype marker genes (vertical bar) with classifications. CMS1-CMS4, designated below (horizontal bar, white colour display prediction confidence p-values). **B)** Heatmap shows the results from gene set analysis, confirming enrichment of known characteristics in each CMS group. Heatmaps show statistical significance and red and blue colours indicates direction of change**. C)** Comparisons between CMS calls obtained from datasets from different normalization and conditions. **D)** Bar plot shows the number of un-classified samples obtained by CMScaller in different datasets. **E)** Plots show the relationship between library size ($log_2$) and CMS calls in different datasets

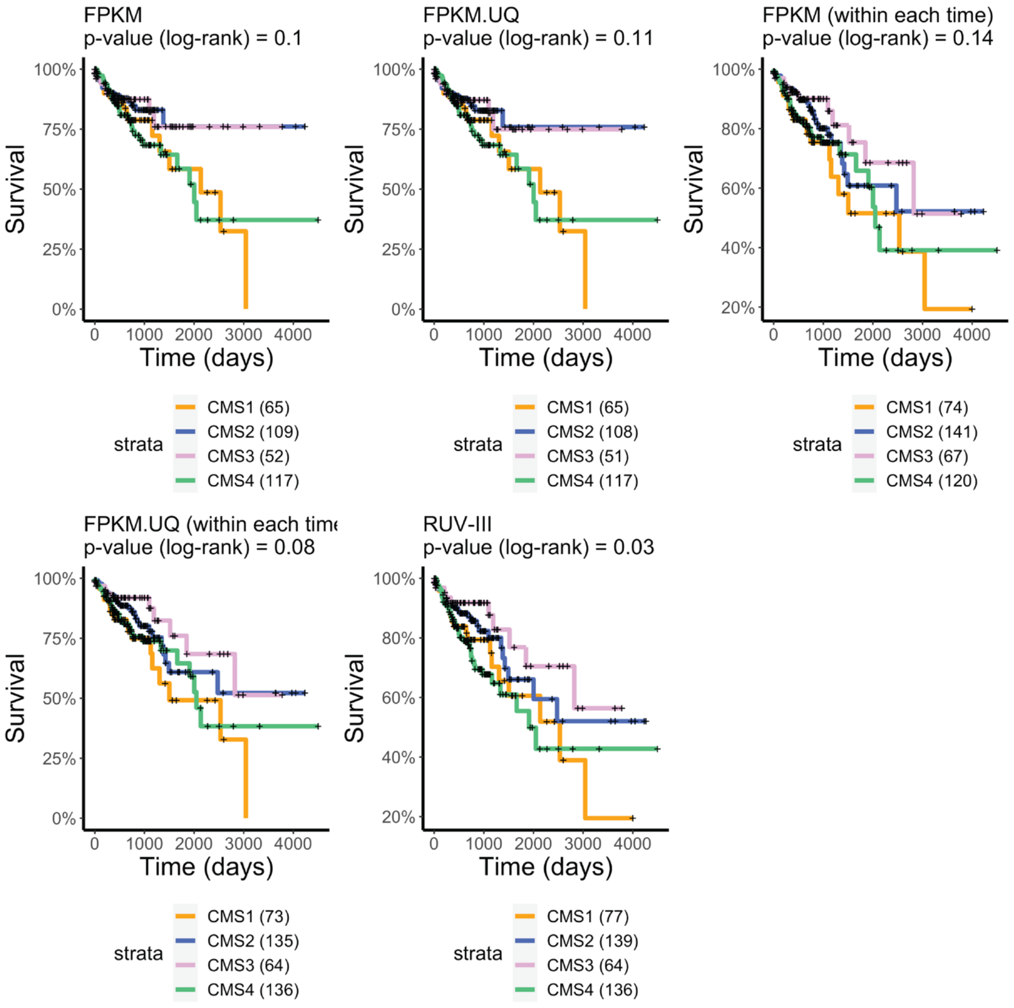


**Supplementary figure 14. Kaplan-Meier survival analyses by consensus molecular subtype in the TCGA COAD RNA-seq normalized by different methods.**

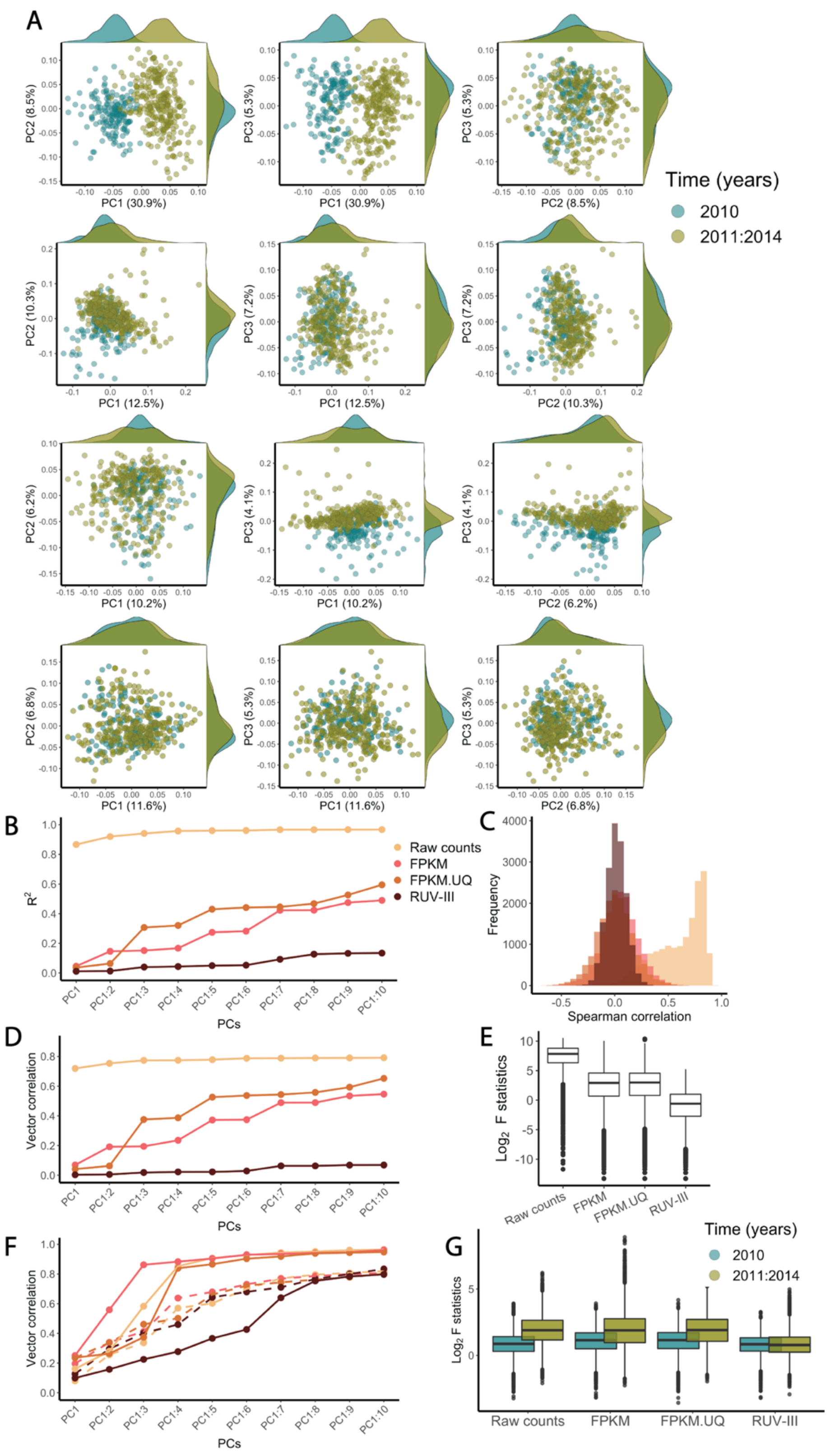


**Supplementary figure 15. Performance of different normalization methods in removing library size and plate effects in the TCGA COAD RNA-seq data. A)** From top row to bottom, Scatter plots of first three principal components for the raw counts, FPKM, FPKM.UQ and RUV-III normalized data coloured by key time intervals (2010 , 2011-2014). **B)** A plot showing the R-squared ($R^2$) of linear regression between library size and up to the first 10 principal components (cumulatively) of different datasets. **C)** Histograms of Spearman correlation coefficients between the gene expression levels and library size. **D)** A plot showing the vector correlation between the key time intervals and up to the first 10 principal components (cumulatively). **E)** Boxplots of $log_2$ F statistics obtained from ANOVA between gene expression with key time intervals being the factor. **F)** A plot showing the vector correlation between plates and up to the first 10 principal components (cumulatively) within each key time interval. **G)** Boxplots of $log_2$ F statistics obtained from ANOVA between gene expression and plates within key time intervals.

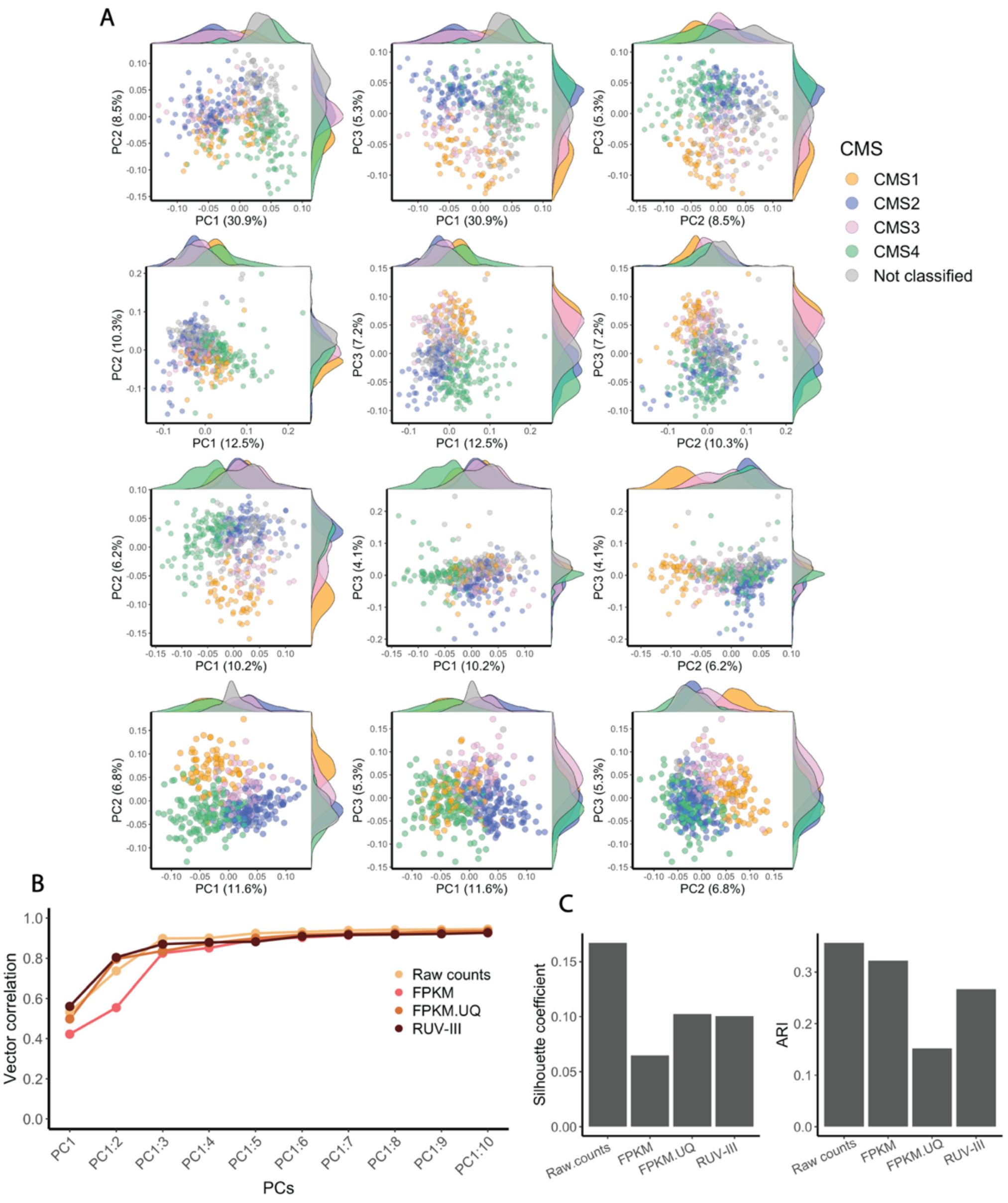


**Supplementary Figure 16. Performance of the different normalization methods in preserving the consensus molecular subtypes in the TCGA COAD RNA-seq data. A)** From top row to bottom: scatter plots of the first three principal components colored by CMS for the TCGA COAD raw counts, FPKM, FPKM.UQ and RUV-III normalized data. The CMS were obtained for each dataset separately. **B)** A plot showing the vector correlation between CMS and up to the first 10 principal components (cumulatively) in the differently normalized datasets. **C)** Silhouette coefficients and ARI index for showing the separation of CMS subtypes in the differently normalized datasets.

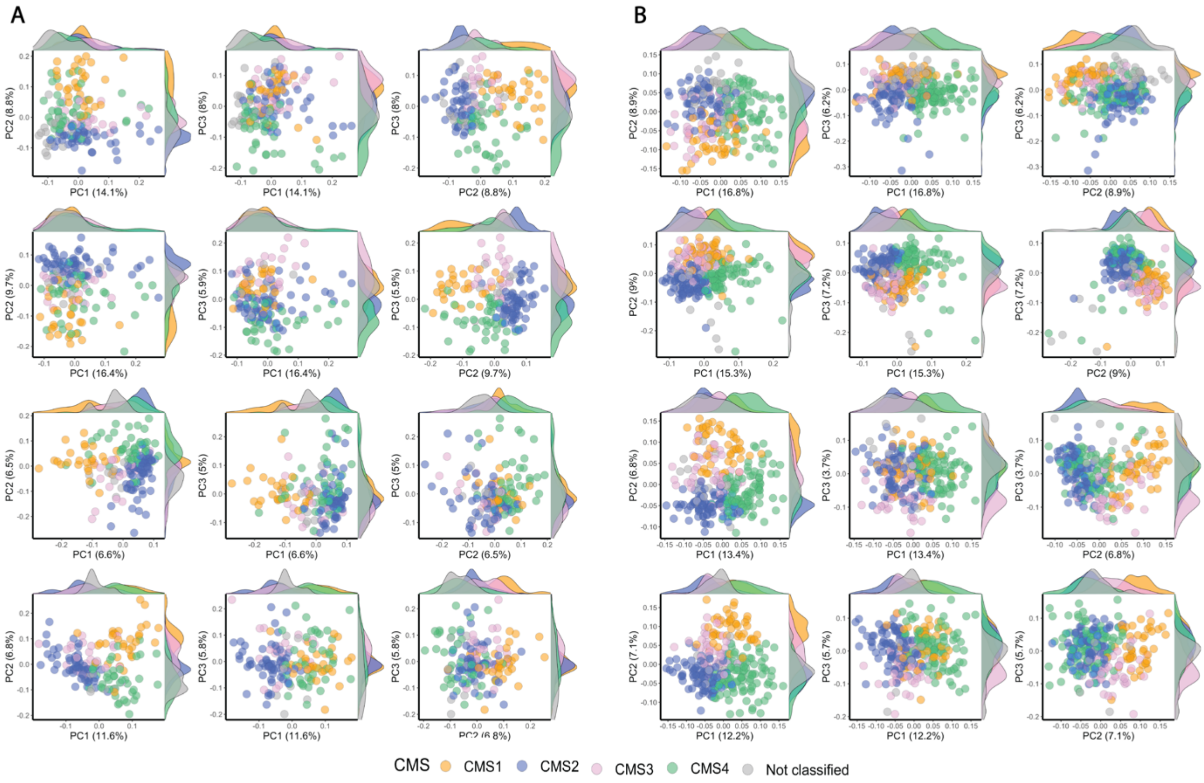


**Supplementary figure 17. PCA plots coloured by CMS within each key time interval in the TCGA COAD RNA-seq data normalized by different methods. A)** Scatter plots of the first three principal components obtained from samples profiled in 2010 (from top to bottom: the TCGA raw counts, FPKM, FPKM.UQ, and RUV-III normalized datasets). **B)** Same as A, for samples profiled in 2011-2014.

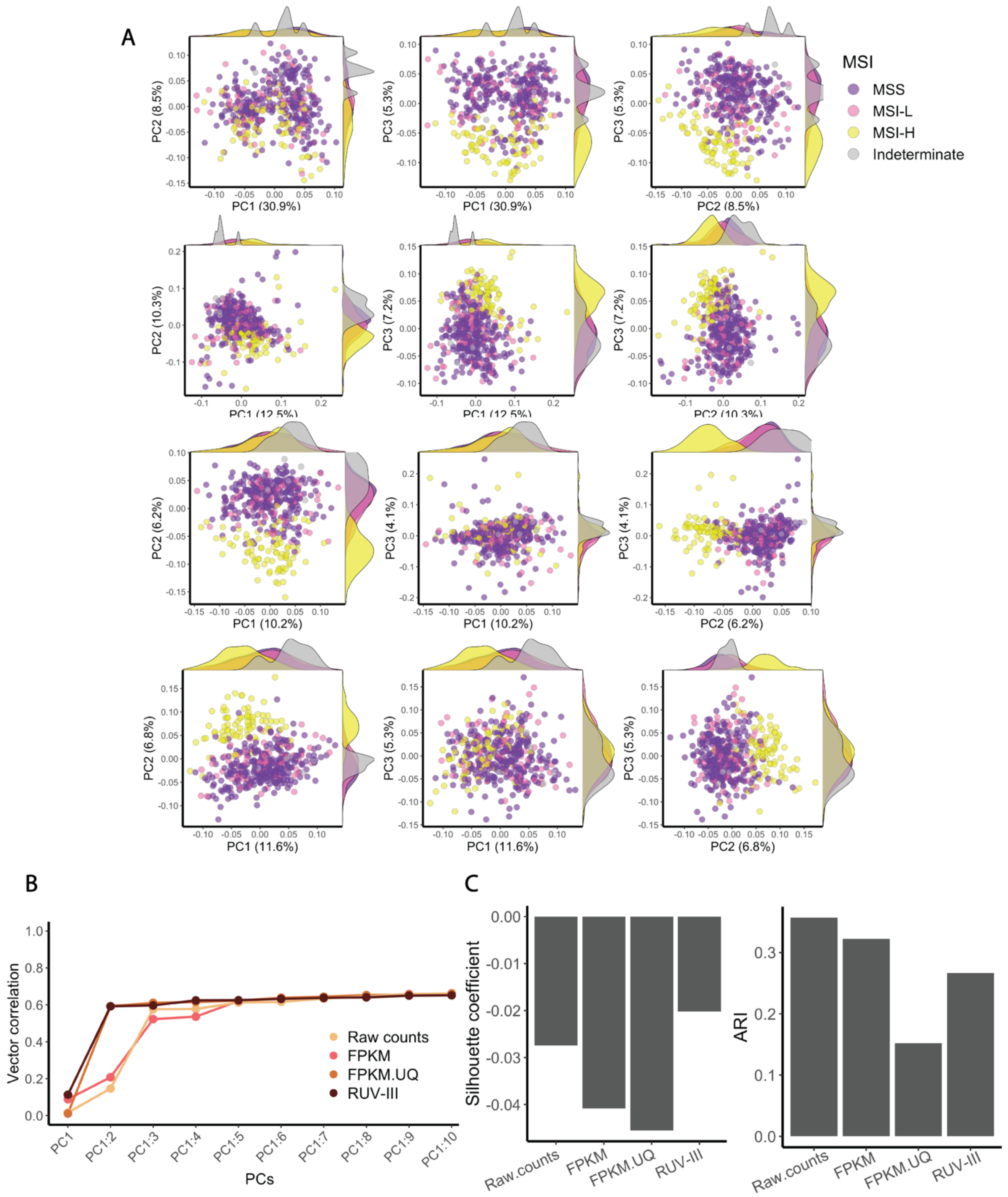


**Supplementary figure 18. Performance of different normalization methods in preserving the microsatellite instability (MSI) clusters in the TCGA COAD RNA-seq data. A)** Scatter plots of the first three principal components colored by MSI for the TCGA COAD raw counts, FPKM, FPKM.UQ and RUV-III normalized data. The MSI details (MSS: microsatellite stable, MSI-L = microsatellite instability low, MSI-H = microsatellite instability high) were obtained from the TCGA pathological data. **B)** Plot showing the vector correlation coefficient between MSI and up to the first ten principal components (cumulatively). **C)** Silhouette coefficients and ARI index for exhibiting the separation of MSI subtypes in different datasets

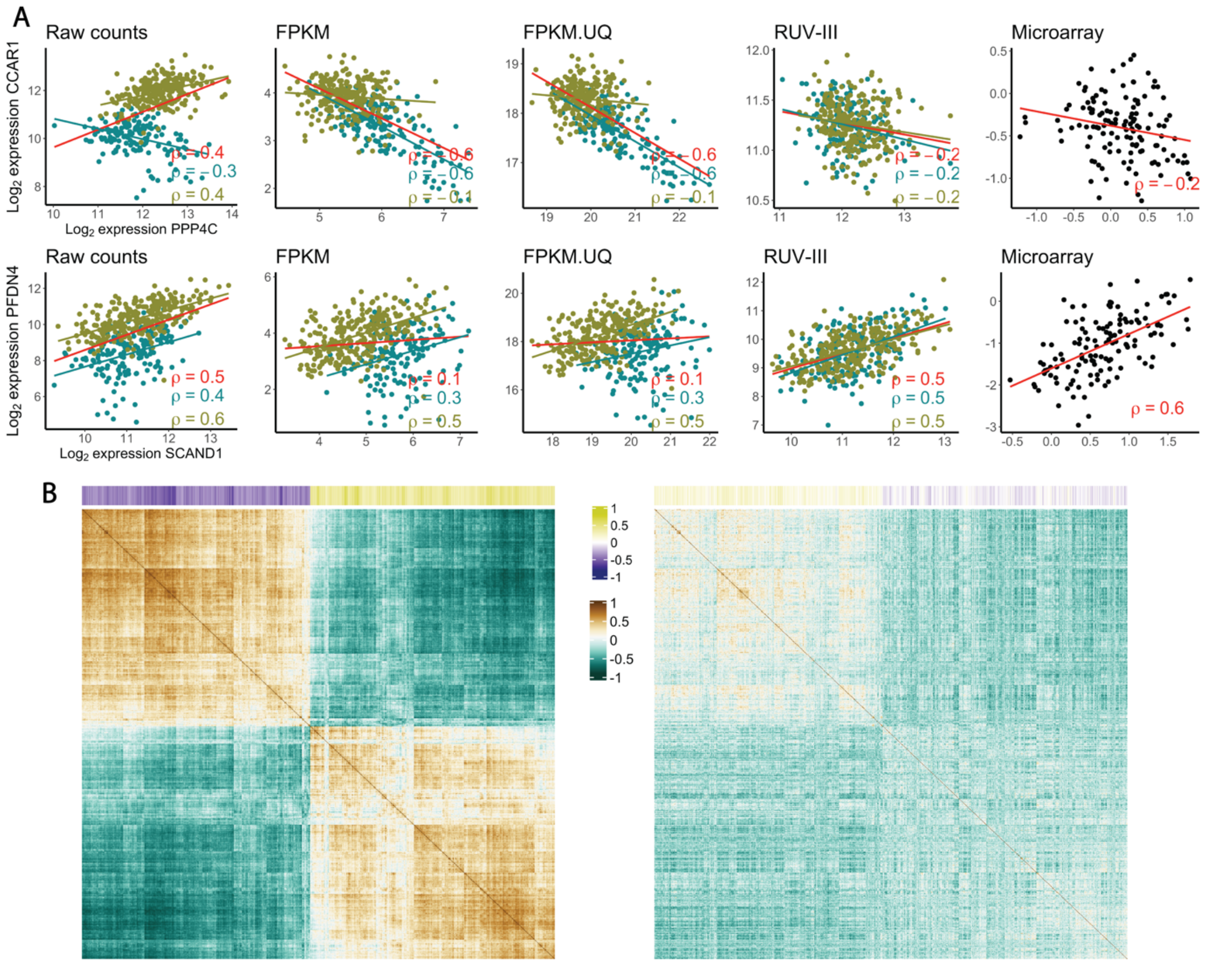


**Supplementary figure 19. Impact of unwanted variation on gene co-expression analyses in the TCGA COAD RNA-seq data. A)** First row: scatter plots between the expression levels of the *CCAR1* and *PPP4C* genes in the raw and differently normalized counts. The red line indicates overall association between the expression levels, while the green and yellow lines show the associations seen in the 2010 samples and the rest, respectively. **B)** First heatmap shows correlation matrix of the 1300 genes whose expression levels have the highest correlation with library size in the FPKM.UQ normalized data. The coloured bars at the top of heatmaps show the correlation of individual gene expression levels with library size. Second heatmap: same as the first one, for the RUV-III normalized data. The order of rows and columns in both correlation matrix are the same.

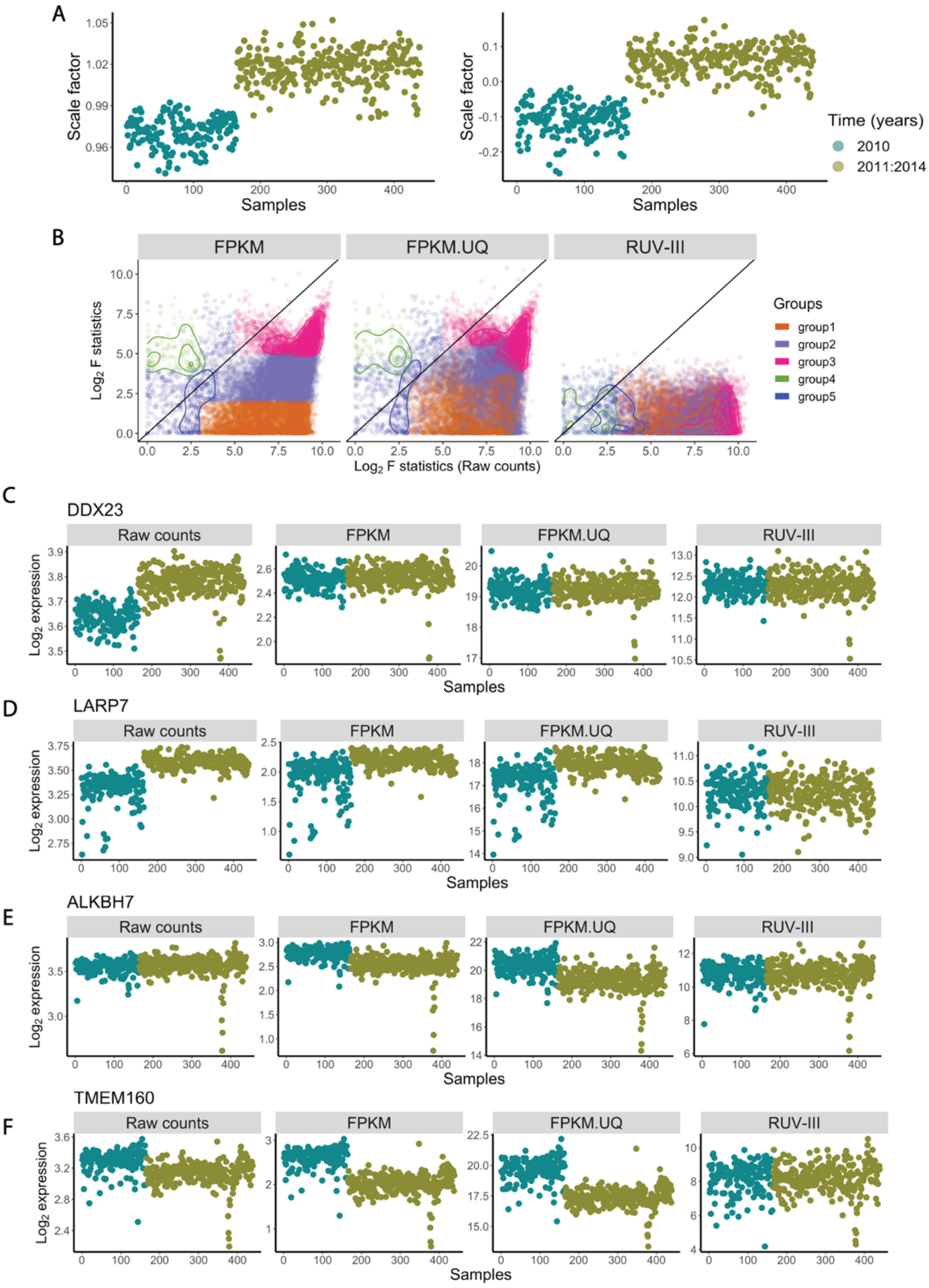


**Supplementary figure 20. Relationship between gene level ($log_2$) counts and ($log_2$) library size in the TCGA COAD RNA-seq. A)** Plots shows the global scale factors obtained by sample library sizes (first plot) and upper-quartiles (second plot) of the COAD raw counts against time. **B)** Scatter plots of the $log_2$ F-value obtained from ANOVA analyses of gene expression levels with the major time variation: 2010 vs all other years; ($log_2$) raw READ counts on the horizontal axes of all plots and differently normalized counts vertically. **C)** Expression patterns of four genes whose counts have different relationships with the global scaling factors calculated from the COAD raw count data.

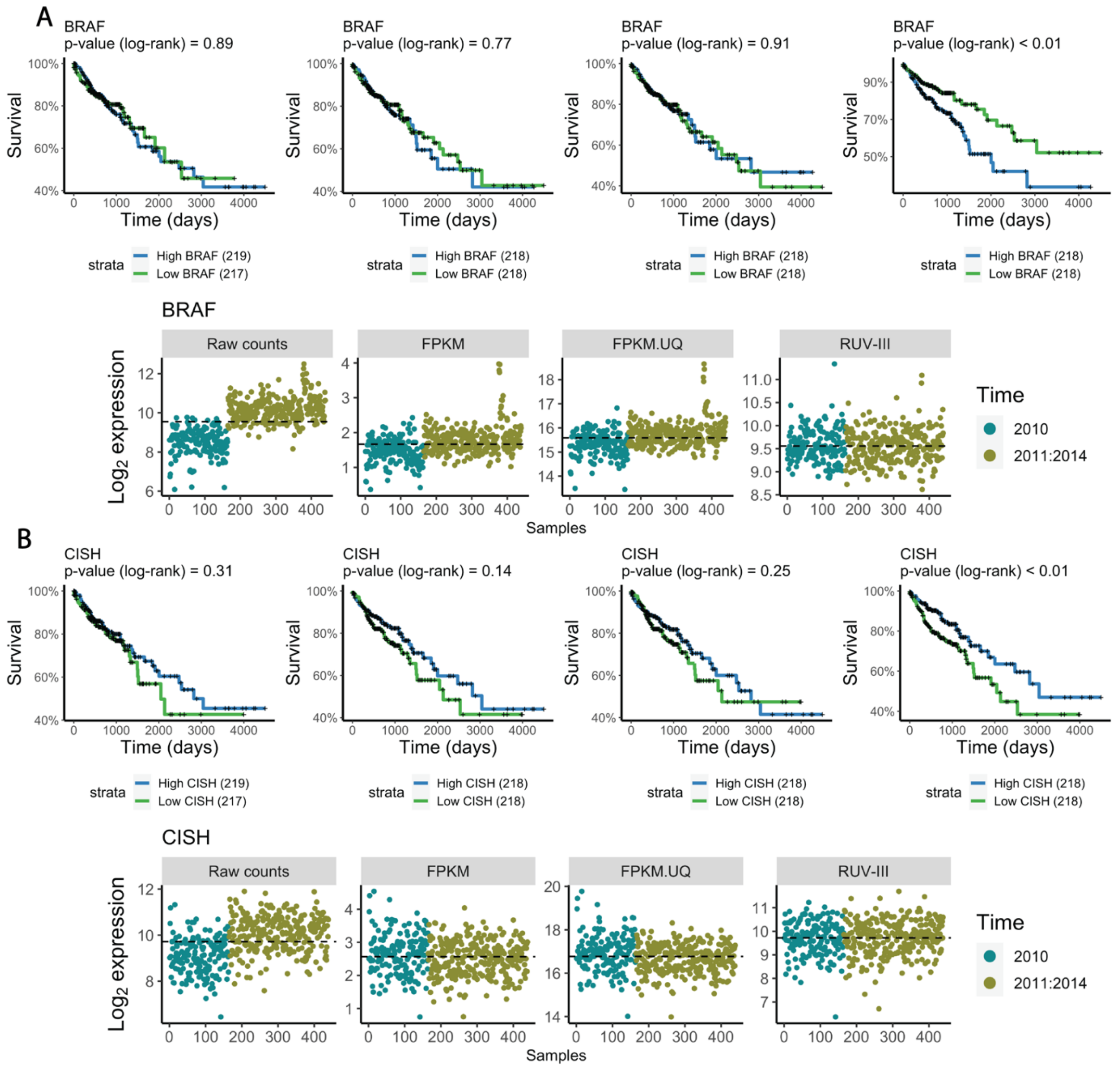


**Supplementary figure 21. Association of the BRAF and CISH gene expression levels with overall survival in the differently normalized TCGA COAD datasets. A)** First row: Kaplan Meier survival analysis shows the association between the BRAF gene expression and overall survival in differently normalized TCGA COAD datasets. Second row: expression patterns of the BRAF gene expression across samples in differently normalized TCGA COAD datasets. **B)** same as A, for the CISH gene**.**

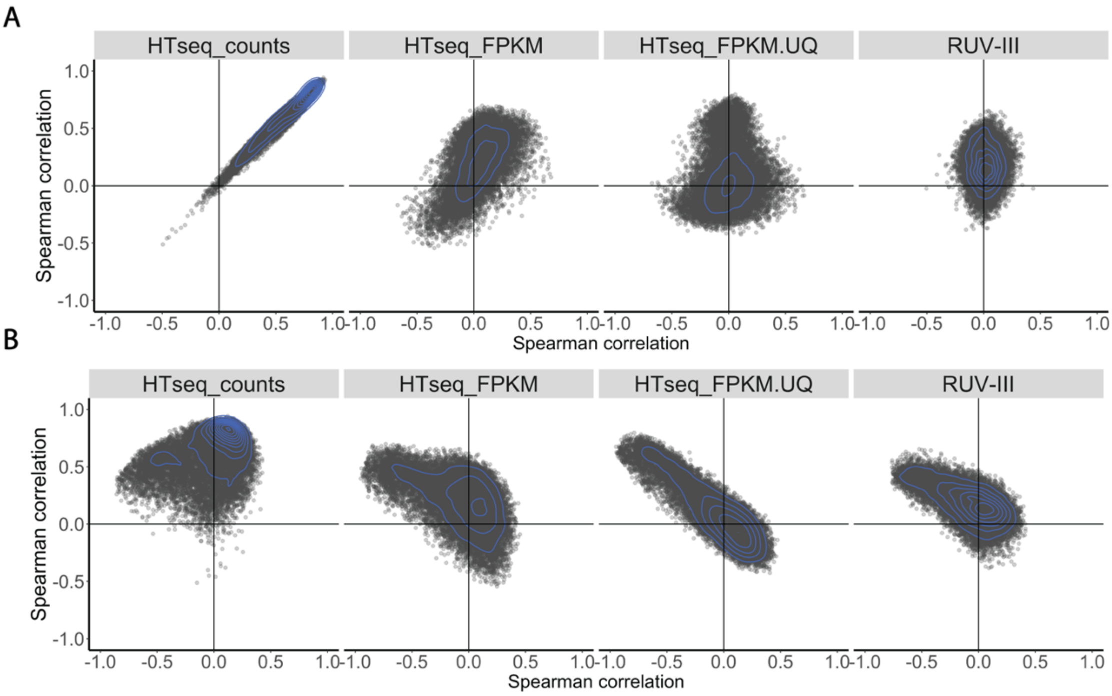


**Supplementary figure 22. Spearman correlation analyses between gene expression levels and RLE plot medians, library size, and tumour purity in the TCGA COAD raw and differently normalized counts. A)** Scatter plots of Spearman correlations between individual gene expression levels with RLE plot medians (Y axis), and library size (X axis) in the different datasets. **B)** Scatter plot of Spearman correlation coefficients between individual gene expression levels with RLE plot medians (Y axis), and tumour purity (X axis) in the different datasets.

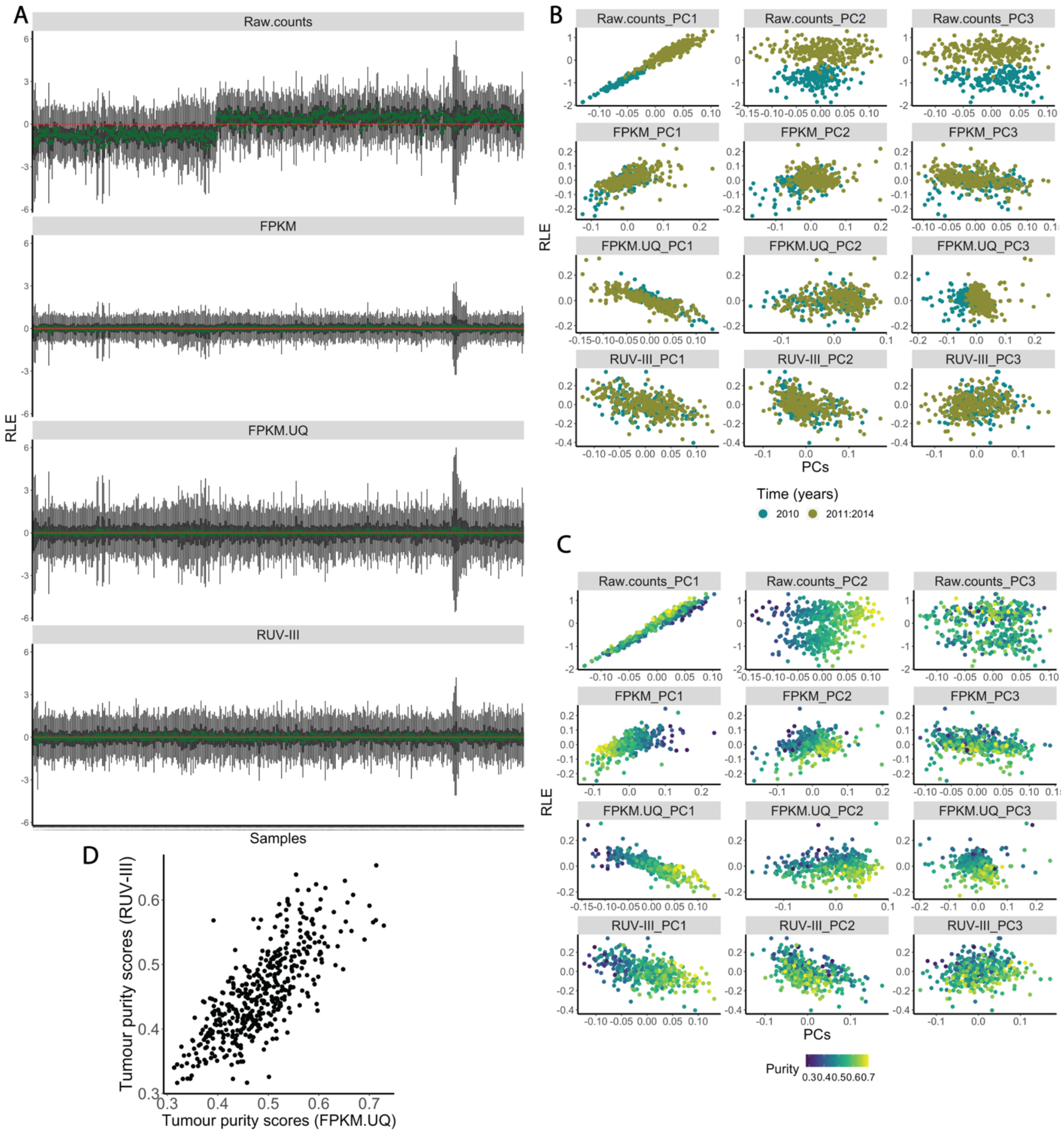


**Supplementary figure 23. RLE plots and relationships between the RLE plot medians and principal components in the raw and differently normalized counts of the TCGA COAD RNA-seq data. A)** RLE plots of raw counts, TCGA and RUV-III normalized datasets. **B)** Scatter plots show the relationships between the first three principal components and the RLE plot medians for raw and differently normalized counts, coloured by different times. **C)** As for B, coloured by tumour purity. **D)** Tumour purity estimates for the TCGA FPKM.UQ and RUV-III normalized data. Note, the RUV-III normalization did not aim to remove the effect of tumour purity variation.

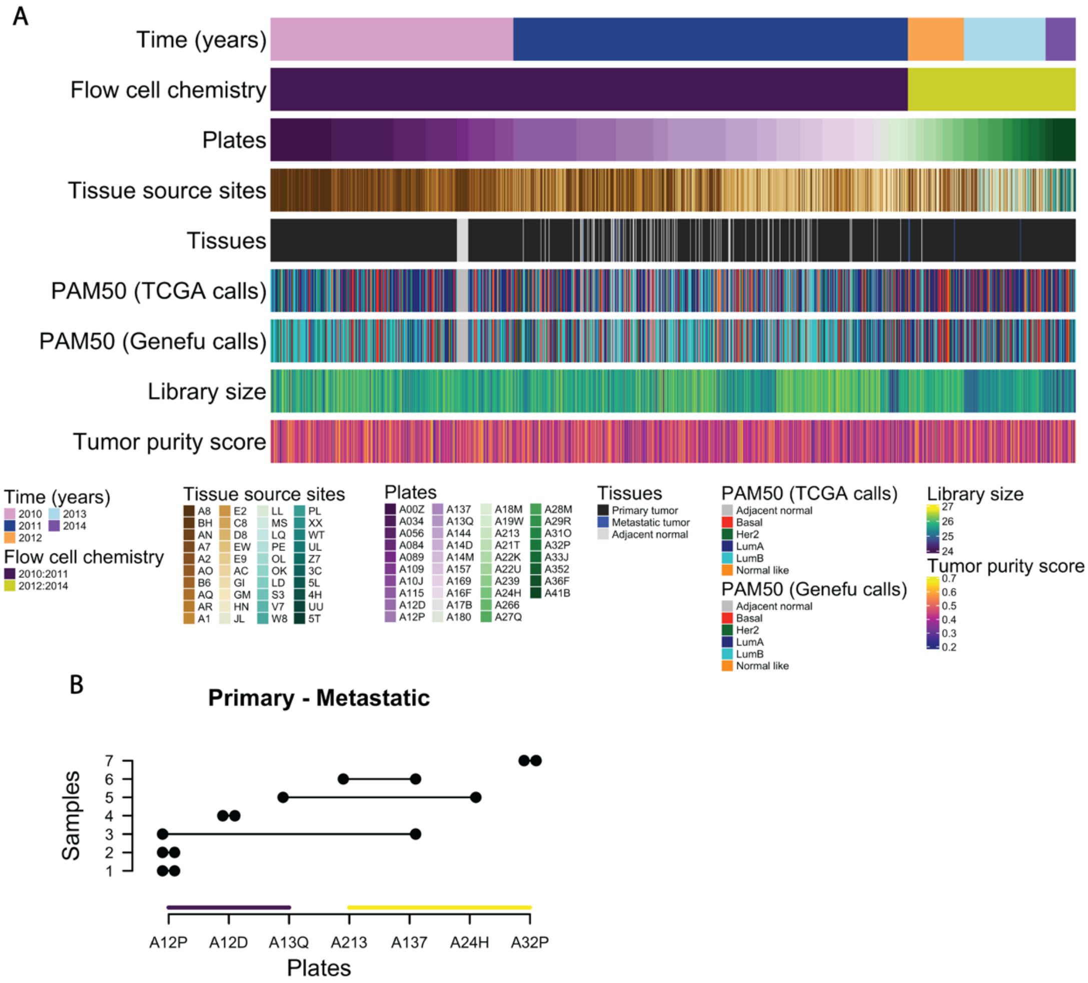


**Supplementary figure 24. a)** 1180 cancer and adjacent normal breast tissues were collected from 40 tissue source sites (TSS) and distributed unevenly to 38 sequencing-plates for profiling at 40 different time points over a span of 5 years from 2010 to 2014. Most samples (958 out of 1212 samples) were profiled in the period 2010-2011 using the first flow cell chemistry, and the rest of the samples were profiled in 2012-2014 using another flow cell chemistry. PAM50 subtypes were called using two different approaches. **b)** The distributions of 7 paired primary-metastatic samples across different sequencing-plates and the two flow cell chemistries. Each two solid points connected by a line shows a primary and metastatic sample pair**.**

### Breast tumour intrinsic subtypes

Tumour intrinsic molecular subtypes given by the prediction analysis of microarray 50 (PAM50) approach were established to better understand the biology and improve clinical outcomes of breast cancer [3]. The intrinsic subtypes of breast cancer are basal-like (Basal), human epidermal growth factor receptor 2 (HER2)–enriched, luminal A (LumA), luminal B (LumB) and normal like. The function *molecular.subtyping* in the genefu R/Bioconductor package [4] and a classifier reported by Picornell et.al [5] were used identify the PAM50 subtypes in the TCGA RNA-sequencing datasets.

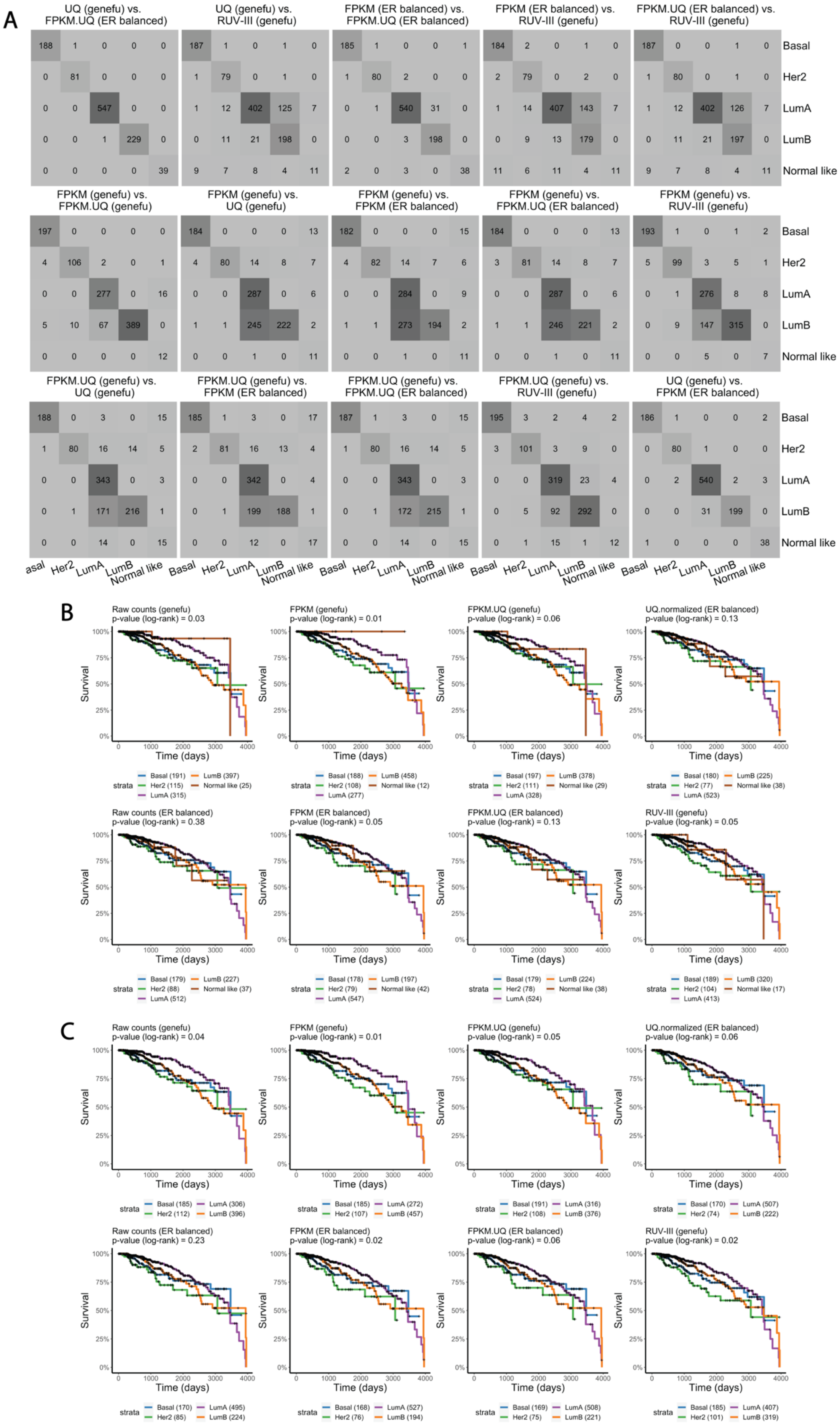

**Supplementary Figure 25. PAM50 subtype identification using two different methods in the TCGA BRCA RNA-sequencing data. A)** Tables show the concordance between the PAM50 subtypes obtained using two methods, on different datasets. UQ is upper-quartile normalization of the TCGA BRCA raw counts data. **B)** Kaplan-Meier survival curves show the association between the PAM50 subtypes and overall survival in the TCGA BRCA RNA-sequencing data. **C)** same as B, without the normal like subtypes.

This method was used on both ER-balanced and ordinary gene expression datasets normalized by differed methods. To obtain ER-balanced data, we first selected a subset of cancer samples which are ER status-balanced (the same sample sizes for estrogen receptor positive and negative values based on clinical measurements) and used the median derived from that subset to centre the entire cohort. Figure 25 A shows the concordance between the PAM50 subtypes identified by the two methods in the different datasets. The PAM50 subtypes called by the genefu on the TCGA FPKM and FPKM.UQ normalized data were used for PRPS.

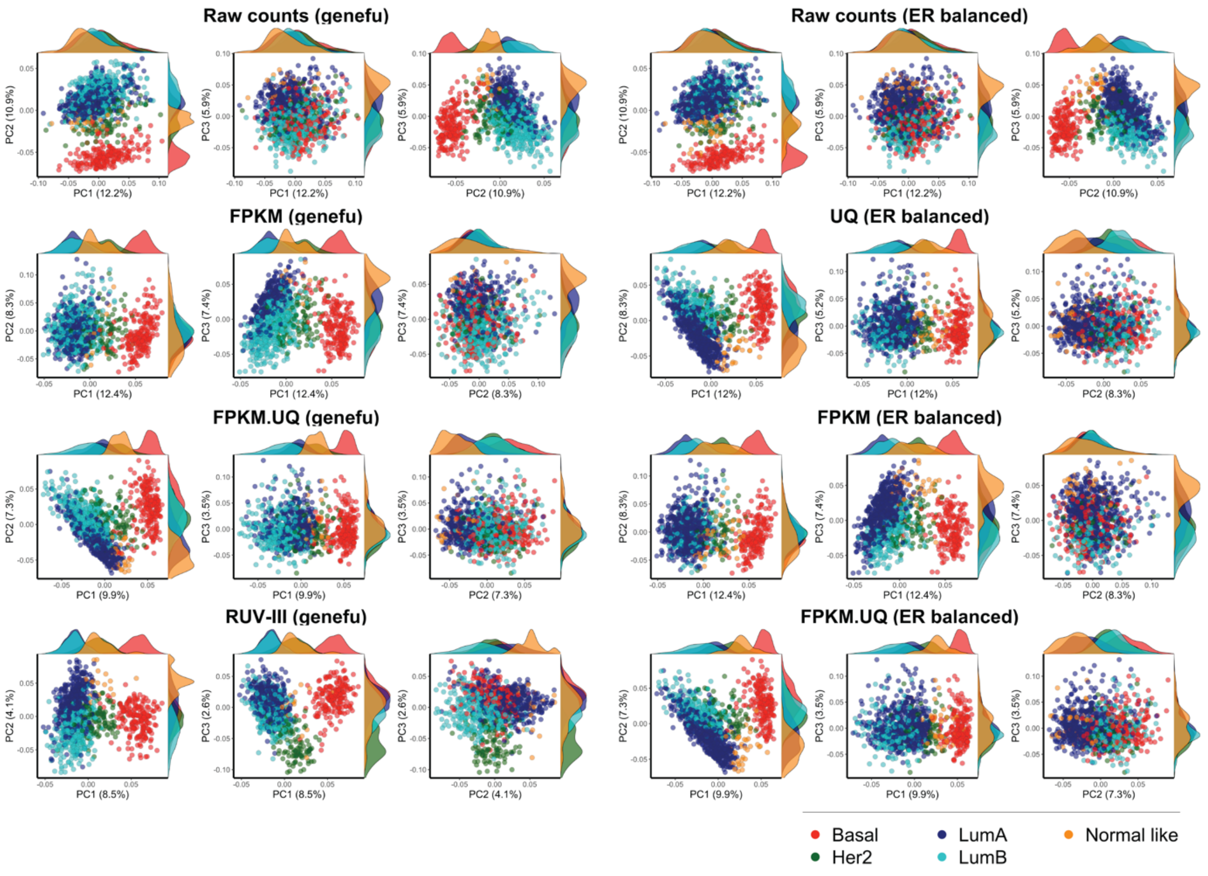


**Supplementary Figure 26. PCA plots coloured the PAM50 subtypes of the TCGA BRCA RNA-sequencing datasets**. Each panel shows the scatter plots for two of the first three PC for the raw data and several normalized versions of the TCGA BRCA RNA-sequencing data. Samples are coloured based on the PAM50 calls obtained using the different approaches. UQ is upper-quartile normalization of the TCGA BRCA raw counts data.

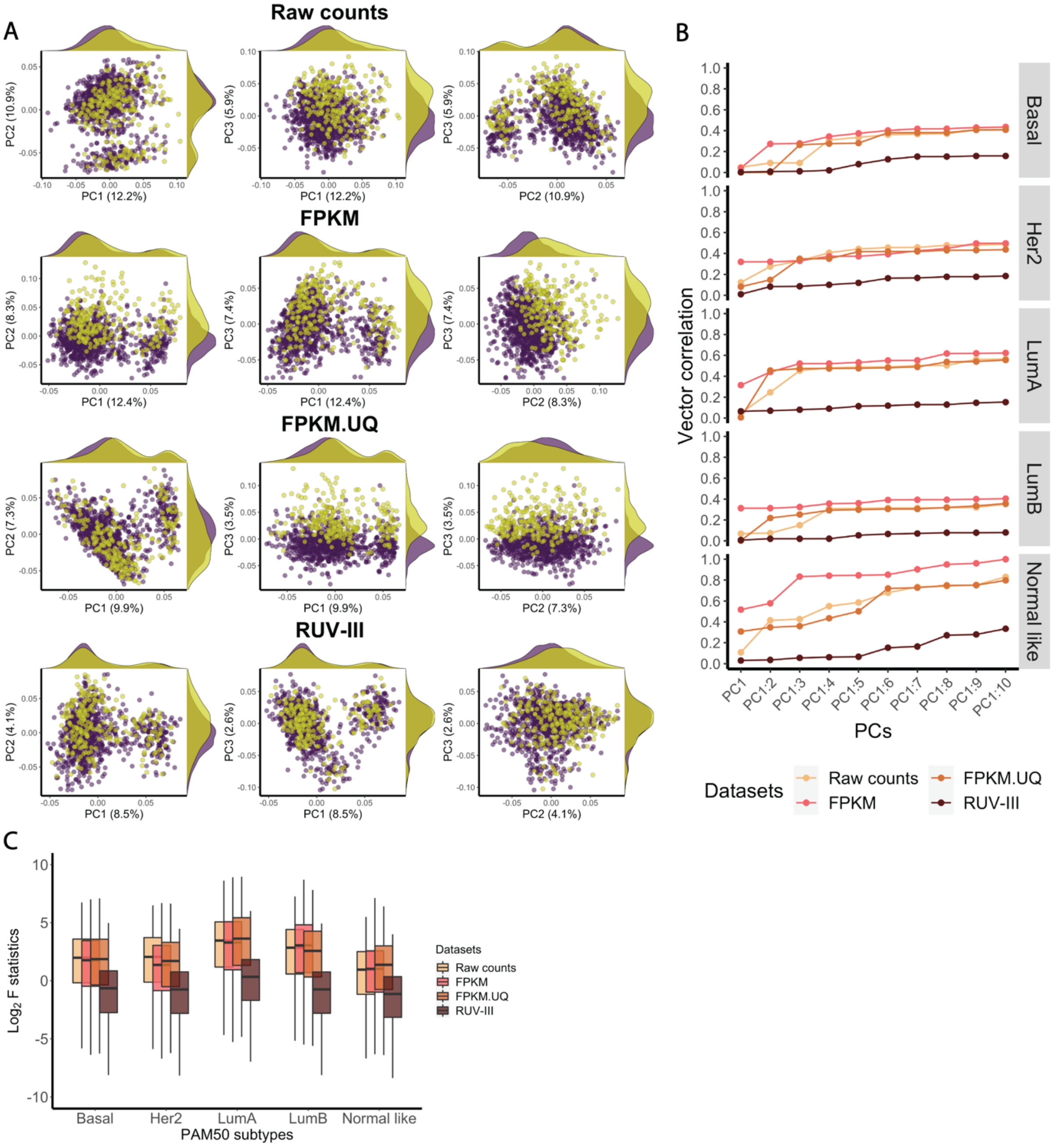


**Supplementary figure 27. RUV-III removes the effects of different flow cell chemistries in the TCGA BRCA RNA-sequencing data. A)** The first three PC coloured by flow cell chemistry in the differently normalized RNA-seq data. **B)** Vector correlation analysis between the first ten PCs cumulatively and flow cell chemistries within each of the PAM subtypes of the differently normalized TCGA BRCA RNA-seq data.

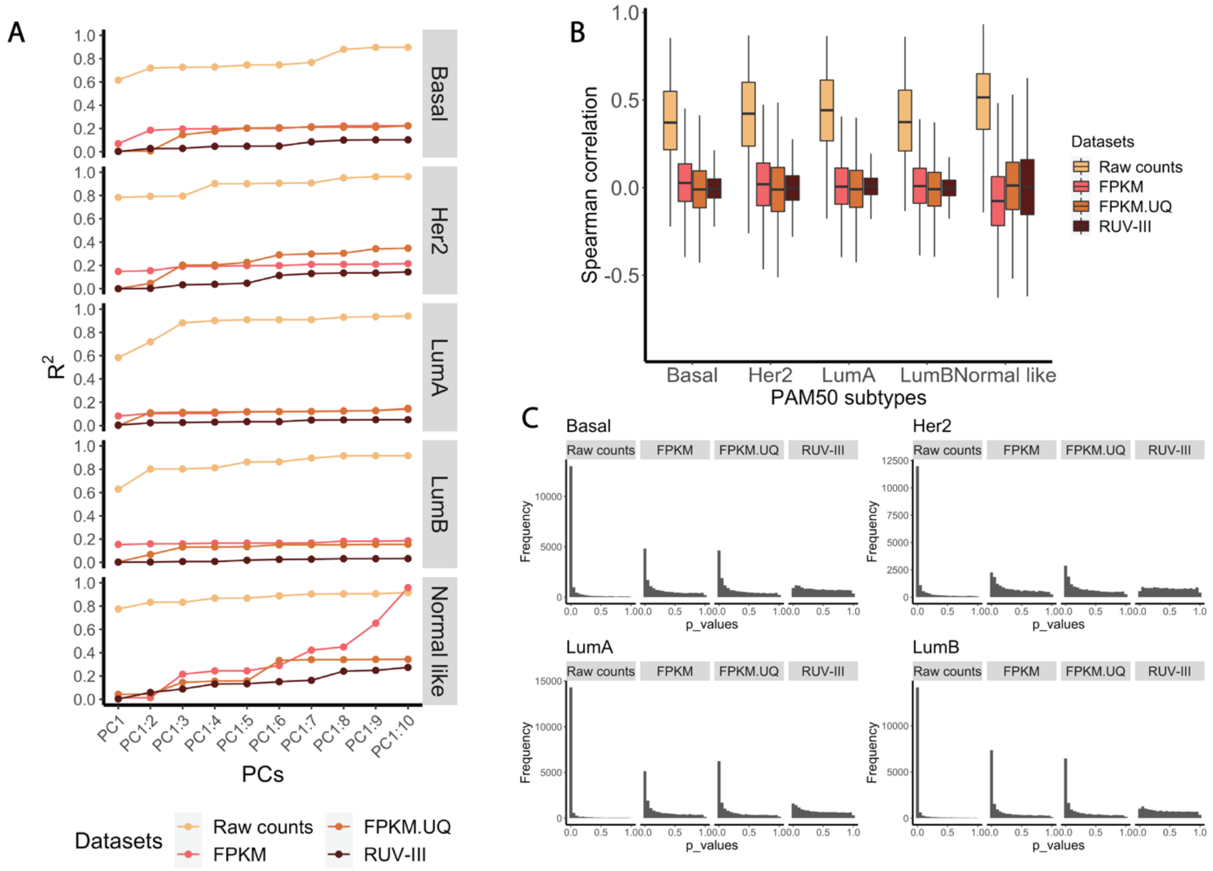


**Supplementary figure 28. RUV-III removes library size from the TCGA RNA-seq data. A)** Plots show $R^2$ calculated using linear regression between the first 10 principal components cumulatively and library size in the TCGA and RUV-III normalized datasets. **B)** The boxplots of the Spearman correlation coefficients between the individual gene expression levels and library size within the PAM50 subtypes in differently normalized datasets. **C)** The histograms of p-values obtained from differential expression analyses between samples with high and low library sizes within each PAM50 subtype in differently normalized datasets.

We also found that the chemistry effects can lead to unreliable gene expression differences between paired primary and metastatic samples assayed across the chemistries. Supplementary Figure 29 showed MA plots of the FPKM.UQ and RUV-III normalized expression values for three primary and metastatic pairs of samples. Samples from two of the pairs were profiled using the same chemistry and those from the third pair were profiled across the two chemistries. These results suggest that a considerable proportion of the gene expression differences between paired primary and metastatic samples that were profiled across chemistries likely results from the different flow cell chemistries, as we did not see this in the RUV-III normalized data.

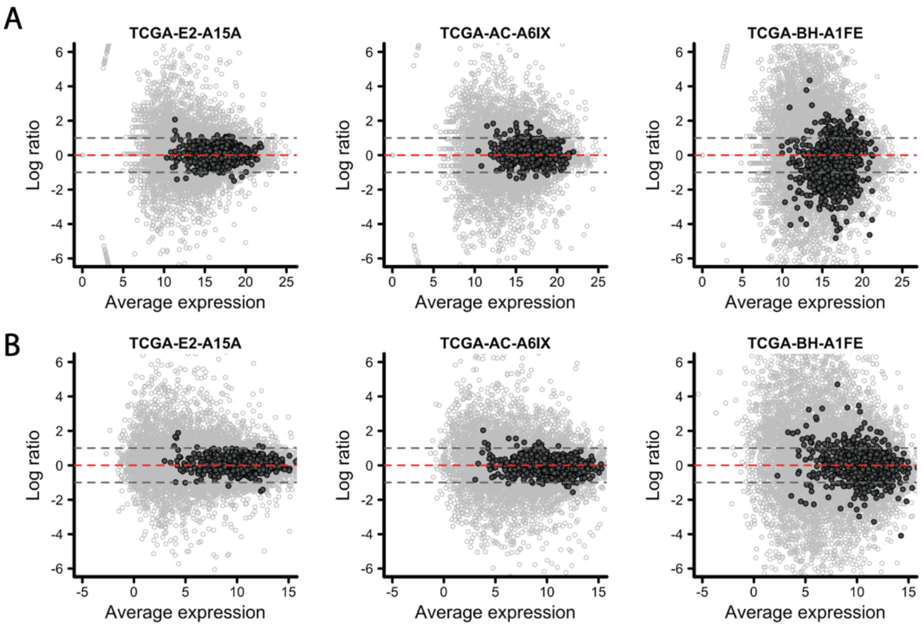


**Supplementary Figure 29. Impact of flow cell chemistries on gene expression differences between paired primary and metastatic samples in the TCGA BRCA RNA-sequencing data**. MA plot of three paired primary and metastatic samples profiled within (the first two plots of top row) and across flow cell chemistries (third plot of top row) in the TCGA FPKM.UQ data. Second row: same a first row, for the RUV-III normalized data. The solid black points in the MA plots represent the genes highly affected by flow cell chemistries.

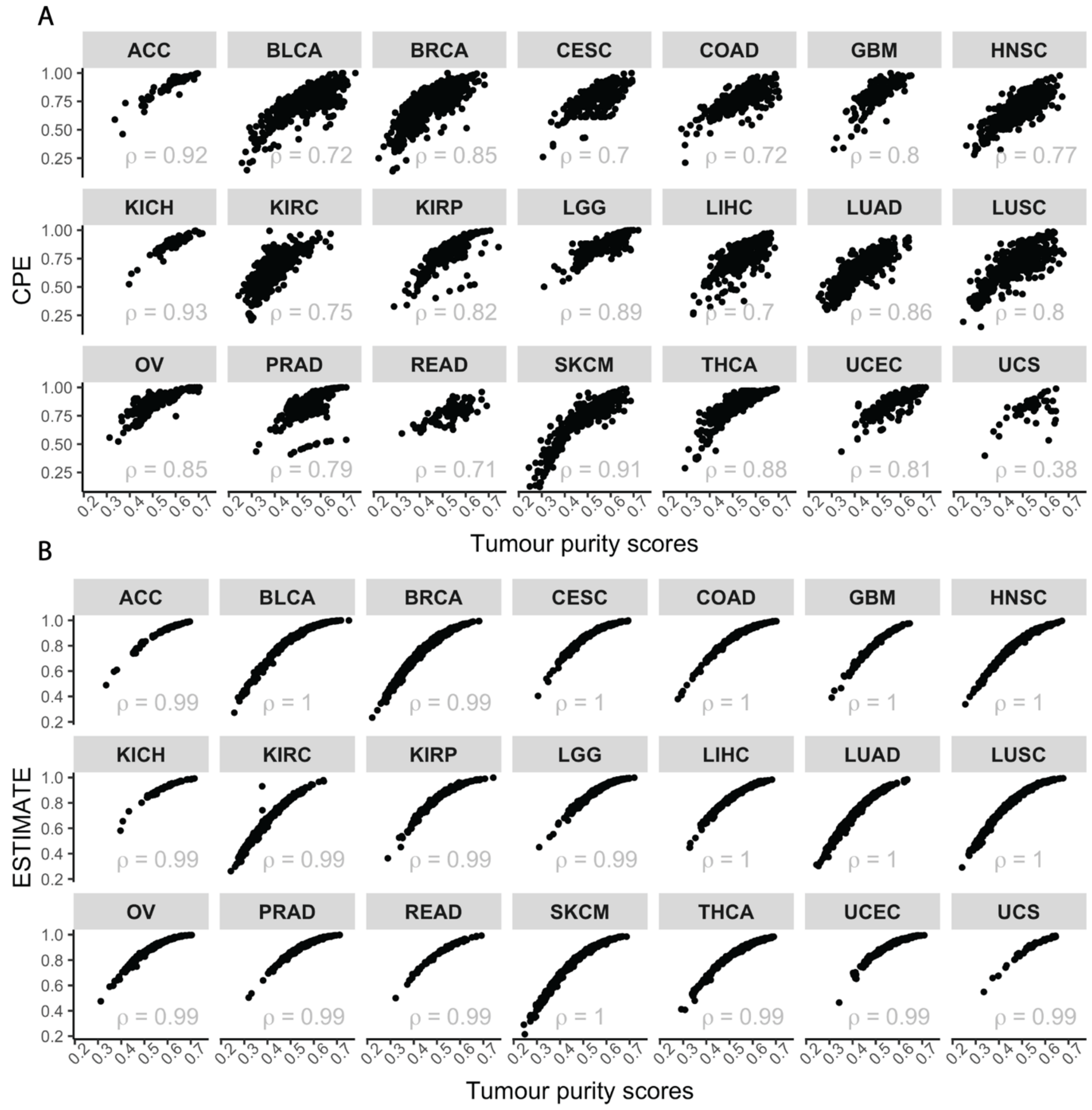


**Supplementary Figure 30. Tumour purity estimates in TCGA RNA-sequencing datasets. A)** The scatter plots show the relationship between CPE and tumour purity scores. **B)** The scatter plots show relationship between the tumour purity calculated by the ESTIMATE method and consensus measurement of purity estimations (CPE) approach in the TCGA RNA-seq datasets.

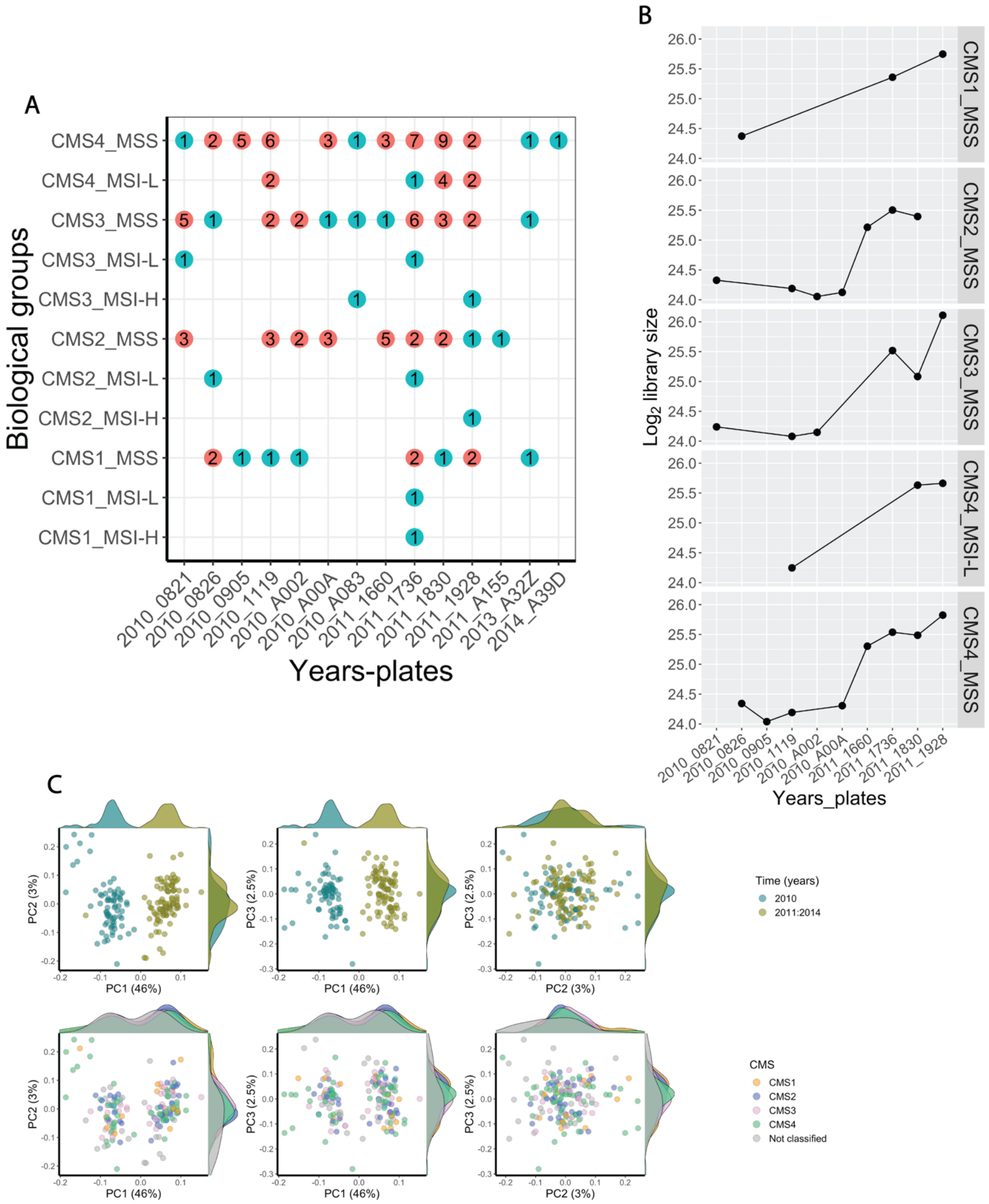


**Supplementary Figure 31. PRPS for RUV-III normalization of the TCGA READ RNA-sequencing data. A)** Plot showing the sample sizes of the major biological groups across plates in the TCGA READ RNA-seq data. **B)** Library sizes of pseudo-samples. **C)** First row: scatter plots show the first three principal components coloured by key time intervals of only 919 negative control genes in the TCGA raw counts RNA-seq data. Second row, same as first row, coloured by CMS.

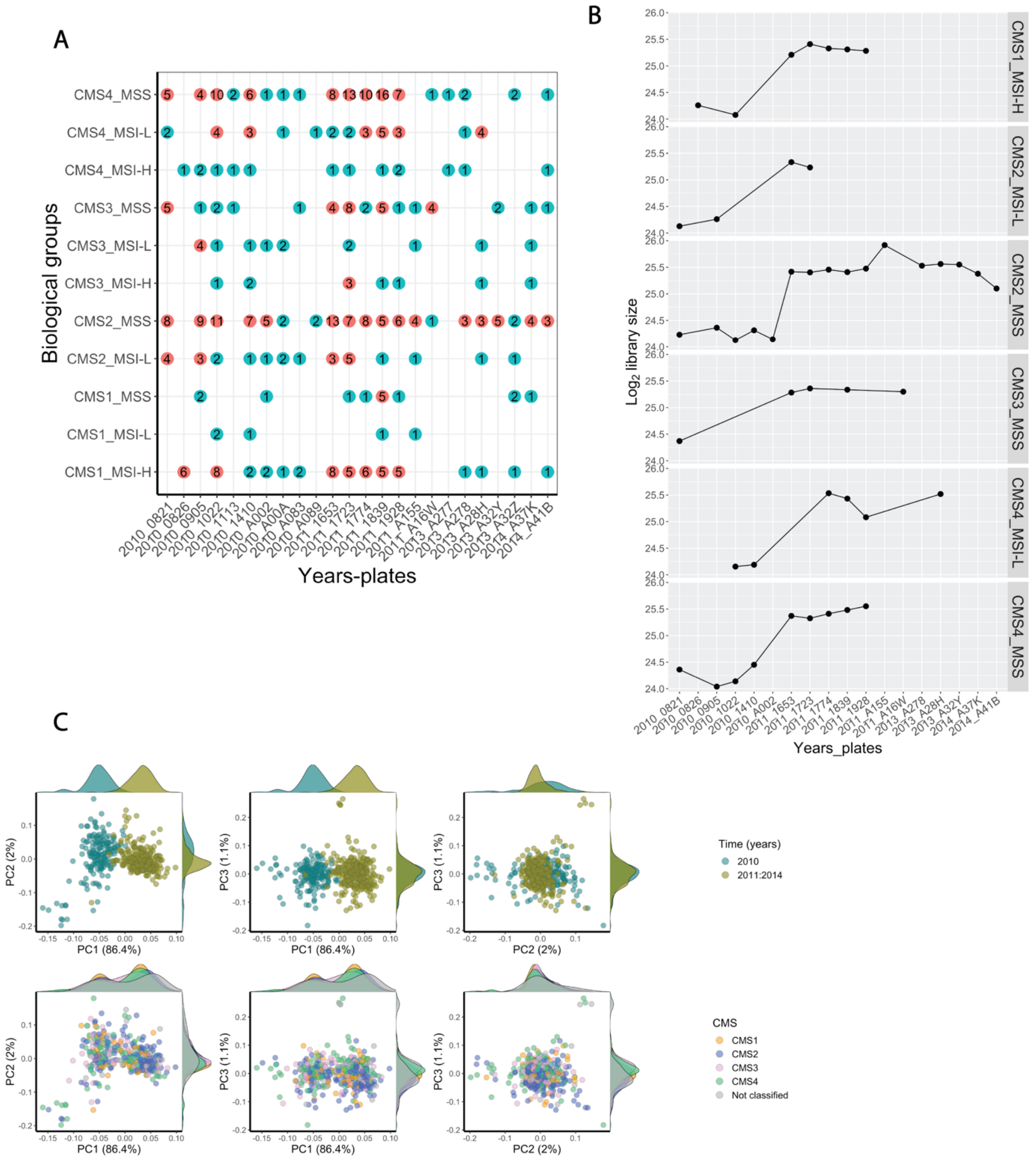


**Supplementary Figure 32. PRPS for RUV-III normalization of the TCGA COAD RNA-sequencing data.** **A)** Plot showing the sample sizes of the major biological groups across plates in the TCGA COAD RNA-seq data. **B)** Library sizes of pseudo-samples**. C)** First row: scatter plots show the first three principal components coloured by key time intervals of only 262 negative control genes in the TCGA raw counts RNA-seq data. Second row, same as first row, coloured by CMS.

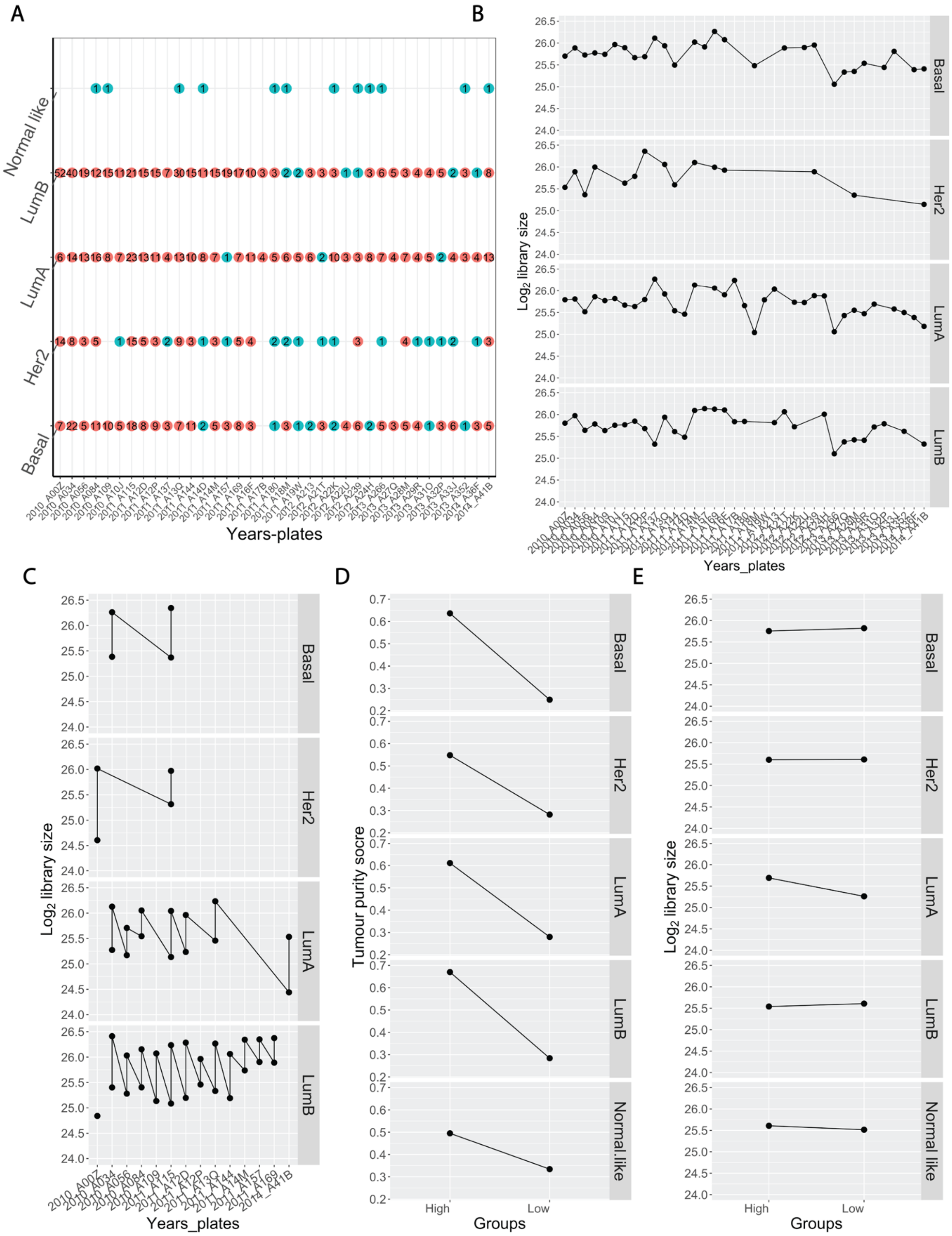


**Supplementary Figure 33. PRPS for RUV-III normalization of the TCGA BRCA RNA-sequencing data. A)** Plot showing the sample sizes of the major biological groups across plates in the TCGA BRCA RNA-seq data. **B)** Library sizes of pseudo-samples created for removing plate effects. **C)** Library sizes of pseudo-samples created for removing plate library sizes. **D)** Tumour purity score of pseudo-samples created for removing tumour purity variation. **E)** Library sizes of pseudo-samples created for removing tumour purity variation.

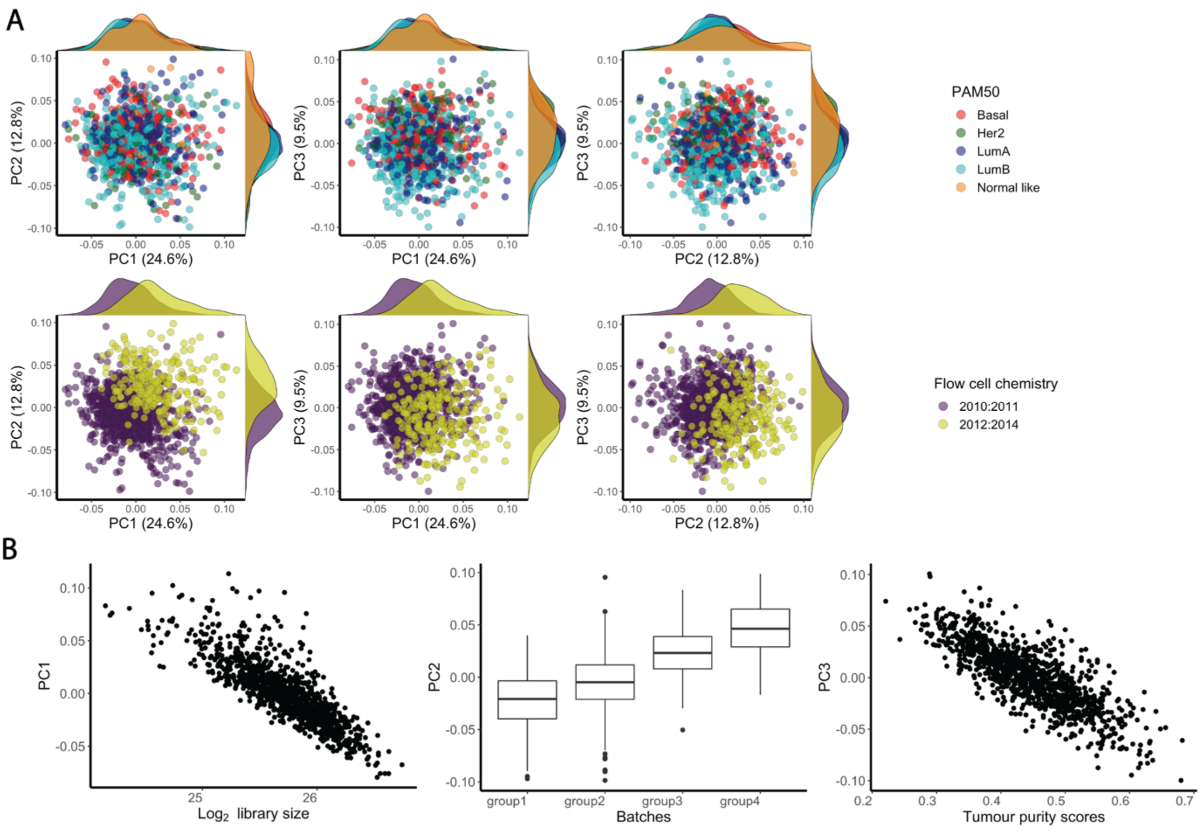


**Supplementary Figure 34. Negative control genes for RUV-III normalization of the TCGA BRCA RNA-sequencing data. A) first row:** the first three PC coloured by PAM50 subtypes of the TCGA BRCA raw counts using a set of negative control genes. Second row: same a first row, coloured by flow cell chemistry. **B)** Scatter plots show relationship between the first three PC of the negative control genes with library size, batches and tumour purity scores.
